# Genetic analysis of a phenotypic loss in the mechanosensory entrainment of a circalunar clock

**DOI:** 10.1101/2022.10.12.511720

**Authors:** Dušica Briševac, Celine Prakash, Tobias S. Kaiser

## Abstract

Genetic variants underlying traits that become either non-adaptive or selectively neutral are expected to have altered evolutionary trajectories. Uncovering genetic signatures associated with phenotypic loss presents the opportunity to discover the molecular basis for the phenotype in populations where it persists. Here we study circalunar clocks in populations of marine midge *Clunio marinus*. The circalunar clock synchronizes development to the lunar phase, and it is set by moonlight and tidal cycles of mechanical agitation. Two out of ten studied populations have lost their sensitivity to mechanical agitation while preserving sensitivity to moonlight. Intriguingly, the F1 offspring of the two insensitive populations regained the sensitivity to mechanical entrainment, implying a genetically independent loss of the phenotype. By combining quantitative trait locus mapping and genome-wide screens, we explored the genetics of this phenotypic loss. QTL analysis suggested an oligogenic origin with one prevalent additive locus in one of the strains. In addition, it confirmed a distinct genetic architecture in the two insensitive populations. Genomic screens further uncovered several candidate genes underlying QTL regions. The strongest signal under the most prominent QTL contains a duplicated *STAT1* gene, which has a well-established role in development, and *CG022363*, an ortholog of the *Drosophila melanogaster CG32100* gene, which plays a role in gravitaxis. Our results support the notion that adaptive phenotypes have a complex genetic basis with mutations occurring at several loci. By dissecting the most prevalent signals, we started to reveal the molecular machinery responsible for the entrainment of the circalunar clock.

## INTRODUCTION

Life on earth adapted to anticipate predictable changes in its environment in order to survive, a case in point is the ubiquity of biological clocks. Due to the earth’s rotation around its axis, most living creatures are exposed to 24-hour cycles, which has resulted in the pervasiveness of circadian clocks (Pittendrigh 1960; Dunlap and Loros 2017). Furthermore, marine organisms inhabiting intertidal zones are exposed to tidal cycles of 12.4 hours, which are modulated across the 29.53-day lunar cycle. Thus, marine organisms have evolved circatidal and circalunar clocks. Due to their universal occurrence, circadian clocks have been intensely studied over the last century (Wager-Smith and Kay 2000; Takahashi 2017). Comparatively much less is known about circatidal and circalunar clocks (Kaiser and Neumann 2021; Goto and Takekata 2015; Andreatta and Tessmar-Raible 2020; Raible et al. 2017), although some argue that as life evolved in the marine environment circadian clocks may have evolved from evolutionarily older circatidal or circalunar clocks (Wilcockson and Zhang 2008; Naylor 2010).

Biological clocks must be appropriately set to fulfill their role in synchronizing endogenous physiological processes, reproduction, and behavior to the exogenous environmental cycles. Environmental variables that reliably fluctuate with geophysical cycles serve as clock synchronizers, so-called zeitgebers. The most studied zeitgeber is the light-dark cycle that synchronizes the circadian clock (Pittendrigh 1960). Two other synchronizers of the circadian clock that were experimentally confirmed are temperature and vibration (López-Olmeda et al. 2006; Simoni et al. 2014; Caldart et al. 2020; Liu et al. 1998). In contrast, many environmental variables fluctuate with the tides and the following have been shown to serve as strong zeitgebers of the tidal clocks: mechanical disturbance of the water (Enright 1965; Jones and Naylor 1970; Hastings 1981), changes in hydrostatic pressure (Jones and Naylor 1970; Gibson 1971; Northcott 1991), temperature fluctuations (Williams and Naylor 1969; Holmstrom and Morgan 1983), changes in salinity (Taylor and Naylor 1977), immersion and emersion (Williams and Naylor 1969).

Not surprisingly, moonlight was shown to be a unique cue for synchronizing lunar clocks (Hauenschild 1960; Bunning and Müller 1961; Neumann 1966; Saigusa 1980; Franke 1985). Furthermore, several synchronizers that were first discovered as tidal cues, were consequently demonstrated to be strong zeitgebers for setting circalunar clocks: vibration that accompanies the rise and fall of the tides (Reid and Naylor 1985; Neumann 1978) and temperature fluctuations (Neumann and Heumbach 1984). Depending on the stability and robustness of the cycles in the environment that the organism inhabits, different zeitgebers provide reliable cues for biological clocks in different organisms. Finally, while biological clocks are not crucial for the survival of all organisms, the harsher the environmental cycles, the stronger the selection pressure on the presence of reliable biological clocks. Studying organisms inhabiting these harsh environments promises to give insight into the nature of yet unexplored biological clocks. One such species whose survival critically depends on its ability to simultaneously synchronize to lunar and circadian cycles is the marine midge *Clunio marinus*.

*Clunio* spends most of its life in a larval stage submerged in the intertidal zone of the Atlantic Ocean. During full moon and new moon, adults emerge on the sea surface, mate, oviposit eggs and die within a few hours. Circadian and circalunar clocks allow them to precisely time reproduction to the lowest of the low tides. Individuals that do not emerge at the appropriate time miss the ecologically suitable low tide for reproduction and the opportunity to mate and are thus eliminated from the population. Therefore, strong selection pressure shapes various timing phenotypes in populations that encounter different tidal regimes along the Atlantic coast (Neumann 1967; Kaiser 2014; Kaiser et al. 2021, 2010, 2011). Moonlight, tidal turbulence and temperature have been shown to be zeitgebers setting the circalunar clock of *Clunio marinus* (Neumann 1966; Neumann and Heimbach 1978; Neumann and Heumbach 1984; Neumann 1978). However, different *Clunio* populations are differentially sensitive to zeitgebers, most likely due to the unreliability of different zeitgebers in certain geographical locations (Neumann and Heimbach 1978). Neumann discovered one population insensitive to moonlight and two that were insensitive to tidal turbulence (Neumann and Heimbach 1978). Tidal turbulence was defined as low frequency, low amplitude vibration that coincides with the rising tide (Neumann and Heimbach 1978). This stimulus shifts every day by 50 minutes resulting in a semi-lunar 14.7 days entrainment pattern (Neumann and Heimbach 1978).

Evolutionary losses of function can have a creative role in evolution (Albalat and Cañestro 2016), and genetic and genomic analysis of the affected populations can identify the genes involved in corresponding molecular pathways (Monroe et al. 2021). Our goal was to establish if the loss of mechanosensory entrainment in the two populations was consistent with it having a common genetic basis, or whether it occurred independently in each population. We also sought to determine if genetic control of this phenotype is likely controlled by a single locus of major effect or whether multiple loci play discernible roles. Finally, we aimed to identify genes likely to be responsible for impacting the trait.

## RESULTS

### Loss of sensitivity to mechanical entrainment is a genetically determined trait that evolved independently in two *Clunio* populations

Circalunar clock robustly regulates the emergence of *Clunio* adults over a lunar month. We study this phenomenon under laboratory conditions by counting the number of emerged adults per day over several lunar cycles, and then assess characteristics of the phenotype using circular statistics: phase, period, rhythmicity, etc. The sensitivity of different strains to the zeitgebers is therefore estimated indirectly via the strength of their emergence rhythms upon entrainment to moonlight or tidal turbulence. Here we tested the entrainment of Plou-2NM, Ros-2NM, Lou-2NM, Bria-1SL, and Por-1SL under tidal turbulence for the first time, while the entrainment to moonlight (Kaiser 2014; Kaiser et al. 2016; Kaiser et al. 2021) and tidal turbulence (Neumann and Heimbach 1978) were previously reported for the other populations (Figure 1 A, Supplemental Figure 1, Supplemental Tables 1 and 2).

**Figure 1.**
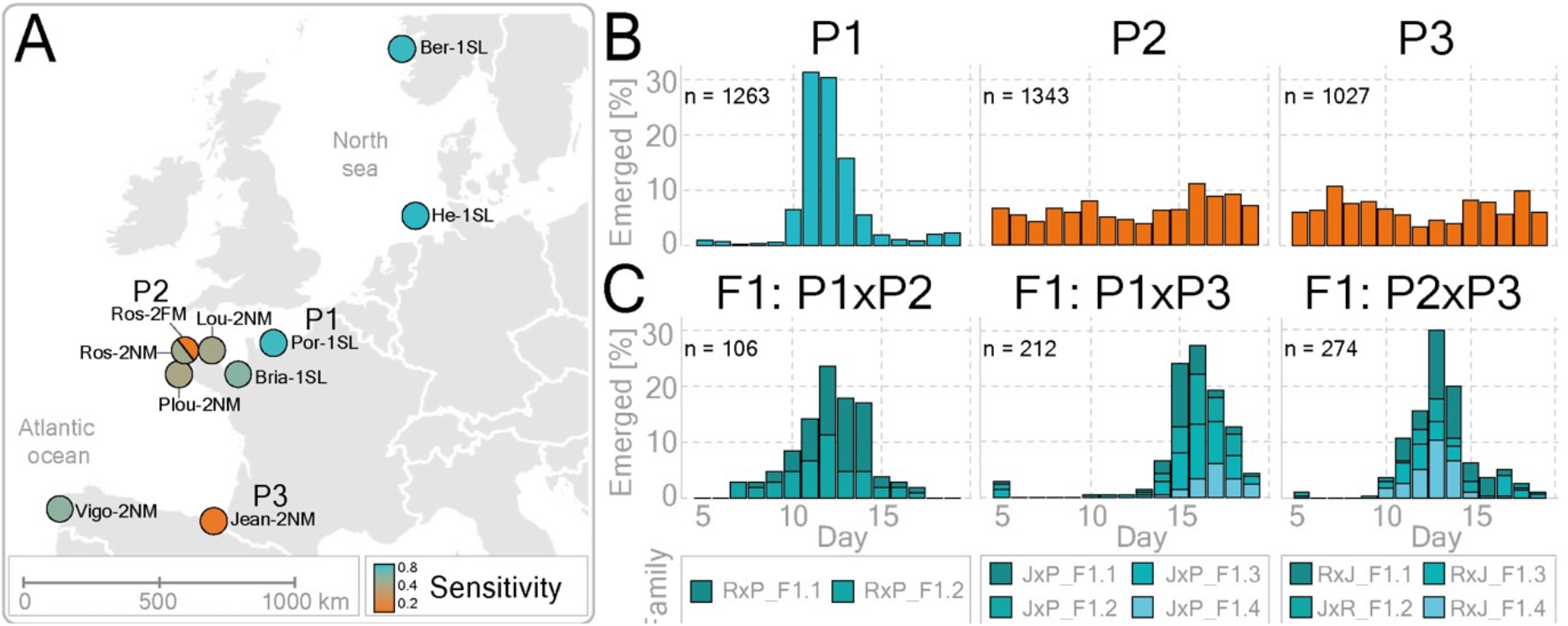
Sensitivity to mechanical entrainment was lost twice independently in European *Clunio* populations. (A) The origin of the ten *Clunio* populations used in this study is shown on the map (Supplemental Figure 1). Heatmap depicts the sensitivity of each strain to mechanical entrainment as estimated by the circular statistics (Supplemental Table 2). (B, C) Graphs show the fraction of emerged individuals per lunar day upon mechanosensory entrainment. The total number of emerged individuals is depicted in the left corner of each bar graph. (B) Graphs depict the emergence patterns of the parental populations: P1 (Por-1SL), P2 (Ros-2FM), and P3 (Jean-2NM). P2 and P3 populations are insensitive to tidal turbulence as shown by the arhythmic emergence patterns, while the P1 population is sensitive. Geographical locations and the years when strains were established are given in Supplemental Table 1. (C) Crossing sensitive (P1) and insensitive (P2 or P3) strains resulted in the sensitive F1 progeny (left and middle). Furthermore, when the two insensitive strains were crossed, the resulting F1 hybrids regained sensitivity to the entrainment (right). The total number of individuals per generation is listed in Supplemental Table 3.

Vigo-2NM is the most southern strain and it is sensitive to tidal turbulence. Going north, we come across Jean-2NM which is insensitive to tidal turbulence, followed by five closely related populations at the coast of Bretagne: Plou-2NM, Ros-2FM, Ros-2NM, Lou-2NM, Bria-1SL, Por-1SL; and finally, the two most northern populations: He-1SL in Germany and Ber-1SL in Norway (Figure 1 A). Bretagne populations vary from very sensitive in the north (Por-1SL and Bria-1SL), and less sensitive in the south (Ros-2NM, Lou-2NM, and Plou-2NM) to completely insensitive (Ros-2FM) (Figure 1 A, Supplemental Figure 1, Supplemental Table 2). This suggests that the frequency of the “insensitive allele” may vary among the Bretagne populations, giving rise to varying degrees of sensitivity. Furthermore, as Ros-2FM and Jean-2NM are arrhythmic under tidal turbulence but rhythmic under moonlight (Figure 1 B and Supplemental Figure 1 O, S) (Neumann and Heimbach 1978), we can conclude that their lunar clocks are intact, but sensory inputs have evolved rendering them insensitive to one of the cues. To characterize the genetic basis for this phenotypic loss, we crossed turbulence-insensitive strains to a strain sensitive to both tidal turbulence and moonlight, Por-1SL (Figure 1 B, Supplemental Figure 1 F, G), and analyzed the emergence of adults in F1 and F2 generations (Figure 1 B, C and Supplemental Figure 2). We used vector length of the summary circular statistics for estimating the strength of the entrainment and found that sensitivity to tidal turbulence is genetically determined and a dominant trait (Supplemental Table 2).

To test if the same mutations are responsible for the loss of sensitivity in Jean-2NM and Ros-2FM we performed a complementation cross. Interestingly, the four F1 families raised separately all regained their sensitivity to mechanical entrainment (Figure 1 C). This finding strongly suggests a different and recessive genetic basis for the loss of trait in Jean-2NM and Ros-2FM.

### Discovering genomic loci responsible for the phenotypic loss in the Ros-2FM population

Quantitative trait loci (QTL) mapping was conducted to locate the regions of the genome containing genetic variants responsible for the loss of sensitivity to tidal turbulence in the Ros-2FM population. The resolution of QTL mapping depends on the number and distribution of markers as well as the recombination events which in turn depends on the number of individuals in the crossing family. To maximize our chances of achieving narrow confidence intervals, we performed a large number of crosses and then selected two families for the analysis: F2 progeny of Ros-2FMxPor-1SL cross (RxP-F2.1) and a backcross progeny of Ros-2FMxPor-1SL F1 female to Ros-2FM male (RxP-BC.1) (Supplemental Table 3). The number of informative markers was 137 in RxP-F2.1 and 123 in RxP-BC.1. The total number of recombination events was 269 and 61, while the number of unique genomic positions of the recombination events was 51 and 36 in RxP-F2.1 and RxP-BC.1 families respectively (Figure 2 B and D).

**Figure 2.**
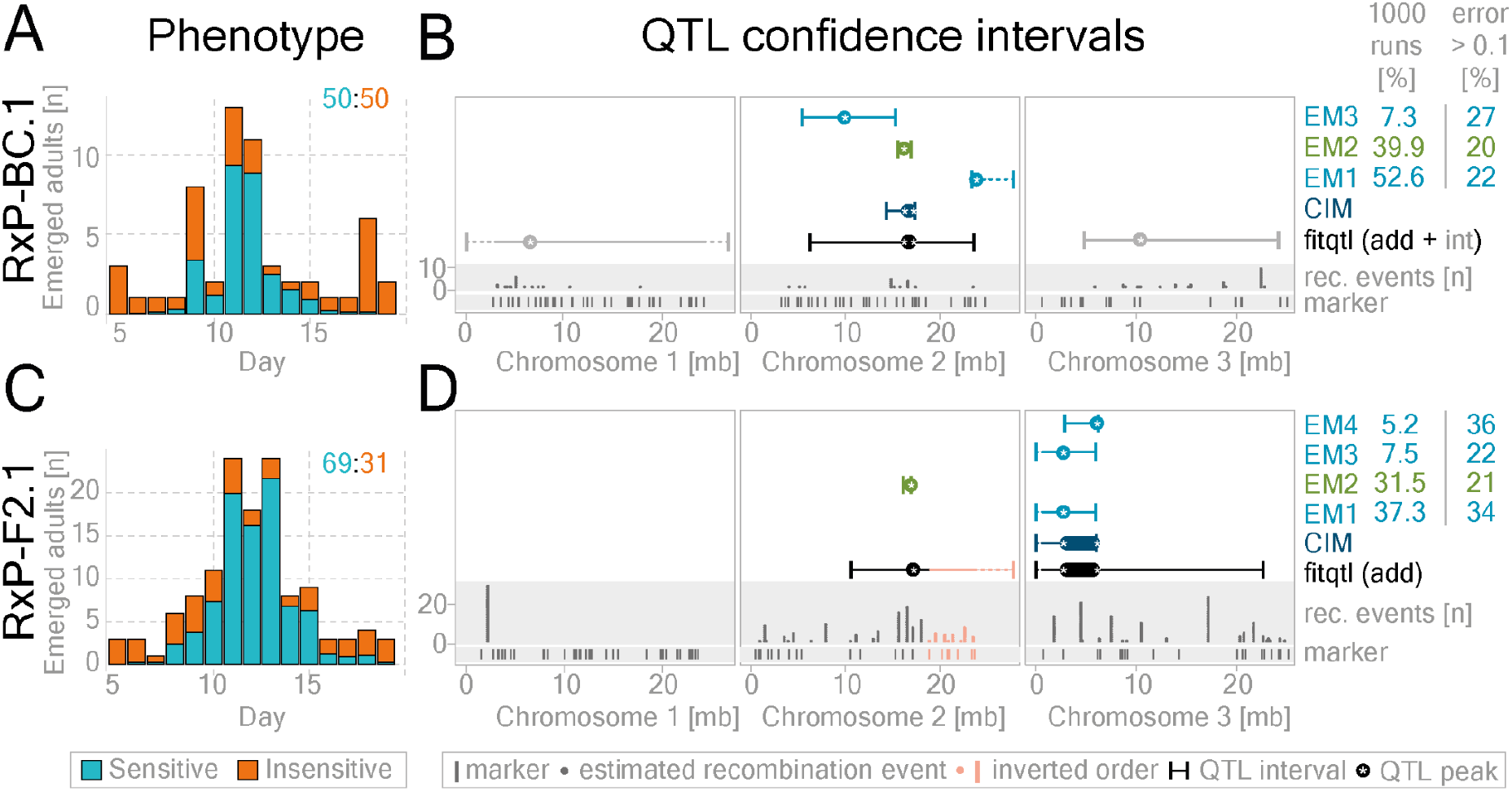
QTL mapping in two Ros-2FMxPor-1SL mapping families reveals one shared additive QTL on the second chromosome. Regions of the genome harboring genes responsible for the loss of mechanical entrainment in the Ros-2FM population were identified in a [Ros-2FM x Por-1SL] x Ros-2FM backcross (RxP-BC.1) see panels A and B, and a [Por-1SL x Ros-2FM] F2 intercross (RxP-F2.1) see panels C and D. Bar graphs show the number of emerged individuals per day (A, C). The proportion of insensitive (orange) and sensitive (blue) individuals found on each day was calculated based on estimated probabilities (Supplemental Figures 3 and 4). The ratio of sensitive and insensitive individuals in each family is indicated in the top right corner. (B, D) QTL confidence intervals are given for: composite interval mapping – dark blue, fitqtl: additive loci – black, fitqtl: epistatic loci – gray, EM-algorithm – light blue. The green marks the phenotypic panel with the highest convergence in EM analysis (i.e. the number of times the panel was found to be the best in 1000 runs) and the lowest error (i.e. the fraction of individuals in each panel for which the binary phenotype differs significantly from the starting probabilities; see methods QTL mapping/EM-pipeline, Supplemental Table 4).

If ~130 markers and ~40 unique recombination events would be evenly distributed along the 80Mb genome, we could achieve the mapping resolution of ~2-3Mb. However, several non-recombining regions were found in both families and one in which the marker order was inverted as compared to the reference (Figure2 B and D). These regions are thought to be large polymorphic inversions (Michailova 1980) which are limiting mapping resolution on the first chromosome and in the right arm of the second chromosome (manuscript submitted, Briševac et al. 2022).

In order to phenotype F2 and BC progenies, we must distinguish between “sensitive” individuals that emerged within the Por-1SL-like peak and “insensitive” individuals that can emerge on any lunar day. However, the emergence peak does not only contain sensitive individuals, but also some of the insensitive individuals. To overcome this issue, we tested different phenotyping strategies and mapping algorithms (Supplemental Figures 3-8) (see methods QTL section for more details). We calculated the probability of finding sensitive and insensitive individuals on each lunar day (Supplemental Figures 3 and 4) and used it as a phenotypic score for the QTL analysis. In addition, we generated a reduced dataset by excluding the individuals with uncertain phenotypes and treated those with the probability of being “insensitive” higher than 0.7 as “insensitive” and lower than 0.3 as “sensitive” (Supplemental Figure 3). Finally, this approach allowed us to estimate relatively precisely the ratio of the two phenotypes in the F2 and BC generations: 69:31 in the RxP-F2.1 intercross (Figure 2 C) and 50:50 in the RxP-BC.1 backcross (Figure 2 A). The difference in ratios is attributed to a higher portion of sensitive individuals (parental and F1 genotypes) in an F1xF1 intercross as compared to an F1xRos-2FM backcross. Similar ratios were found in Jean-2NMxPor-1SL intercross families (see below). Such segregation of parental phenotypes in F2 and BC progenies indicates that this trait is determined by a small number of loci.

Furthermore, in order to screen for additive QTLs, we ran standard interval mapping with *scanone* (Supplemental Figure 6 E-H) and composite interval mapping (Figure 2 B and D, Supplemental Figure 6 I-L). To investigate QTLs in epistasis we ran a two-dimensional scan with *scantwo* function (Supplemental Figure 7). QTLs identified with *scanone* and *scantwo* were then fed into the multiple-QTL-mapping pipeline implemented in the R/qtl package with the *fitqtl* function (Figure 2 B and D, Supplemental Figure 6 Q-T and 7). Since various models can be significant with *fitqtl*, we also tested a Bayesian method implemented in R package *qtlbim* designed to find the best QTL model for *fitqtl* (Supplemental Figure 6 Y-AB).

Multiple QTL mapping pipeline revealed one additive QTL and two QTLs in epistasis in the RxP-BC.1 family, and two additive QTLs in the RxP-F2.1 family (Figure 2 B and D, Supplemental Figures 6-8). The additive QTL on the second chromosome was found in both crossing families. The QTL on the third chromosome interacts additively with the QTL on the second chromosome in the RxP-F2.1 reduced dataset (Supplemental Figure 7 E-F), while in the RxP-BC. 1 family it is in a negative additive-by-additive epistatic interaction (Cockerham 1954) with the QTL on the first chromosome: the QTL on the first chromosome has a positive additive effect in the heterozygous AB background of the QTL on the third chromosome and vice-versa (Figure 2 D, Supplemental Figure 7 A-D). The QTLs in epistasis were found only in one of the families, potentially because the presence of the epistatic interaction depends on the genetic background. This can occur if the mutations underlying QTLs in epistasis are not fixed in the two populations. In other words, if a mutation underlying QTL1 only has an effect in the presence of another mutation underlying QTL2, and one of the two alleles is absent in the parent of that crossing family, the epistatic interaction would not be identified. Thus, to find the regions of the genome containing the loci most likely pervasive in the natural populations, we further focused only on the additive QTLs.

In order to further estimate the effect of the phenotyping uncertainty on additive QTLs, we generated the *scanone*-optimized expectation-maximization pipeline (see methods for more details, Figure2, and Supplemental Figures 3-8). In a nutshell, all individuals are assigned binary phenotypes (0 or 1) depending on their starting phenotype possibilities. Then the algorithm changes the phenotypes of individuals in order to find the binary phenotype panel with the highest LOD score (Figure 2 C and D, Supplemental Figure 6 M-P, for more details see methods). The resulting binary phenotype panels are assessed for their credibility by how often the algorithm converges to a specific panel (% convergence) and by which fraction of animals differs by more than 0.1 to the starting probability. In the RxP-BC.1 family, the QTL interval of the EM binary panel with the lowest percentage of individuals with error>0.1 (20%) and the second-highest percentage of convergence (39.9%) perfectly overlaps with the QTL interval provided by CIM, and *fitqtl* mapping on probability phenotypes (Figure 2 B, Supplemental Figure 8 A-B). In the RxP-F2.1 family, the best panel according to the same criteria is also on the second chromosome: the percentage of convergence is 31.5, percentage of individuals with error > 0.1 is 21% (Figure 2 D and K, Supplemental Figure 8 C-D). The high level of convergence in the phenotype panels shows that the QTL landscape does not contain too many potential local optima. This suggests that the phenotyping uncertainty is limited.

Other than the phenotyping uncertainty, a polygenic or oligogenic origin could also lead to a reduction in the QTL LOD scores. In the RxP-BC.1 family, the full *fitqtl* model explains 28% of the phenotypic variance (Supplemental Figures 6-8 and Supplemental Tables 4). In the reduced (binary) dataset that number increases to 40.79% (Supplemental Figures 6-8 and Supplemental Tables 4). The QTL on the second chromosome alone explains 9.27% and the epistatic interaction between the first and the third chromosome explains 11.09%; while in the reduced dataset that becomes 2.82% and 11.68% respectively. In the RxP-F2.1 family, the *fitqtl* model explains 13.74% in the full and 21.8% in the reduced dataset. The percentage of variance explained by the QTL on the second chromosome is 5.9% and the third 8% (18.29% and 16.56% respectively in the reduced dataset). Previous QTL analysis on mapping the lunar phase, a phenotype of discrete nature, identified two QTLs: one explains 23% of the variation and the other 14% (Kaiser and Heckel 2012). In both the present and the previous study, we find a small number of significant loci impacting a trait, with a comparable proportion of phenotypic variance explained. This potentially indicates that we did not lose significant mapping power due to the non-discrete nature of the phenotype in the current study. Furthermore, this implies that if a small number of loci collectively accounts for up to 20-40% of the phenotypic variance, the unidentified loci or loci of small effect size may still play a substantial role.

Finally, although the multiple QTL mapping is crucial for investigating the most likely number of QTLs, it tends to overestimate the QTL confidence intervals. Thus, to investigate the genes underlying the additive QTLs we relied on composite interval mapping conducted in 10cM windows (dark blue in Figures 2 B and D and Figure 3 A) while being informed by the QTL intervals resulting from the best binary phenotypic panels found by the EM algorithm (light green in Figure 2 B and D and Figure 3A).

**Figure 3.**
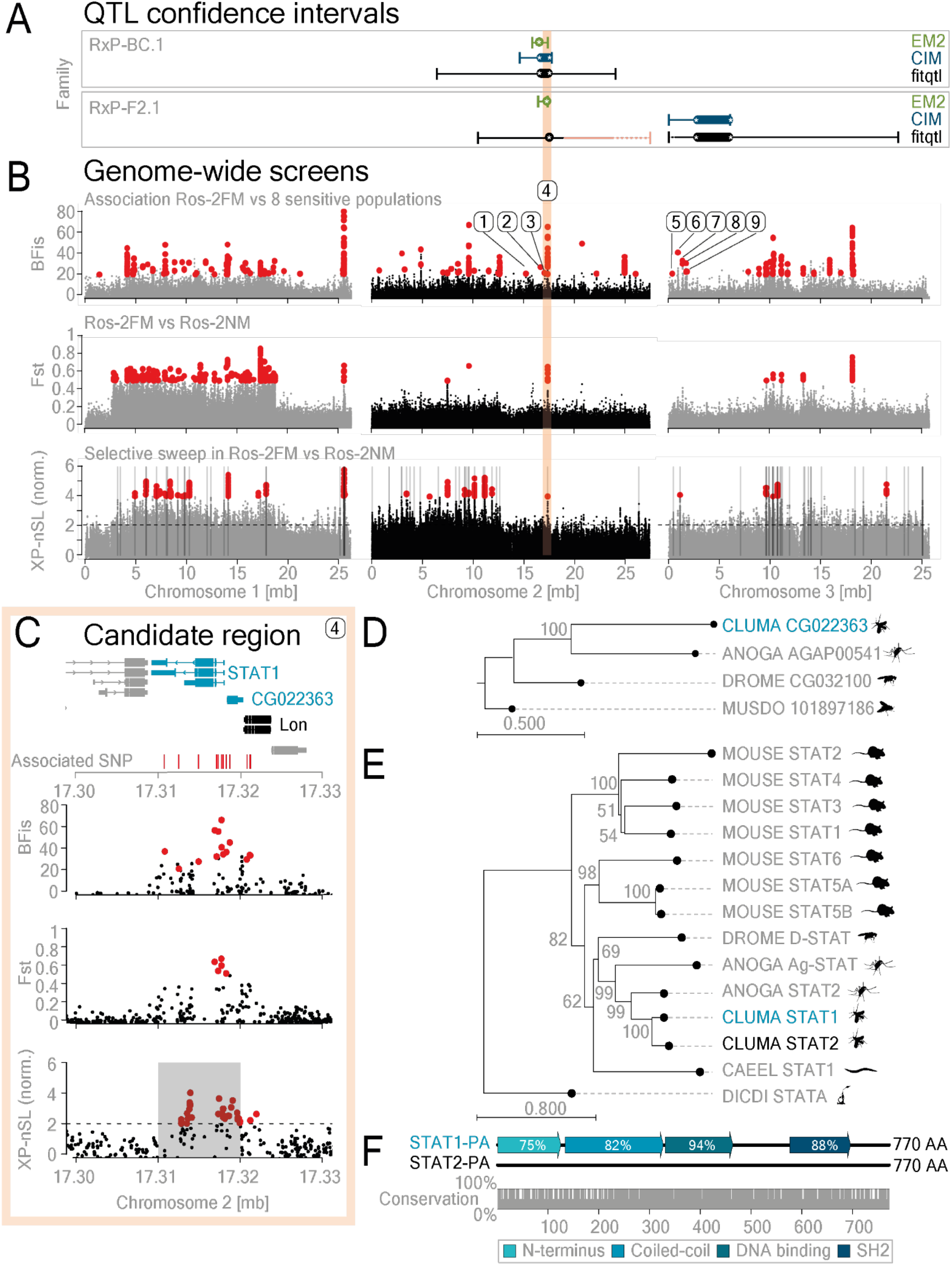
A combination of QTL mapping and genome-wide screens points to *STAT-1* and gravitaxis gene *CG022363* as likely contributing to the loss of sensitivity to mechanical entrainment in the Ros-2FM population. (A) QTL confidence intervals from the two mapping families are plotted along the three chromosomes (modified from Figures 2 B and D). (B-top). The 746.887 variants called from Ros-2FM and eight differentially sensitive populations were screened for their association with the sensitivity to mechanical entrainment using BayPass. Sensitivity to mechanical entrainment was estimated from emergence patterns using circular statistics (Supplemental Figure 10 A, Supplemental Table 3). Median vector length was used as the phenotypic score. The Bayesian factor (BFis) depicting the strength of the association is plotted for all the variants along the three chromosomes (gray and black). 375 significantly different variants, as determined by BFis >= 20, eBPis >= 2, XtXst >= 21.67 are marked in red (see Methods section for details). (B-middle and bottom plots): To expose potentially adaptive genetic variants under positive selection as a result of local adaptation, we contrasted the turbulence-insensitive Ros-2FM population with the sympatric Ros-2NM population, determined to be sensitive to this entrainment (Supplemental Figures 1 and 10A). (B-middle). Genetic differentiation (F_ST_) was plotted for the variants found in Ros-2FM and Ros-2NM populations. Red marks F_ST_ values above 0.5. (B-bottom). The cross-population nSL (number of segregating sites by length) statistic shows the decay of haplotype homozygosity surrounding adaptive alleles as a result of a selective sweep in Ros-2FM in contrast to Ros-2NM. The top 1% clusters of at least 10 variants with XP-nSL values above 2 in a 10kb window were called significant (highlighted in gray). (C) The region of the genome under the prevalent additive QTL on the second chromosome that also contains the most associated variants with the highest association scores, strong F_ST_ signal, and a significant signature of a partial sweep, harbors the *STAT-1* and *CG022363* genes (shown in red shaded area). The position of the associated variants is shown in red, the candidate gene in blue, and two neighboring genes in gray. For the depiction of all genes affected by associated mutations under the two QTLs see Supplemental Figures 10-12. (D, E) The phylogenetic trees of the *STAT* and *CG022363* gene families were shown for *Caenorhabditis elegans, Drosophila melanogaster, Musca domestica, Anopheles gambiae, Mus musculus, Clunio marinus* candidate gene (blue), and *Clunio marinus* ortholog of the candidate gene (black). (F) *C. marinus* has two STAT genes: CG012971 (STAT-1) and CG022905 (STAT-2). The alignment, conserved domains, and percentage of conservation between the two amino acid sequences are shown.

### Whole-genome sequencing reveals genetic variants associated with insensitivity to tidal turbulence in Ros-2FM

As discussed above, the resolution of the QTL mapping in our model system can theoretically go down to 2-3Mb which can still harbor several hundred genes. Therefore, in order to further identify specific genomic loci underlying QTL regions, we combined QTL mapping with genome-wide association analysis (Figure 3 A and B). We sequenced 20-24 field-caught males from nine *Clunio* populations differentially sensitive to tidal turbulence (Supplemental Tables 1, 2, and 5, Figure 1A, Supplemental Figure 1) and called 746,887 SNPs and small indels. We used circular summary statistics (vector length) as the population-wide phenotypic score for sensitivity to tidal turbulence (Figure 3 B, Supplemental Table 2). We then applied the bayesian tool BayPass for calculating the strength of association of each of the variants to the insensitivity score while using a kinship matrix to correct for population structure (Supplemental Figure 9). Out of 746,887 variants, 357 were significantly associated with sensitivity to tidal turbulence. These variants affect 178 genes, as determined by SNPeff (Supplemental Table 6, Supplemental Figures 9-10). Most of the variants are located in non-coding regions and only a handful have potentially disruptive impacts on the neighboring genes (Supplemental Figure 9 C-D).

Crucially, the Ros-2NM population, which is sympatric to the insensitive Ros-2FM population, is sensitive to tidal turbulence (Supplemental Figure 1 and Supplemental Table 2). We can therefore ask if this phenotypic loss occurred as a result of a recent selective sweep due to local adaptation. To explore this, we first estimated genomic differentiation (F_ST_) between the two populations and found that the most prominent loci identified in the BayPass screen also have high F_ST_ values (Figure 3 B). Furthermore, if we assume that the causal genetic variant underwent positive selection as a result of a selective sweep, we can expect that it would leave a characteristic pattern of long high-frequency haplotypes and low genetic diversity in its vicinity (Szpiech and Hernandez 2016). The selective sweep occurring as a result of a local adaptation is calculated as the decay of haplotype homozygosity between the two populations (Szpiech and Hernandez 2016) and is implemented in cross-population statistic XPnSL (nSL: number of segregating sites by length) (Ferrer-Admetlla et al. 2014) in selscan 2.0 (DeGiorgio and Szpiech 2021)(for details see Methods section). The top 1% of the 10kb regions containing a cluster of alleles with high XPnSL values were considered to be candidate regions under selection in Ros-2FM (Figure 3 B).

Finally, we combined the results from QTL analysis, the genome-wide association screen, and the selection screen. We identified the loci underlying additive QTLs (Supplemental Figure 10) and found the orthologues in model organisms of all the genes in the vicinity of the associated mutations (Supplemental Figures 11 and 12). When we zoomed into the genomic region underlying the shared additive QTL on the second chromosome (Figure 3 A) and looked for the variants with the highest association score as shown by BayPass, Fst, and potentially a result of a selective sweep as shown by XP-nSL, (Figure 3 B) we uncovered a cluster of SNPs in one locus containing three genes: signal transducer and activator of transcription 1 (*STAT-1*), *CG022363*, and *Lon* (Figures 3 C). *CG022363* is an ortholog of the *Drosophila melanogaster CG32100* gene (Figures 3 D), which plays a role in gravitaxis (Armstrong et al. 2006) but is otherwise poorly investigated. The STAT protein family is conserved in most vertebrates and invertebrates (Figure 3E). *Clunio*, unlike *Drosophila melanogaster*, has two paralogues: CG012971 (STAT-1) and CG022905 (STAT-2). STAT-1 is most likely the ancestral STAT protein: ortholog of *Anopheles gamibiae* STAT2 and *Mus musculus* STAT5a,5b, and 6; while STAT-2 is newly duplicated in *Clunio* (Figure 3 E). The two *Clunio* STAT proteins are 83% identical in amino-acid sequence (Figure 3 F). The most divergent protein domains in the two *Clunio* STAT proteins are the N-terminal domain, coiled-coil domain, and sh2 domain (Figure 3 F). Lon is a highly conserved protease (Supplemental Figure 11 F) which is crucial for mitochondrial homeostasis (Pinti et al. 2016).

As the QTL mapping explains at most 40% of the phenotypic variance, other loci of smaller effect must exist and are potentially picked up by the association analysis. We, therefore, explored all the genes identified by BayPass and SNPeff by conducting a gene ontology (GO) term enrichment analysis (Supplemental Figure 13). Out of 178 genes, 67 went into the GO analysis as they passed the criteria of having known orthologues, and 51 of those genes drove 78 significant GO terms (Supplemental Figure 13). Interestingly, gravitaxis was one of the highest significant GO terms. This result, together with the previous identification of the gravitaxis gene *CG022905* under the prevalent QTL, prompted us to look more closely into the genes with known roles in gravitaxis (Supplemental Table 7). We found that out of 27 such genes in Drosophila, 6 are on our list of genes potentially associated with the loss of sensitivity to tidal turbulence (Supplemental Table 7).

### Complex genetic basis for the loss of sensitivity to tidal turbulence in Jean-2NM population

As detailed above, complementation crosses between the two insensitive strains identified a separate origin for the insensitivity to tidal turbulence in Jean-2NM and Ros-2FM. To corroborate this finding, we further explored the genetic basis in the Jean-2NM population (Figure 4).

**Figure 4.**
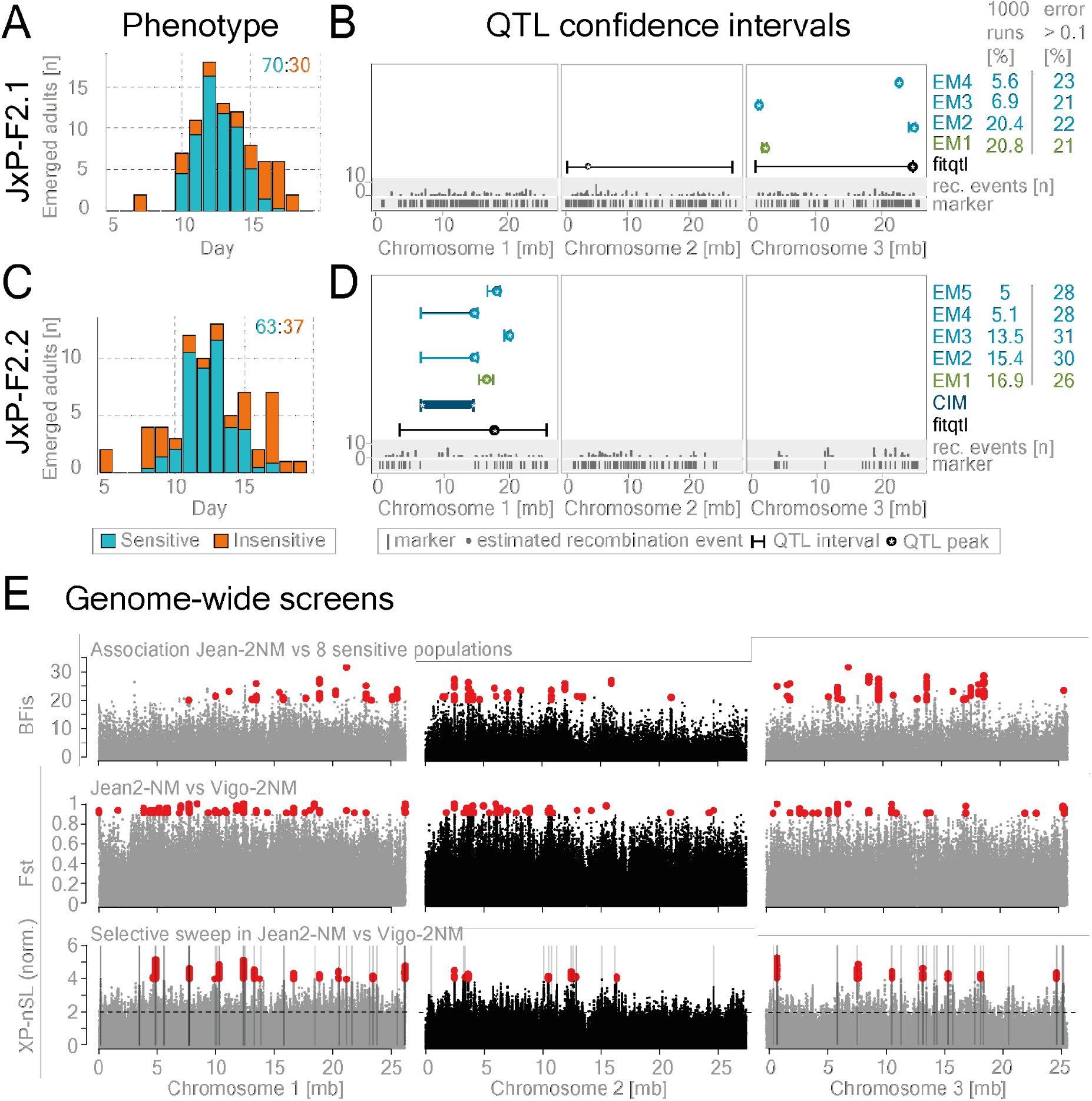
The oligogenic basis for the loss of sensitivity to tidal turbulence in the Jean-2NM population. QTL mapping in two Jean-2NM x Por-1SL intercross families was performed to find genomic regions harboring genes responsible for the loss of mechanical entrainment in Jean-2NM (A-D). (A, C) The proportion of insensitive (orange) and sensitive (blue) individuals found on each day was calculated based on estimated probabilities (Supplemental Figure 14). The ratio of the two phenotypes is indicated in the top right corner. (B, D) QTL confidence intervals are given for: composite interval mapping – dark blue, fitqtl: additive loci – black, fitqtl: EM-algorithm – light blue. The green marks the phenotypic panel with the highest convergence in EM analysis, and the lowest error (see methods QTL mapping/EM-pipeline, Supplemental Table 8). (E) Association analysis was performed to find mutations associated with the loss of sensitivity to tidal turbulence in the Jean-2NM population. (E-top): We screened for variants associated with the loss of sensitivity to tidal turbulence using 769.379 variants called from 210 individuals belonging to 9 differentially sensitive populations (Supplemental Figures 1 and 17). Bayesian factor (BFis) is plotted for each variant along the three chromosomes. We found 173 significantly associated SNPs and indels (BFis > 20, eBPis > 2, XtXst > 20.02; see Methods section for details) marked in red. The list of SNP effects and genes affected by them is given in Supplemental Table 9). (E-middle and bottom) To find loci under selection in Jean-2NM that could be responsible for the loss of sensitivity, we contrasted it with the closest turbulence-sensitive population Vigo-2NM. (E-middle) plots show the results of genomic differentiation analysis (Fst) between Vigo-2NM and Jean-2NM. Red marks Fst values above 0.5. (E-bottom) Plots depict the results of the selective sweep analysis in Jean-2NM as compared to the Vigo-2NM. The top 1% 10kb regions under selection are gray.

QTL mapping was conducted in two intercross families: in the JxP-F2.2 family we found one additive QTL on the first chromosome, while in the JxP-F2.1 family two additive QTLs on chromosomes 2 and 3 appeared (Figure 4 A-D, Supplemental Figure 14-15, Supplemental Table 8). Interestingly, while the ratio of sensitive to insensitive individuals was consistently 73:27 in three independent intercross families including JxP-F2.1 (Supplemental Figure 16), the JxP-F2.2 family had a unique ratio of 62:38 (Supplemental Figure 16). This could indicate that the genetic basis for the insensitivity in JxP-F2.2 is unique and may explain why we found different QTLs in JxP-F2.1 and JxP-F2.2. In addition, this finding suggests that there is an oligogenic origin for the trait and that the alleles responsible for the loss of sensitivity are not fixed in the Jean-2NM population.

We then performed the same genetic screens in Jean-2NM as in Ros-2FM (Figure 4 E). The association analysis identified 173 SNPs significantly associated with the phenotypic loss in Jean-2NM as compared to the eight sensitive populations (Figure 4 E, Supplemental Table 9). As in the Ros-2FM association analysis, most associated SNPs are found in the non-coding regions of the genome (Supplement Figure 17). To investigate the potential selective sweep in Jean-2NM, we tried contrasting it with the closest turbulence-sensitive population we had: Vigo-2NM (Figure 4 E, Supplemental Figure 1). However, since the two populations are geographically quite far from each other, the F_ST_ values were very high overall (Figure 4 E). Thus, Vigo-2NM is not the most suitable reference population for discovering reliable selective sweeps in Jean-2NM. Taken together, due to the complexity of the QTL mapping results in Jean-2NM, as well as the lack of prominent peaks in the association analysis, we were not able to identify candidate genes with enough precision. Nevertheless, the absence of a prominent QTL on chromosome 2 in Jean-2NM corroborates the finding that this phenotype was lost independently in Ros-2FM and Jean-2NM.

## DISCUSSION

### Loss of sensitivity to mechanosensory entrainment: result of convergent evolution?

Loss-of-function alleles were once only associated with deleterious mutations, and loss of genes with the loss of redundant gene duplications. It is now understood that the loss of alleles and genes can drive adaptive phenotypic diversity (Monroe et al. 2021; Albalat and Cañestro 2016). Furthermore, in contrast to the early evolutionary theories, we now come to understand that convergent evolution is more of a rule than an exception. A few recent studies show that the loss of traits can appear as a result of convergent evolution: repeated eye loss in Mexican cavefish (Sifuentes-Romero et al. 2020) and the loss of flight in paleognathous birds (Sackton et al. 2019).

*Clunio* colonized the European Atlantic coast from south to north following the last ice age about 10.000 to 20.000 years ago (Kaiser et al. 2010). Vigo-2NM is the most southern population tested in the laboratory and it is sensitive to tidal turbulence (Figure 1 A, Supplemental Figure 1 U). This hints that the ancestral *Clunio* population was likely also sensitive, and the insensitivity in certain populations can be considered a loss of trait. The adaptive value of this loss remains speculative (see below). However, we now know that it has evolved independently in the two *Clunio* populations. The obvious difference in Ros-2FM and Jean-2NM in identified QTLs (Figures 2 and 4 A-D) and the positions of the associated SNPs (Figures 3 and 4 E), corroborates the results from the complementation cross (Figure 1 C) and leads to the strong conclusion that this trait indeed evolved independently in the two locations. Thus, if this phenotype has adaptive value, we have here uncovered an example of recent convergent evolution in the process of local adaptation to different timing habitats.

### Evolution of differential sensitivity to circalunar synchronizers

In agreement with Neumann (Neumann and Heimbach 1978), the northern populations (Por-1SL, Bria-1SL, He-1SL, and Ber-1SL) are very sensitive to tidal cues while southern ones are less sensitive or entirely insensitive (Jean-2NM, Plou-2NM, Ros-2NM, Ros-2FM, and Lou-2NM). He argued that moonlight is an ill-suited zeitgeber in the north due to the low position of the moon on the horizon (Neumann and Heimbach 1978). However, that does not explain why tidal turbulence would be unreliable in the south. There is no obvious advantage for Ros-2FM or Jean-2NM to lose the sensitivity to this cue since the tides are as strong and predictable in those locations as in any other tested location.

We also observed that populations most sensitive to the tidal turbulence have a semi-lunar period in adult emergence, i.e. they emerge twice a lunar month, while less sensitive populations have a lunar period (except Vigo-2NM). Tidal turbulence is a semi-lunar zeitgeber as it comes from the tides, and it could therefore be a more appropriate cue for the populations emerging twice a month in contrast to those that are emerging once a month for which moonlight, as a monthly zeitgeber, might be a more suitable cue.

Furthermore, we discovered that the two sympatric populations in Roscoff are differentially sensitive: Ros-2NM, sensitive to tidal turbulence, and Ros-2FM, insensitive to tidal turbulence (Supplemental Figure 1, M and O). Although we tested the two zeitgebers moonlight and tidal turbulence separately in the lab, they are perceived together in the wild. Furthermore, as the timing of the tides changes along the Atlantic coast, their phase-relationship varies in different habitats (Kaiser et al. 2011). Thus, if the two zeitgebers set the phase differently, losing sensitivity to one of them can be an evolutionary strategy to set the phase according to the most informative zeitgeber. In line with that, we find the same QTL locus harboring STAT-1 and CG022363 as one of the QTLs responsible for the phase-difference between Ros-2FM and Ros-2NM (manuscript submitted, Briševac et al. 2022).

### Genes responsible for the loss of sensitivity to tidal turbulence in Ros-2FM

Tidal turbulence is a vibration, perceived by the mechanosensory nervous system, and mechanosensory pathways are even in model organisms still largely unknown. Most molecular players were identified in genetic screens on phenotypes associated with defects in mechanosensory systems in *Drosophila melanogaster, Caenorhabditis elegans*, and *Mus musculus* (Ernstrom and Chalfie 2002). Genes found in these analyses are not only directly involved in mechanosensation: ion channels, tethering of the ion channels, extracellular matrix, cytoskeleton; but also indirectly in the development of the sensory organs or the function and development of the cells downstream in the neuronal circuits (Ernstrom and Chalfie 2002). Many complex phenotypes are polygenic in origin, which makes simple gene–function relationships hard to infer. Additionally, mutations in regulatory regions, rather than mutations in coding regions, are found to shape most emerging phenotypes (Sackton et al. 2019). Similarly, the loss of complex phenotypes has been shown to be driven by divergence in cis-modulatory elements of developmental genes in the loss of limbs in snakes and degeneration of eyes in subterranean mammals (Roscito et al. 2018). Therefore, we investigated both, the region of the genome with the highest association score (Figure 3 C), but also other potential candidate genes (Supplemental Figures 10-13, Supplemental Tables 6-7).

#### STAT-1 locus

Signal Transducer and Activator of Transcription (STAT) protein is a part of the evolutionary conserved JAK-STAT pathway that controls developmental decisions and participates in the immune response (Wang and Levy 2012). Archetypical members of each of the components were present at the time of the emergence of Bilateria: JAK, STAT, SHP, and the three SOCS proteins (Liongue and Ward 2013). STAT proteins were duplicated many times throughout metazoan evolution, and while some pseudogenized, many evolved into novel genes through rapid sequence diversification and neofunctionalization (Wang and Levy 2012). Insect STATs form a single clade in phylogenetic analyses and constitute an ancient class of STATs together with mammalian STAT5 and 6 (Wang and Levy 2012). While most insect species like *Drosophila melanogaster* and *Apis mellifera* have a single STAT whose function remains conserved (Wang and Levy 2012), in others like *Anopheles gambiae* STAT duplicated and the new gene acquired diverse functions. Duplicated *Anopheles* STAT has a role in defense against bacteria (Barillas-Mury et al. 1999), *Plasmodium* infection (Gupta et al. 2009), and innate immunity (Souza-Neto et al. 2009). In addition, duplicated STAT acts as an upstream regulator of the evolutionarily conserved STAT protein (Gupta et al. 2009).

In contrast, in vertebrates, all components of the JAK-STAT pathway duplicated several times and STAT proteins attained specialized functions in various cells. Interestingly, the expression of the TrpA1 mechanosensitive channel is regulated via the JAK-STAT pathway in nociceptive neurons in mice (Malsch et al. 2014). Similarly, STAT3 is necessary for the differentiation and regeneration of inner ear hair cells, the basic mechanosensory receptors for hearing and balance in zebrafish (Liang et al. 2012).

Finally, the JAK-STAT pathway is directly coupled to the mechano-gated channels in various non-neuronal cells, regulating gene expression downstream of the channel activation (Lammerding et al. 2004; Millward-Sadler et al. 2006; Shah et al. 2010; de Andrés et al. 2011; Busch-Dienstfertig and González-Rodríguez 2013; Kunnen et al. 2018).

*Clunio marinus* has two STAT proteins: CG012971 (STAT-1), the ortholog of *Anopheles gamibiae* STAT2, and *Mouse* STAT5a,5b and 6; and CG022905 (STAT-2) which based on phylogenetic analysis appears to be a novel *Clunio* duplication (Figure 3 D). Two *Clunio* STATs are 83% identical in amino-acid sequence (Figure 3 E), while *Anopheles* STATs are almost identical in protein length but share only 47% overall sequence identity (Wang and Levy 2012). Two *Clunio* STATs differ the most in the N-terminal domain which has a role in nuclear translocation and protein-protein interactions, and the coiled domain which is involved in nuclear export and regulation of tyrosine phosphorylation (Liongue and Ward 2013). This indicates that the two STATs could be regulated differently or be a part of different signalling pathways by interacting with different proteins and thus obtaining different roles.

Taken together, we can speculate that *Clunio* STAT-1 has a role in the perception of tidal turbulence by being involved in the development or differentiation of mechanosensory organs, or mechanosensory receptors appropriated this JAK-STAT pathway for regulation of gene expression. Further functional analysis is necessary to test this hypothesis. If proven, this would be the first evidence of a STAT role in mechanosensation in invertebrates.

#### Gravitaxis: potential role of chordotonal organs

CG022363 also falls into the region with the highest SNP density together with STAT-1 (Figure 3 C). This gene is an ortholog of the *Drosophila CG32100* gene, which has a role in gravitaxis although the exact molecular function of this gene remains unknown (Armstrong et al. 2006). Graviception is a function of the mechanosensory system, and as is the case with all mechanosensory functions, it is poorly understood on a molecular level. To this day, most of the molecular machinery was identified through genetic screens of behaviors associated with impaired gravitaxis (Armstrong et al. 2006). As a result, 27 genes were associated with gravity-sensing in *Drosophila*: some in detecting gravity directly: *inactive, nanchung, painless*, and *pyrexia* (Sun et al. 2009); but the majority seem to have an indirect role most likely in the development of the sensory organs: *alan shepard, escargot, broad, cryptochrome, nemo* and others (Sun et al. 2009; Armstrong et al. 2006). Strikingly, out of those 27 genes, we found 6 that were associated with loss of sensitivity to tidal turbulence in Ros-2FM (Supplemental Table 7): *shep, snaill1_CG000103, broad, cry1, nmo*, and the above-mentioned *CG022363*. In line with that, gravitaxis was found as one of the top GO terms (Supplemental Figure 13). Three of the six belong to the 15 candidate genes under the QTL regions: *shep, snaill1_CG000103*, and *CG022363* (Supplemental Figure 10). In *Drosophila shep* is involved in neuronal development and remodeling of the sensory neurons (Chen et al. 2014; Olesnicky et al. 2018) and *escargot* has a role in neurogenesis (Ashraf et al. 1999). Therefore, it is likely that they are indirectly involved in gravitaxis in *Drosophila* by contributing to the development of the sensory organs responsible for detecting gravity. *Drosophila* larvae detect both vibration and gravity via chordotonal organs (Kamikouchi et al. 2009; Ishikawa et al. 2020). In addition, chordotonal organs are necessary for the mechanosensory entrainment of the circadian clock in *Drosophila* adults (Simoni et al. 2014). Taken together, it is possible that chordotonal organs are responsible for mechanosensory entrainment of the circalunar clock in *Clunio* as well. Mutations in genes responsible for the development of the chordotonal organs could lead to impaired gravity sensing as well as detection of vibration and thus impair mechanosensory entrainment of the circalunar clock in Ros-2FM.

Here we show for the first time a convergent loss of sensitivity to tidal turbulence in two *Clunio* populations. We found several loci to be responsible for this loss. A detailed analysis suggests that in one of the populations the JAK/STAT pathway and gravitaxis may play a prominent role in the detection of tidal turbulence. While in Baltic and Northern European populations complete lunar arrhythmicity seems to be a highly polygenic trait (Fuhrmann et al.), the selective loss of sensitivity to a zeitgeber seems to have a less complex, oligogenic basis. If in the future tools for molecular manipulation of *Clunio* are developed, this setting is a good starting point to identify novel genes and pathways involved in mechanosensation.

### METHODS

#### *Clunio* cultures

*C. marinus* cultures were established from different locations (Supplemental Table 1) and maintained in the laboratory according to Neumann (Neumann 1966). Around 1000 larvae were kept in 20×20×5 cm plastic boxes with sand from the natural habitat and 15‰ seawater. They were fed twice a week with diatoms (Phaeodactylum tricornutum, strain UTEX 646). Nettle powder was added twice a month with each water exchange. *Clunio* larvae were raised under a 16h light and 8h darkness regime and a temperature of 18°C. In experiments with moonlight entrainment, the artificial moonlight was simulated with neutral white LED ~4000K light (Hera 610 014 911 01) on 4 consecutive nights every 30 days. The 24-hour period when moonlight was first applied was marked as day 1. In the experiments with mechanosensory entrainment, cycles of vibration were used to simulate tidal turbulence in a setup established by Neumann (Neumann 1978). Briefly, an electromotor generating vibration of 50 Hz, 30dBa above background noise was attached to the shelves with *Clunio* cultures and controlled by a custom-made “tidal clock”. The clock kept the motor on for 6h 10 min and off for 6h 15min which gave a 12.4-hour tidal rhythm. The onset of vibration shifted every day by 50 min which resulted in a 14.7-day semi-lunar cycle. The day when vibration started in the middle of their subjective night was arbitrarily marked as day 1. Phenotypes were recorded by collecting emerged adults from three culture boxes per strain every day for at least 60 days or two lunar cycles.

#### Crossing experiments

To explore the genetic basis of insensitivity to tidal turbulence, we crossed the insensitive Ros-2FM or Jean-2NM strains with the sensitive Por-1SL strain, as well as the two insensitive strains with each other (Supplemental Table 3). Detailed description can be found in the Supplemental methods.

#### QTL mapping

QTL mapping was performed to identify genetic regions harboring genes where natural variants that underlie the loss of sensitivity to tidal turbulence are segregating. Two families from the Ros-2FM x Por-1SL cross were chosen for QTL mapping: Ros-2FMxPor-1SL-F2-24 in the further text referred to as RxP-F2.1, and Ros-2FMxPor-1SL-BC-15 in the further text referred to as RxP-BC. 1 (Supplemental Table 3). Similarly, two intercrosses of Jean-2NM x Por-1SL families were selected: Jean-2NMxPor-1SL-F2-8.6 and Jean-2NMxPor-1SL-F2-11.3 in the further text referred to as JxP-F2.1 and JxP-F2.2.

##### Phenotyping (Supplemental Figure 3)

Emergence data was collected for parental, F1 and F2, and BC generations, and lunar emergence days under turbulence entrainment were assigned as described above (Script 1, Supplemental Table 2, Supplemental Figures 1 and 2). To resolve the problem of the overlapping “sensitive” and “insensitive” phenotypes emerging during the peak in the F2 and BC progeny, we designed a pipeline to calculate the probability of finding “sensitive” and “insensitive” individuals on each day. For more details see Supplemental Methods.

##### Genotyping

DNA was extracted from adults collected in crossing experiments with the salting-out method (Reineke et al. 1998), it was aplified using RepliG, and single-digest or double-digest RAD sequencing was performed (Etter et al. 2011; Etter and Johnson 2012; Baird et al. 2008). A detailed protocol can be found in the Supplemental methods. The script containing read processing, mapping, genotype calling and filtering for informative variants is given in the supplement as Script 2.

###### Read processing and mapping

For discriminating individuals, P1 and P2 adaptors contained unique barcode sequences (Supplemental Table 10). Raw reads were trimmed to remove adapters and low-quality bases with Trimmomatic v0.38 (Bolger et al. 2014). Trimmomatic parameters used for paired-end reads were ILLUMINACLIP:<PE_adapter_file>:2:30:10:2:true LEADING:20 TRAILING:20 MINLEN:50 and for single-end reads ILLUMINACLIP:<SE_adapter_file>:2:30:10 LEADING:20 TRAILING:20 MINLEN:50. For paired-end library RxP-F2.1, overlapping read pairs were assembled into single reads with PEAR v.0.9.10 (Zhang et al. 2014) using default parameters. Paired (PEAR unassembled) and single reads (PEAR assembled and unpaired reads from R1 or R2 after adapter trimming) were mapped independently with NextGenMap v0.5.5 (Sedlazeck et al. 2013) to the CLUMA2.0 reference genome (available at https://doi.org/10.17617/3.42NMN2; currently unpublished) with default parameters except for --min-identity 0.9 and --min-residues 0.9. Read groups were specified during mapping using --rg-id and --rg-sm. The independently mapped reads were then merged into a single file using samtools v1.9 (Li et al. 2009) merge, with parameters -u -c -p. For single-end libraries, trimmed reads were directly mapped with NextGenMap with previously mentioned parameters. Mapped reads were sorted and indexed with samtools sort and samtools index respectively.

###### Variant calling

Single nucleotide polymorphisms (SNPs) and insertion-deletion (indel) genotypes were called using GATK v3.7-0-gcfedb67 (McKenna et al. 2010). Steps include initial genotype calling using GATK HaplotypeCaller with parameters --emitRefConfidence GVCF and -stand_call_conf 30, filtering of variants using GATK SelectVariants with ‘-select DP > 30.0’, recalibration of base qualities using GATK BaseRecalibrator with ‘-knownSites’, preparing recalibrated BAM files with GATK PrintReads using -BQSR and finally, recalling of genotypes using GATK HaplotypeCaller with previously mentioned parameters. Individual VCF files were combined into a single file using GATK GenotypeGVCFs.

###### Informative variants and genotype matrix

VCF files were filtered for minimum genotype quality (minGQ) 20, minor allele frequency (maf) 0.10, and maximum fraction of samples having missing genotypes (max-missing) 0.60. Genotypes were coded as ‘AA’, ‘AB’, or ‘BB’ based on the inferred inheritance pattern (Supplemental Table 11). To maximize the number of informative markers in a backcross, we included markers for which both parents were heterozygous or, the F1 parent was heterozygous and the Ros-2FM parent was homozygous (Supplemental Table 11. To infer from which parent the ‘A’ or ‘B’ allele comes from at ambiguous loci, we chose genotypes based on the consistency of the genotypes along the chromosome (i.e. the assignment that had a smaller number of genotype switches across the individuals in the BC progeny) (Supplemental Table 11, consistency genotype assignment in Script 2). Our final genotype matrix was manually inspected before importing it into R/qtl. Marker order and genotype errors were further investigated in R/qtl. One inversion was identified in the right arm of the second chromosome in the RxP-F2.1 family and the order of markers was inverted in that region. The final number of markers was 117 for RxP-BC.1 and 137 for the RxP-F2.1 family. The final genotype matrix is given in Supplemental Table 4.

Samples from parents’ and F1s of the two Jean-2NMxPor-1SL families, unfortunately, had very few good genotypes. Thus, we designed an alternative approach for reconstructing the recombination matrix. For details see Supplemental Methods.

##### QTL mapping

Standard interval mapping and multiple QTL mapping were done with R/qtl package functions: *scanone, scantwo*, and *fitqtl* (Karl W. Broman and Saunak Sen 2009). QTL intervals were estimated with *bayesint* function. Composite interval mapping was analyzed in Windows QTL Cartographer Version 2.5_011 (number of covariates 5, window 10 cM) (Wang et al. 2012). In addition, to confirm the model found by *fitqtl* multiple QTL mapping, we used the Bayesian QTL mapping R package “qtlbim” (Yandell et al. 2007). Function *qb.best* was used to identify the best model, and *qb.scanone* to compare additive and epistatic QTLs found by R/qtl.

###### Expectation-maximization (EM) algorithm (Supplemental Figure 5)

To explore the effect of uncertainty in phenotyping on the QTL mapping results, we devised an EM algorithm to assign binary phenotypes to the entire dataset (Supplemental Figure 5). For details see Supplemental Methods. The script can be found in the supplement as Script 3.

#### Association analysis

The complementation cross indicated that the genetic basis for the loss of sensitivity is different in the two populations (Figure 1). Therefore, the two insensitive populations were analyzed separately. To identify variants associated with the loss of sensitivity to tidal turbulence in Ros-2FM we performed a genome screen on 746.887 SNPs and small indels called in 210 males from nine populations differentially sensitive to tidal turbulence (Supplemental Table 5). Similarly, to find potentially causative mutations in Jean-2NM, we used a dataset of 769.379 SNPs and indels from 210 males from Jean-2NM and the same eight populations sensitive to tidal turbulence.

##### Genotyping

DNA from field-caught males stored in 100% ethanol was extracted using the salting-out method (Reineke et al. 1998). Genomic DNA was amplified with standard RepliG protocol (REPLI-g Mini Kit QIAgen 150025). Whole genomes of 20-24 adults from nine populations were sequenced on Illumina HiSeq3000 with paired-end 150-bp reads (Supplemental Table 5).

###### Read processing

Reads from several sequencing runs were merged with the *cat* function. Adapters were trimmed using Trimmomatic tool (Bolger et al. 2014) and the following parameters: ILLUMINACLIP <Adapter file>:2:30:10:8:true, LEADING:20, TRAILING:20, MINLEN:75. Overlapping read pairs were assembled using PEAR with the following parameters: -n 75 -c 20 -k (Zhang et al. 2014). Reads were mapped using bwa mem version0.7.15-r1140 (Li and Durbin 2009) using the latest Cluma_2.0 reference genome (available at https://doi.org/10.17617/3.42NMN2; currently unpublished; private url for viewing it during the review process: https://edmond.mpdl.mpg.de/privateurl.xhtml?token=79417ae6-4696-4f31-b436-16cd358905f4). The independently mapped reads were then merged into a single file, filtered for -q 20, and sorted using samtools v1.9 (Li et al. 2009).

###### Variant calling

SNPs and small indels were called using GATK v3.7-0-gcfedb67 (McKenna et al. 2010). All reads in the q20 sorted file were assigned to a single new read-group with the ‘AddOrReplaceReadGroups’ script with LB=whatever PL=illumina PU=whatever parameters. Genotype calling was then performed with HaplotypeCaller and parameters --emitRefConfidence GVCF -stand_call_conf 30, recalibration of base qualities using GATK BaseRecalibrator with ‘-knownSites’. Preparing recalibrated BAM files with GATK PrintReads using -BQSR. Recalling of genotypes using GATK HaplotypeCaller with previously mentioned parameters. Individual VCF files were combined into a single file using GATK GenotypeGVCFs.

###### BayPass genotype matrix

The genotype matrix for BayPass association analysis (Gautier 2015) was generated by filtering for minor allele frequencies larger than 0.05, the maximal number of missing values per variant was set to 20%, the maximal number of alleles was 2, and minimal read quality minQ was set to 20 with VCFtools (0.1.14) (Danecek et al. 2011). Allele count per population was calculated using the VcfR package (Knaus and Grünwald 2017). Briefly, a previously filtered vcf table containing 24 individuals from 9 populations was separated into vcf files containing individuals from distinct populations. Individual vcf files were read with read.vcfR function and allele frequency per population per site was calculated using the gt.to.popsum function. Population allele frequencies were then combined into a genotype matrix.

##### Phenotyping

Sensitivity to turbulence was estimated for each population using summary circular statistics (see methods section QTL mapping/Phenotyping). Vector length was used as a phenotypic score (Supplemental Table 2).

##### BayPass

BayPass was run with 3 random seeds (1, 1988, 11273), and the median of BFis, eBPis, XtXst, and −log10 p-value of XtX was calculated. To find the correct significance threshold for XtX statistics, pseudo-observed data set (POD) was generated by sampling 100.000 SNPs with R function *simulate.baypass* and found that 1% of XtXst POD values was 21.67 in Ros-2FM dataset, and 20.02 in Jean-2NM. To subset highly associated variants in Ros-2FM, we filtered for BFis >= 20, eBPis >= 2 and XtXst >= 21.67 (Supplemental Figure 9 A) and BFis >= 20, eBPis >= 2 and XtXst >= 20.02 in Jean-2NM (Supplemental Figure 17 C). Association analysis in BayPass is corrected for the population structure based on a kinship matrix Ω.

### SNPeff

SNP effects were analyzed in CLUMA2.0_M, a version of the reference genome that contains manual curations to the reference sequence made during genome annotation (available at https://doi.org/10.17617/3.42NMN2; currently unpublished). SNPs were transferred from CLUMA2.0 to CLUMA2.0_M using a Python3 script (Script 4), which creates a map of positions from CLUMA2.0 to CLUMA2.0_M by accounting for insertions and deletions. As input, the script uses a GFF file with manual reference edits, exported from Web Apollo version 2.5.0 (Lee et al. 2013). With the CLUMA2.0_M reference sequence, the location and putative effects of the SNPs and indels relative to CLUMA2.0_M gene models were annotated using SnpEff 4.5 (build 2020-04-15 22:26, non-default parameter ‘-ud 0’) (Cingolani et al. 2012). The complete list with the number of variants with distinct effects is given in Supplemental Tables 6 and 9.

### Phylogenetic trees

The identity of the 15 candidate genes was explored by the reciprocal blast between *Clunio* and *Drosophila melanogaster* protein sequences. eggNOG 5.0 database was then used to identify orthologs in other model organisms: *Anopheles gambiae, Mus musculus, Homo sapiens*, and *Caenorhabditis elegans* (Huerta-Cepas *et al*. 2019). The most distant protein sequence in eggNOG phylogenetic trees was taken as an outgroup sequence. Protein sequences were then aligned, and phylogenetic trees were created in QIAGEN CLC Main Workbench version 7.9.3. Bootstrap values in 1000 runs were reported (Figure 3 D,E, Supplemental Figures 11 and 12).

### Selective sweep analysis

To investigate if the associated loci evolved as a result of a recent selective sweep in the process of local adaptation, we calculated cross-population nSL (number of segregating sites by length) developed by (Szpiech et al. 2021). XP-nSL is designed to detect selective sweeps due to local adaptation within a query population by comparing its integrated haplotype homozygosity (iHH) with one of a reference population. Here, positive scores suggest long haplotypes in population A with respect to population B and a potential sweep in A, whereas negative scores suggest long haplotypes in B with respect to A. nSL, in contrast to EHH, was developed to accommodate the lack of genetic maps in favor of physical maps (Ferrer-Admetlla et al. 2014). We used selscan 2.0 as it was recently revised to work with unphased multi-locus genotypes (DeGiorgio and Szpiech 2021; Szpiech et al. 2021). Details can be found in the Supplemental Methods.

### Genetic differentiation (fst)

To provide a bridge between the association analysis conducted on ten populations, and the cross-population selective sweep analysis calculated between the two populations (see Methods section on association analysis and selective sweeps), we estimated genetic differentiation between those two contrasted populations: Ros-2FM compared to Ros-2NM and Jean-2NM compared to Vigo-2NM. The same vcf files containing GATK-called SNPs and indels used for selective sweep analysis were used (see Methods section on selective sweeps). Genetic differentiation between the two populations (fst) was estimated using vcftools version 0.1.14 (Danecek et al. 2011) parameters --weir-fst-pop --fst-window-size 1 --fst-window-step 1.

### GO term enrichment

To investigate if the genes identified by BayPass and SNPeff perform some of the known biological functions, we ran Gene Ontology (GO) term enrichment. We previously annotated 5,393 out of 15,193 *C. marinus* genes with GO terms (Fuhrmann et al.). In our current reference genome CLUMA2.0, 5436 out of 13751 genes were annotated with GO terms. In brief, GO terms were annotated using the longest protein sequence per gene with mapper-2.0.1.(Huerta-Cepas et al. 2017) from the eggNOG 5.0 database (Huerta-Cepas et al. 2019), using DIAMOND (Buchfink et al. 2014), BLASTP e-value <1e-10, and subject-query alignment coverage of >60%. Only GO terms with “non-electronic” GO evidence from best-hit orthologs restricted to automatically adjusted per-query taxonomic scope were used. To assess the enrichment of “Biological Process” GO terms, the weight01 Fisher’s exact test was implemented in topGO (version 2.42.0, R version 4.0.3) (Alexa and Rahnenfuhrer 2022).

## Supporting information

Supplemental Script 1

Supplemental Script 2

Supplemental Script 3

Supplemental Script 4

## DATA ACCESS

Ros-2FM and Ros-2NM sequence reads are deposited at ENA under Accession PRJEB54033. Por-1SL, He-2SL and Ber-1SL raw sequence reads are deposited at ENA under Accession PRJEB43766. Jean-2NM, Vigo-1NM, Plou-2NM, Lou-2NM, Bria-1SL raw sequence reads are deposited at ENA under Accession PRJEB55328. RAD-seq reads for QTL mapping are deposited at ENA under Accession PRJEB55328. The CLUMA2.0 reference genome is available on the Open Research Data Repository of the Max Planck Society (EDMOND) (https://doi.org/10.17617/3.42NMN2; currently unpublished)

## COMPETING INTEREST STATEMENT

The authors declare no conflict of interest.

## ACKNOWLEDGMENTS

Kerstin Schäfer assisted with RAD sequencing library preparations and Jürgen Reunert provided animal care. We thank the members of the research group Biological Clocks for their feedback for the entire duration of the project. We also thank Diethard Tautz, Guy Reeves, and Miriam Liedvogel for their feedback in the process of writing the manuscript.

This work was supported by the Max Planck Society via an independent Max Planck Research Group and by an ERC Starting Grant (No 802923) awarded to Tobias S Kaiser.

## AUTHOR CONTRIBUTIONS

DB tested sensitivity to tidal turbulence in various *Clunio* strains, performed crosses, RAD sequencing, QTL mapping, association mapping, co-conceived the EM algorithm, and wrote the manuscript. CP established genotyping pipeline for QTL mapping and ran SNPeff. TSK conceived and supervised the project, wrote the EM algorithm and edited the manuscript.

## Notes

### Competing Interest Statement

The authors have declared no competing interest.

