## Supplemental Figures for "Genetic analysis of a phenotypic loss in the mechanosensory entrainment of a circalunar clock"

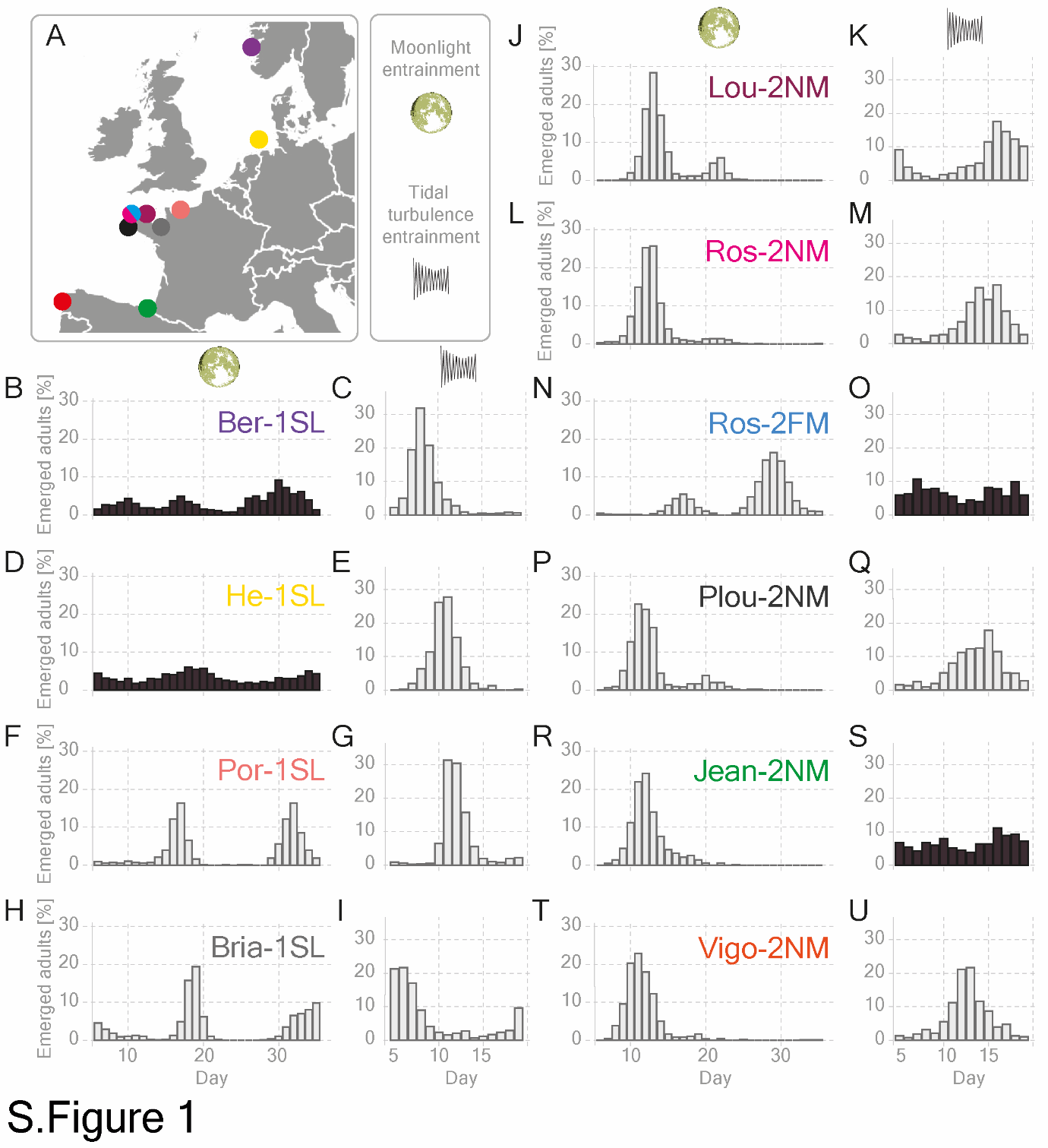

### **Supplemental Figure 1. *Clunio* populations are differentially sensitive to moonlight or tidal turbulence**

(A) Origin of the *Clunio* strains. Strains are color-coded and their names are depicted in the body of the graph. (B-U) Graphs show the fraction of emerged individuals entrained under laboratory conditions by either artificial moonlight (four nights of light every 30 days) or tidal turbulence (vibration of ~50 Hz 30dB above background noise in 6h 10min ON – 6h 15min OFF intervals resulting in a 15-day pattern). The total number of individuals, exact names of geographical locations, and the year when strains were established are given in Supplemental Table 1. Strains differ in the period, phase of emergence, and sensitivity to the synchronizers. The emergence of strains considered insensitive to moonlight or tidal turbulence is marked in black.

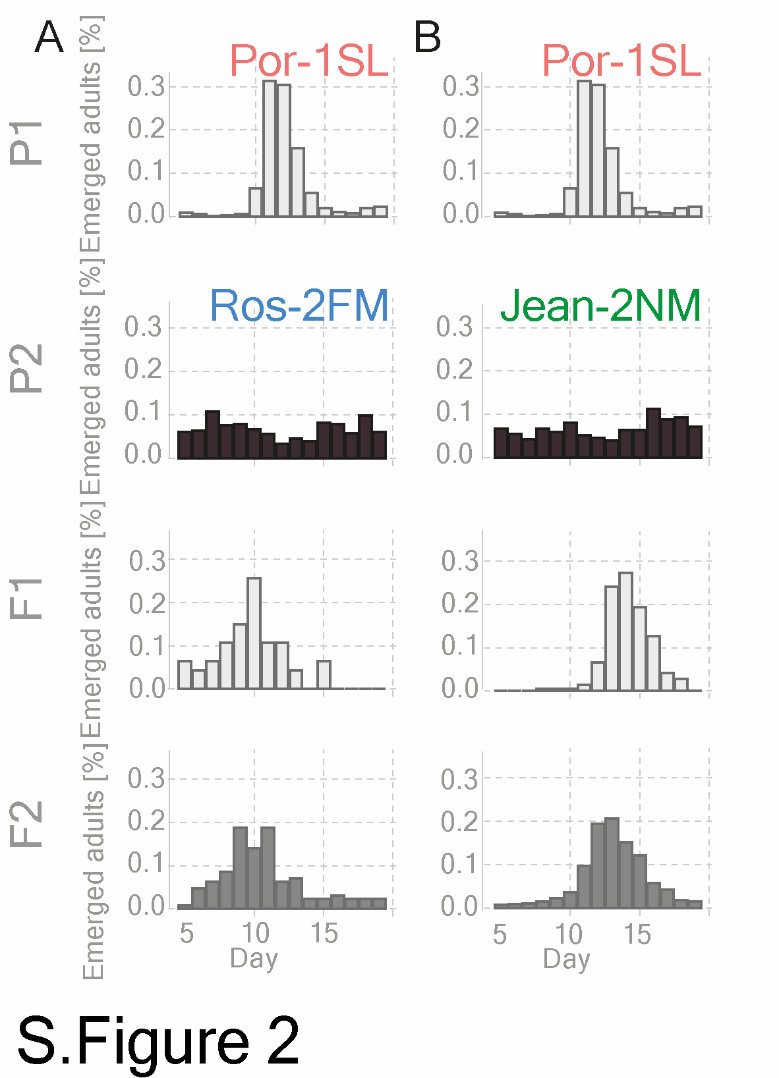

**Supplemental Figure 2. Sensitivity to tidal turbulence is genetically determined and a dominant trait.**

Crossing experiments were performed to assess the inheritance of sensitivity to tidal turbulence. Graphs show fractions of emerged adults per day in parental populations, F1 and F2 (F1xF1) generations. (A) Intercross between Por-1SL and Ros-2FM. (B) Intercross between Por-1SL and Jean-2NM. The total number of individuals per generation is listed in Supplemental Table 3. The color of the bars represents increasing levels of sensitivity to tidal turbulence from sensitive (light gray) to insensitive (black).

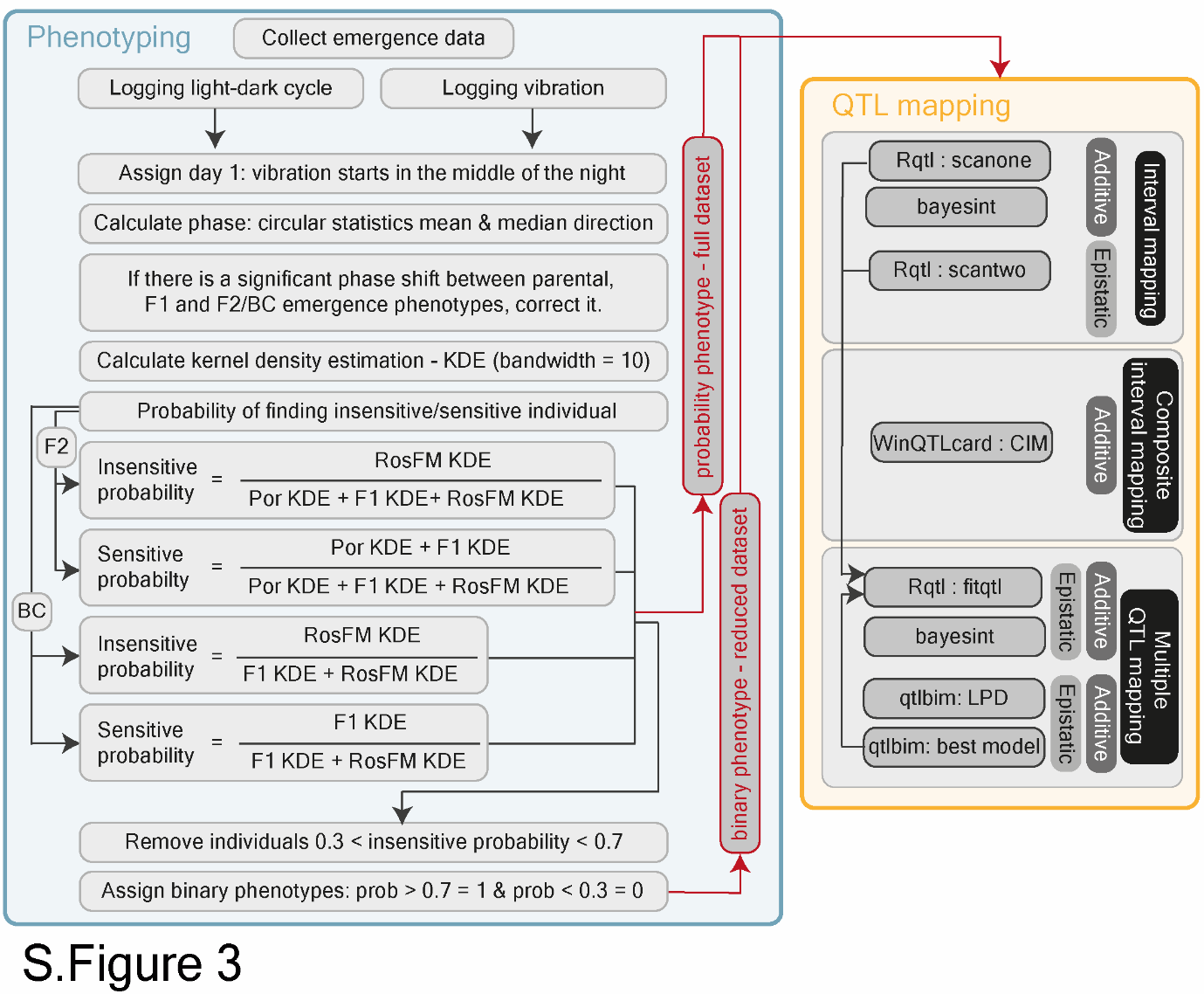

**Supplemental Figure 3. Schematic overview of the QTL mapping strategy**

Left: Phenotyping strategy. Light and vibration were logged throughout the experiment and used to calculate the first day of the entrainment as the day when vibration starts in the middle of their subjective night. Emerged adults were collected and the number of emerged individuals per day was recorded. Circular summary statistics were used to test if there was a phase-shift in emergence rhythms between parental, F1, F2, and BC generations. If there was a phase shift, emergence days were corrected so that the only phenotype assessed is rhythmicity. To generalize the emergence distributions, kernel density estimates were calculated (bandwidth = 10) for each generation. The probability of finding sensitive and insensitive individuals on each experimental day in F2 or BC progenies was calculated according to the given equations. The probability of finding an insensitive individual was used as a phenotypic score. In addition, a reduced dataset was generated by removing individuals with uncertain phenotypes between 0.3 and 0.7. Remaining individuals with probability phenotypes > 0.7 or < 0.3 were given binary phenotypes 1 and 0 respectively. Right: QTL mapping strategy. Several mapping pipelines were tested to examine additive and epistatic QTLs. Full and reduced datasets of each crossing family were analyzed using interval mapping (scanone and scantwo), composite interval mapping (WinQTL cartographer), and multiple QTL mapping (fitqtl and qtlbim). See methods section QTL mapping.

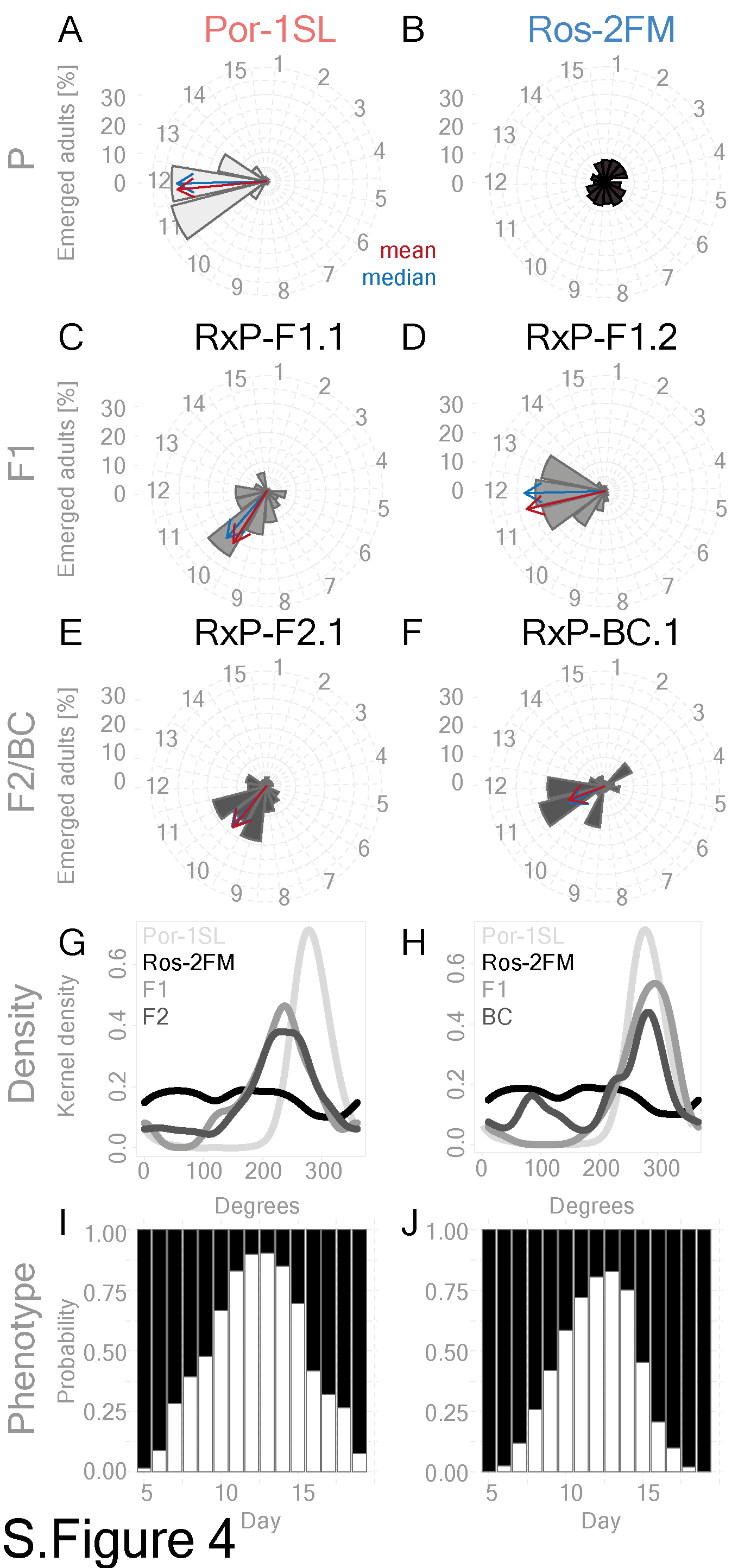

### **Supplemental Figure 4. Calculating probability phenotypes for RxP-F2.1 and RxP-BC.1 family.**

(A-F) The fraction of emerged adults per generation is shown on a circular plot together with the mean and median vectors. (A) Por-1SL strain. (B) Ros-2FM strain. (C) RxP-F1.1 and RxP-F2.1 generation. (D) RxP-F1.2 three crossing families were raised together (gave rise to RxP-BC.1). (E) RxP-F2.1 is a F1-24 x F1-24 intercross. (F) RxP-BC.1 is a backcross of an F1.2 individual to Ros-2FM. (G-H) Kernel density estimates for parental, F1, and F2/BC generations for each of the two mapping families. RxP-F2.1 crossing family shows a 2-day phase shift as compared to Por-1SL. RxP-BC.1 crossing family does not show considerable phase-shift. (I-J) Bar graphs show probabilities of finding sensitive Por-1SL-like (white) or insensitive Ros-2FM-like (black) individuals on each day in the two crossing families RxP-F2.1 (I) and RxP-BC.1 (J).

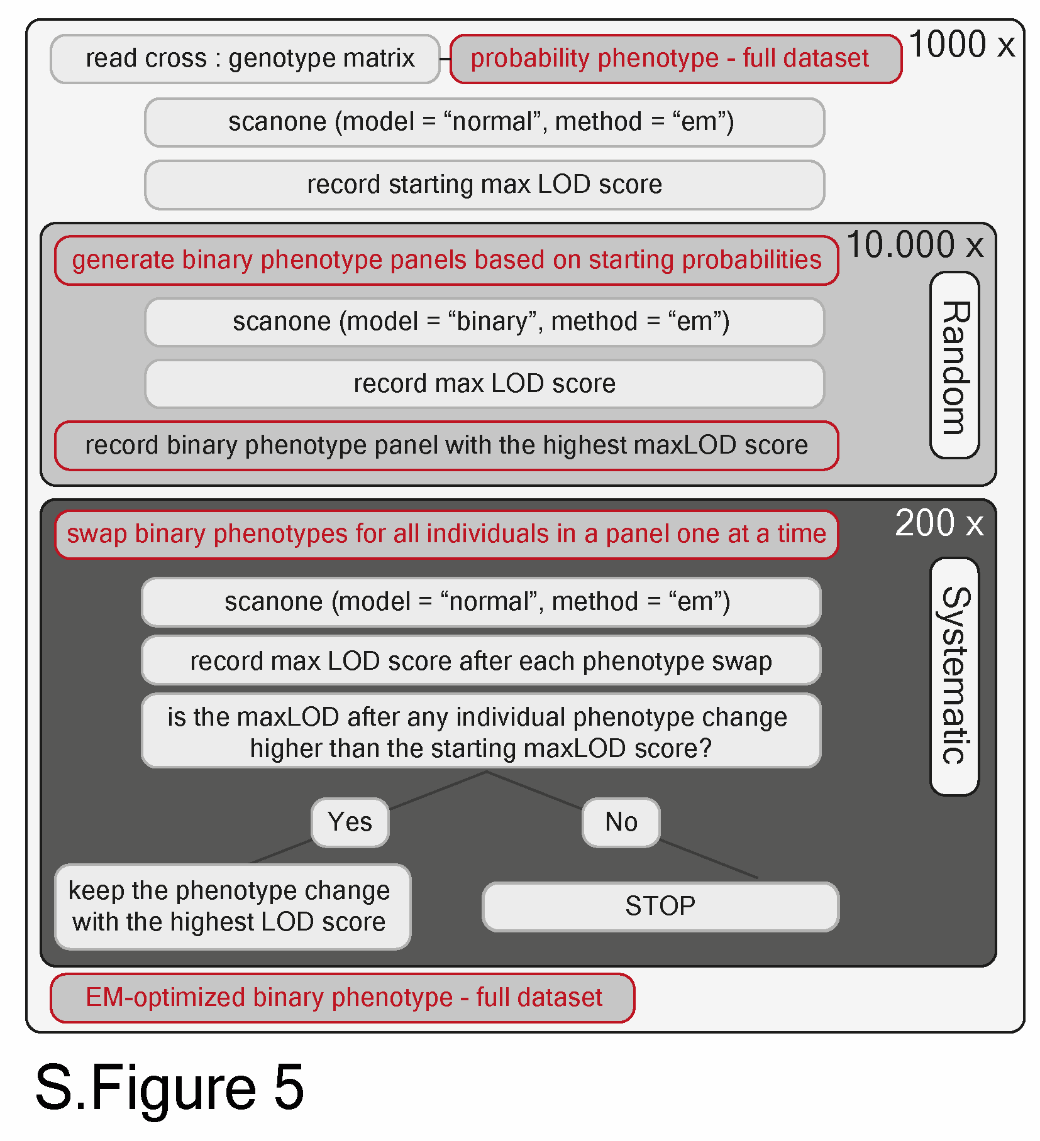

**Supplemental Figure 5. An expectation-maximization (EM) algorithm**

An EM algorithm was designed to generate optimized binary phenotype panels to a crossing family given the calculated probability of finding insensitive individuals on each experimental day. For a detailed explanation see the Methods/QTL mapping/EM Expectation-maximization (EM) algorithm paragraph.

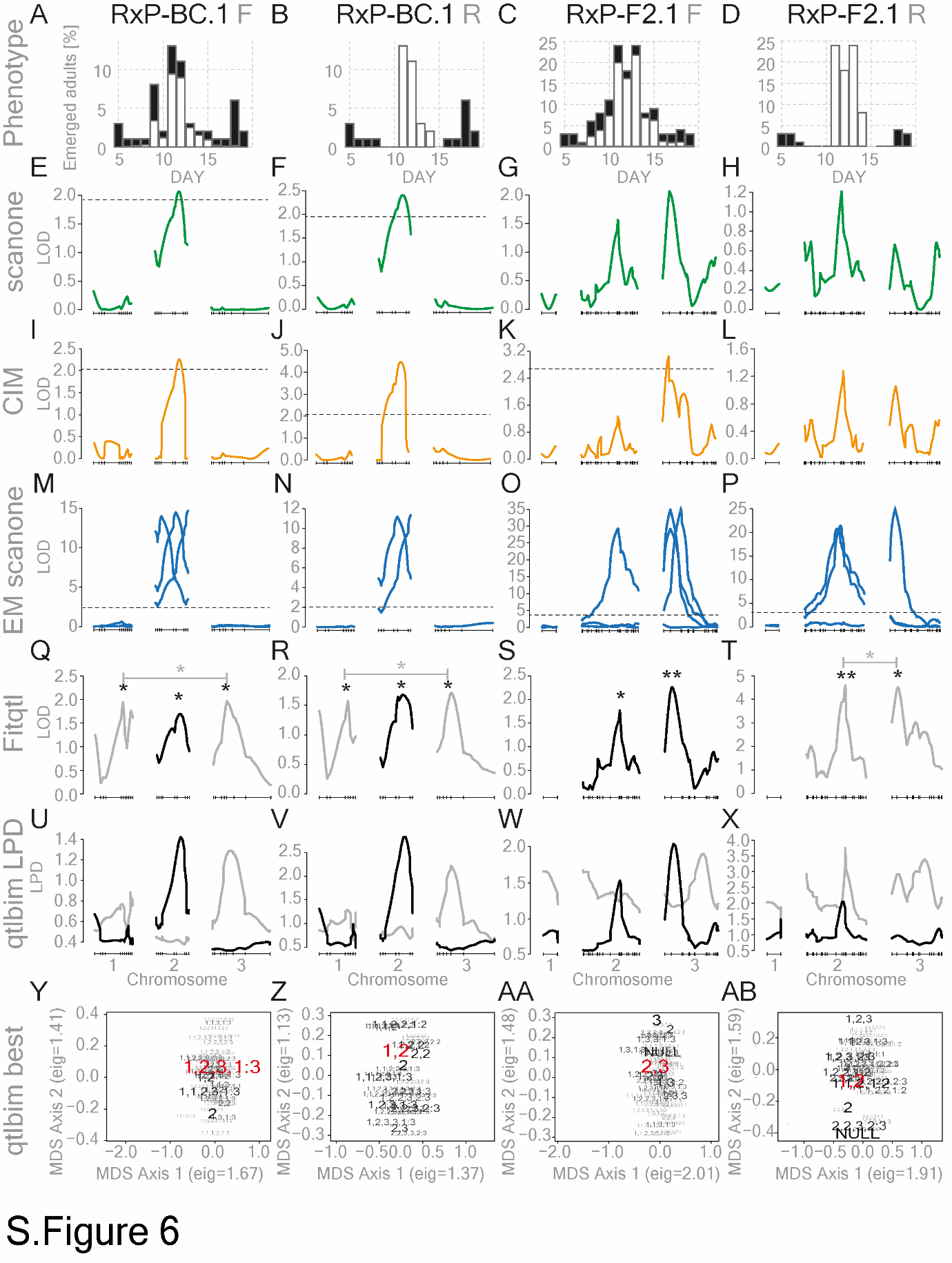

**Supplemental Figure 6. Ros-2FMxPor-1SL QTL mapping full analysis**

Complete QTL mapping results for the two crossing families (RxP-F2.1 and RxP-BC.1) and two datasets each (F = full and R = reduced) are given. (A-D) Bar graphs show the number of emerged individuals per day. The predicted ratio of insensitive (black) and sensitive (white) individuals is plotted. (E-H) LOD scores of interval mapping analysis (*scanone*) are designed to detect additive QTLs. Probability phenotypes were used for full datasets and binary phenotypes for reduced datasets. The significance threshold (dashed line) was estimated in 1000 permutations with a 5% cutoff. (I-L) CIM analysis with backward regression method, 5 control markers, and a window size of 10 cM. Threshold values are given in Supplemental Table 4. (M-P): LOD scores of scanone on EM-optimized binary phenotypes. Results are shown for panels obtained in at least 5% of the cases in 1000 runs (Supplemental Table 4.) The significance threshold (dashed line) was estimated in 1000 permutations with a 5% cutoff. (Q-T) LOD scores of significant QTLs in multiple QTL mapping pipeline (*fitqtl*). Black lines: additive QTLs, gray lines: QTLs in epistasis. p-value of F statistic is marked: * p-value < 0.05; ** p-value of <0.01. *Fitqlt* statistics are given in Supplemental Table 4. (U-AB) To find the best model for multiple QTL mapping (*fitqtl*) we also tried the *qtlbim* package. (U-X) We first ran LPD (Log Posterior Density) scan that uses Bayesian model averaging to explore the most probable models. Most likely additive QTLs are shown in black and most likely QTLs in epistasis are shown in gray. (Y-AB) The “*best*” function of the *qtlbim* package then selects the most probable model. The larger the font size the larger posterior probability the pattern has. The 2-D multidimensional scaling (MDS) projection is based on the square of the attenuation. If the loci agree exactly, there is no attenuation. The best model is marked in red. The numbers represent chromosomes. (U) The best model for the RxP-BC.1 full dataset contains 3 QTLs and one epistatic interaction on chromosomes 1, 2, 3, and 1:3. (V) The best model for the RxP-BC.1 reduced dataset is 1, 2. (W) The best model for the RxP-F2.1 full dataset is 2, 3. (X) The best model for RxP-F2.1 reduced dataset is 1, 2.

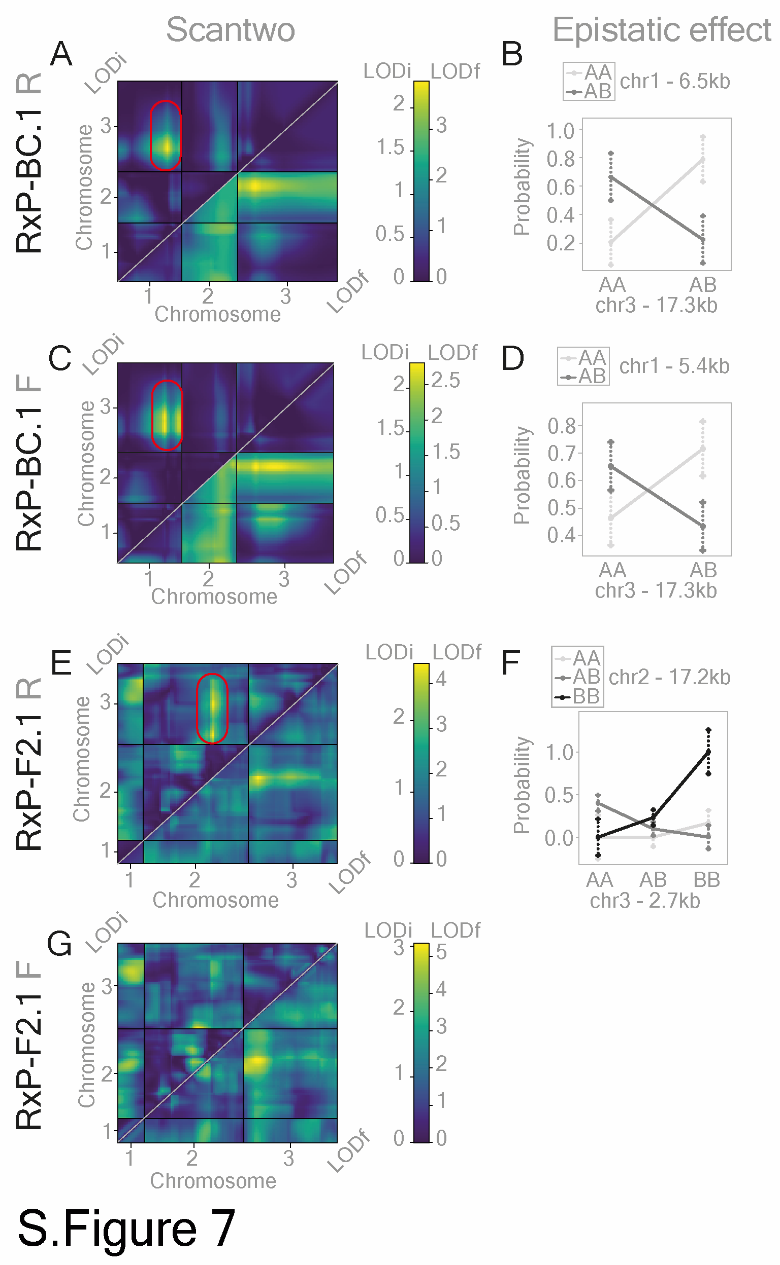

**Supplemental Figure 7. Ros-2FMxPor-1SL QTL mapping: epistatic interactions**

To scan specifically for QTLs in epistasis, we used the *scantwo* function (*rqtl* package). Datasets are labeled (left gray; F = Full and R = Reduced) and correspond to Supplemental Figures 6 and 8. (A, C, E, G): *Scantwo* heatmaps for three chromosomes show LODf in the lower right corner that measures the improvement in the fit of the full two-locus model over the null model and indicates the evidence for at least one QTL with allowance for interaction. LODi heatmap is plotted in the upper left corner and measures the improvement in the fit of the full model over that of the additive model, and so indicates evidence for an interaction. Significant QTL epistatic interaction is marked by a red circle. (B, D, F): The epistatic effect for each of the significant interactions on the left is shown. The marker that shows the strongest interaction on each chromosome was selected and its location is marked in gray letters.

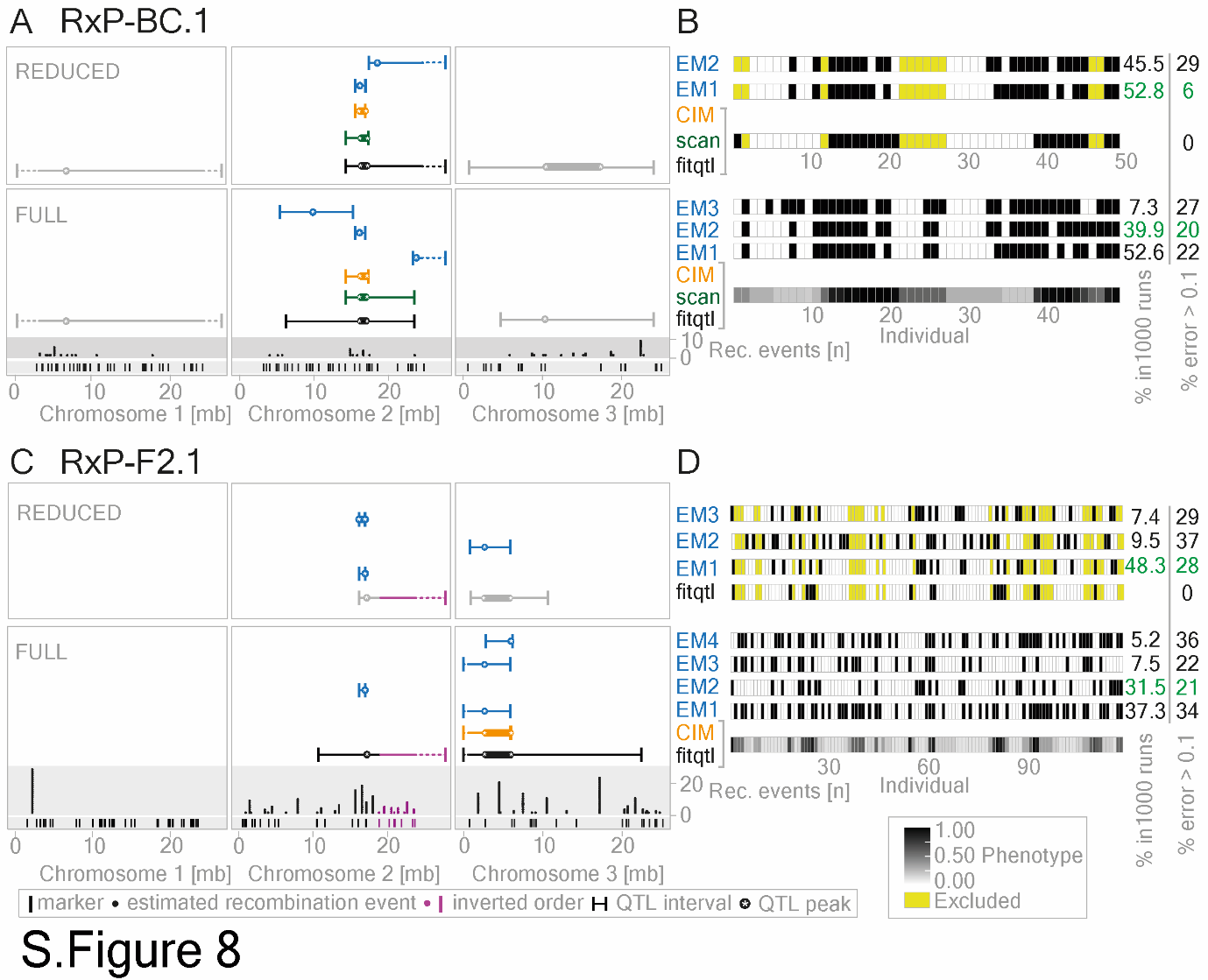

**Supplemental Figure 8. Ros-2FMxPor-1SL QTL mapping: intervals and phenotype scores**

Confidence intervals and phenotype panels for the full QTL analysis (Supplemental Figure 6). (A, C) QTL confidence intervals: composite interval mapping – orange, *scanone* – green, *fitqtl*: additive – black, *fitqtl*: epistatic – gray, EM-algorithm – blue (see all LOD score profiles in Supplemental Figure 6 and the exact coordinates of the markers, recombination events and QTL confidence intervals in Supplemental Table 4). (B, D): Phenotype panels for the corresponding QTL analysis are on the left. The probability of being sensitive (white) or insensitive (black) is shown for each individual. Yellow boxes indicate the individuals that were excluded in the “reduced” dataset due to the probability phenotype between 0.3 and 0.7 (see methods QTL mapping section and Supplemental Figure 3). Numbers on the right indicate for each EM panel how many out of 1000 runs that panel was found, and the fraction of individuals in each panel which had an error > 0.10 from the original data (see methods QTL mapping/EM-pipeline, Supplemental Table 4. The green marks the panel with the highest convergence and lowest error.

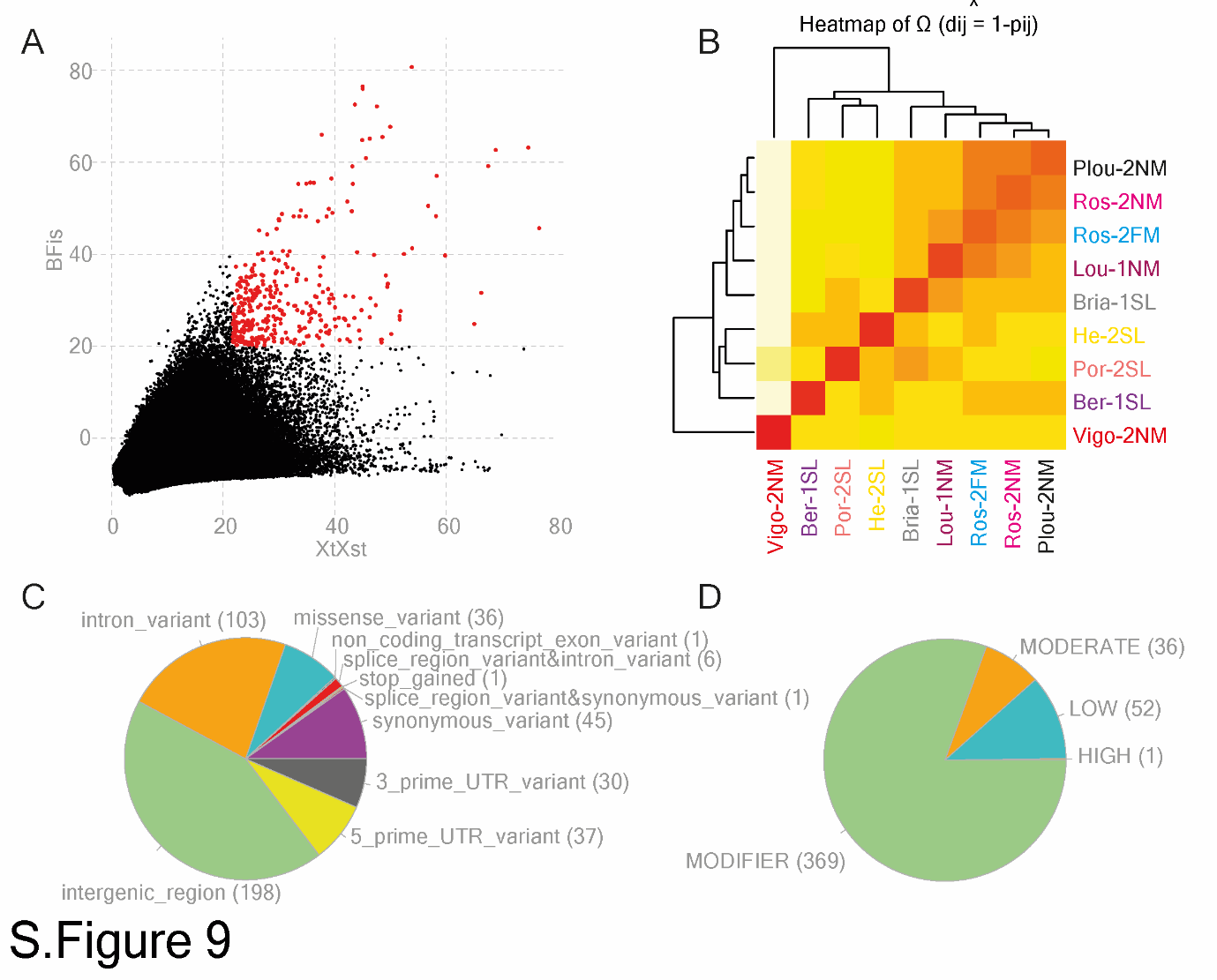

**Supplemental Figure 9. BayPass analysis for Ros-2FM dataset**

(A) The Bayesian factor (BFis) showing the strength of the association is plotted against the differentiation measure (XtXst) for all polymorphic variants analyzed by BayPass. 357 significantly associated SNPs and indels (BFis > 20, eBPis > 2, XtXst > 21.67) are depicted in red. (B) Kinship matrix Ω is given as a heatmap showing reconstructed relationships between nine tested populations. (C-D) The effect of 357 associated variants was analyzed by SNPeff. (C) The effects of the variants on the surrounding genes are depicted in a pie chart. (D) The estimated impact of 357 associated variants is represented in a pie chart.

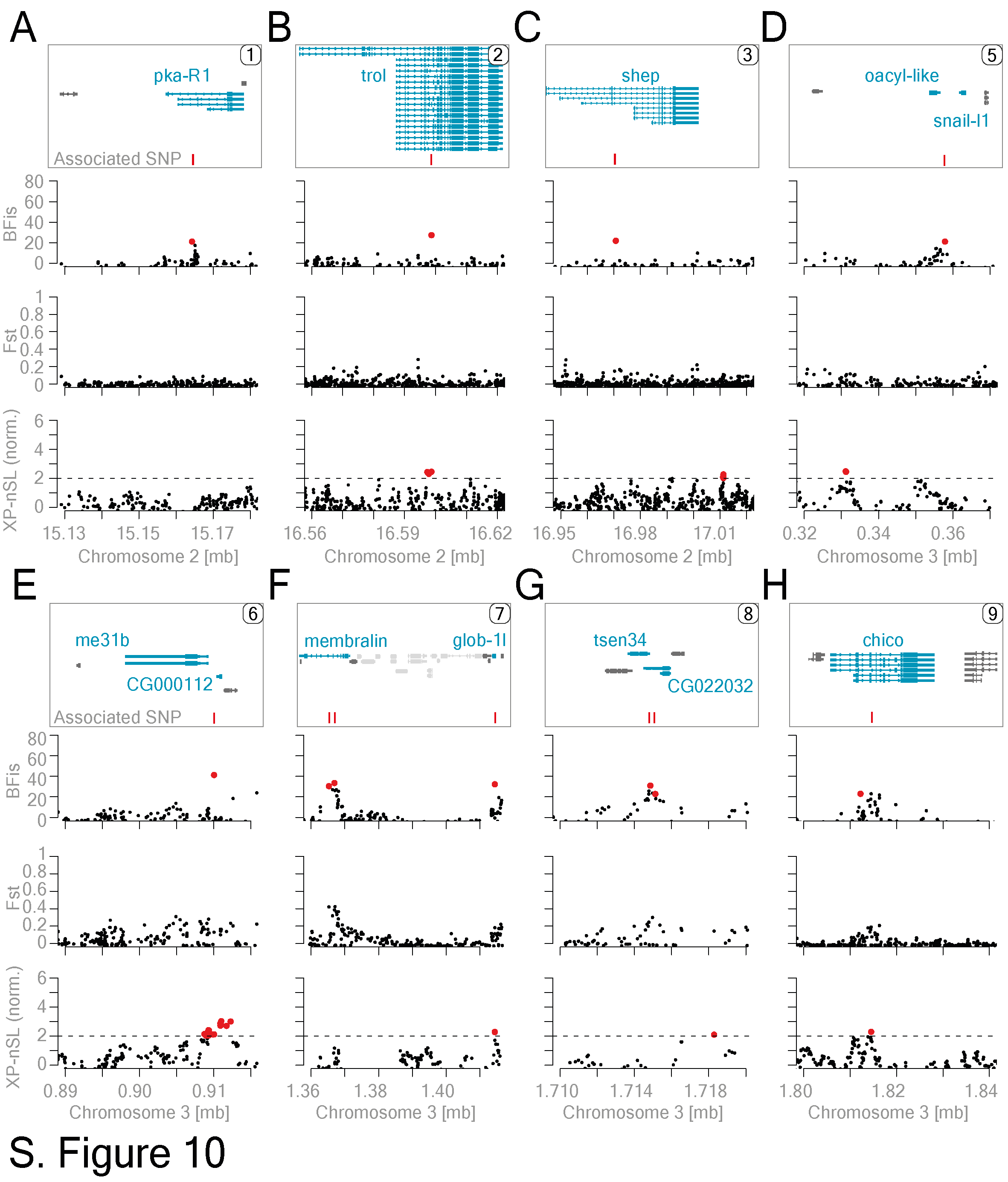

### **Supplemental Figure 10. SNPs associated with the loss of sensitivity to tidal turbulence in Ros-2FM**QTL mapping and association mapping was performed to determine the most likely causative mutations underlying the phenotypic loss in Ros-2FM. Nine loci were identified (Figure 3). Although STAT1 locus was the most likely causal one (Figure 3 C), we investigated all other polymorphisms underlying the QTLs on the second and third chromosomes. A-H panels show for each of these loci: gene affected by the associated SNPs (blue gene models), association score (BFis), genomic differentiation (Fst) between Ros-2FM and Ros-2NM, and selective-sweep analysis (normalized XP-nSL). Phylogenetic trees of all 15 candidate genes (blue gene models) can be seen in Supplemental Figures 11 and 12.

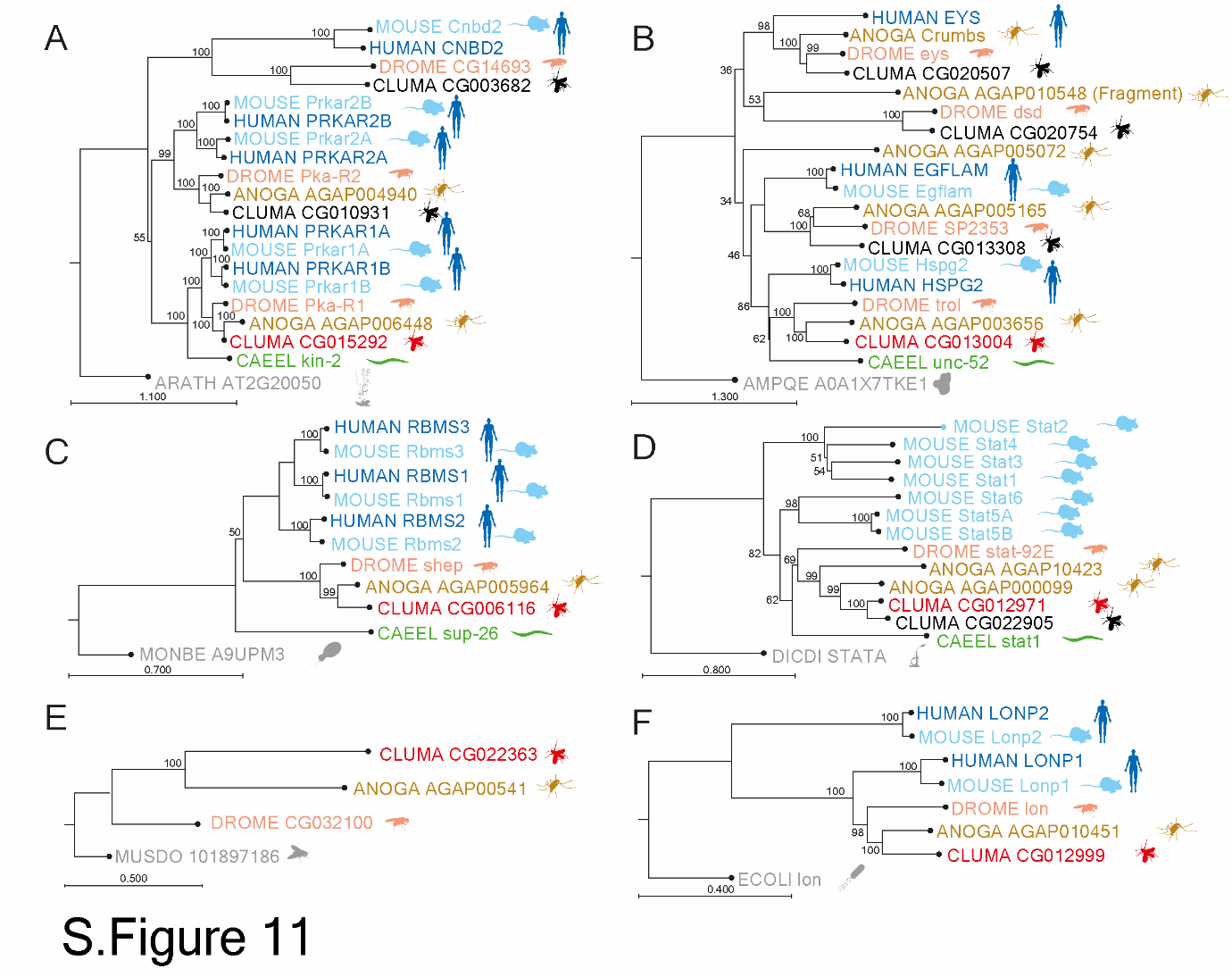

**Supplemental Figure 11. Phylogenetic trees of all the potential candidate genes on the second chromosome.**

Species are color-coded and represented by a pictogram next to the gene names: outgroup – gray, *Caenorhabditis elegans* – green, *Drosophila melanogaster* – orange, *Anopheles gambiae* – brown, *Mus musculus* – light blue*, Homo sapiens* – dark blue, *Clunio marinus* candidate gene – red, *Clunio marinus* other orthologs of the candidate gene – black. Bootstrap values are written above each node. The estimated distance is given below each tree. (A) Protein kinase regulatory subunits. (B) Heparan sulfate proteoglycan Perlecan / Terribly reduced optic lobe. (C) Alan shepard. (D) Signal transducer and transcription activator. (E) Unnamed gravitaxis gene. (F) Lon protease mitochondrial.

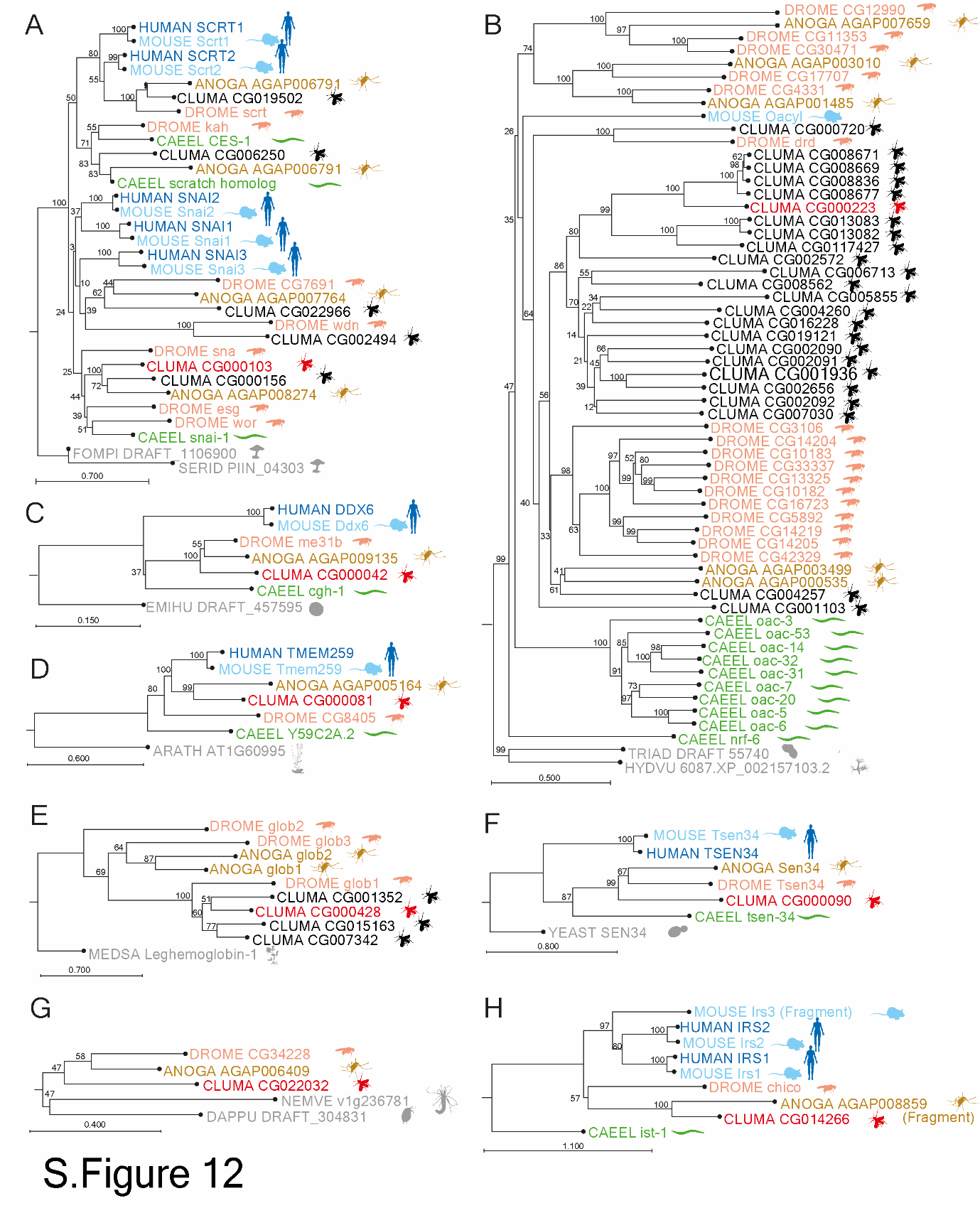

**Supplemental Figure 12. Phylogenetic trees of all the potential candidate genes on the third chromosome.**

Species are color-coded and represented by a pictogram next to the gene names: outgroup – gray, *Caenorhabditis elegans* – green, *Drosophila melanogaster* – orange, *Anopheles gambiae* – brown, *Mus musculus* – light blue*, Homo sapiens* – dark blue, *Clunio marinus* candidate gene – red, *Clunio marinus* other orthologs of the candidate gene – black. Bootstrap values are written above each node. The estimated distance is given below each tree. (A) Snail-like family of transcription factors. (B) O-acyltransferase family. (C) ATP-dependent RNA helicase me31b. (D) Membralin. (E) Globin family. (F) tRNA splicing endonuclease subunit 34 (Tsen34) (G) Unknown protein. (H) Chico, insulin receptor substrate.

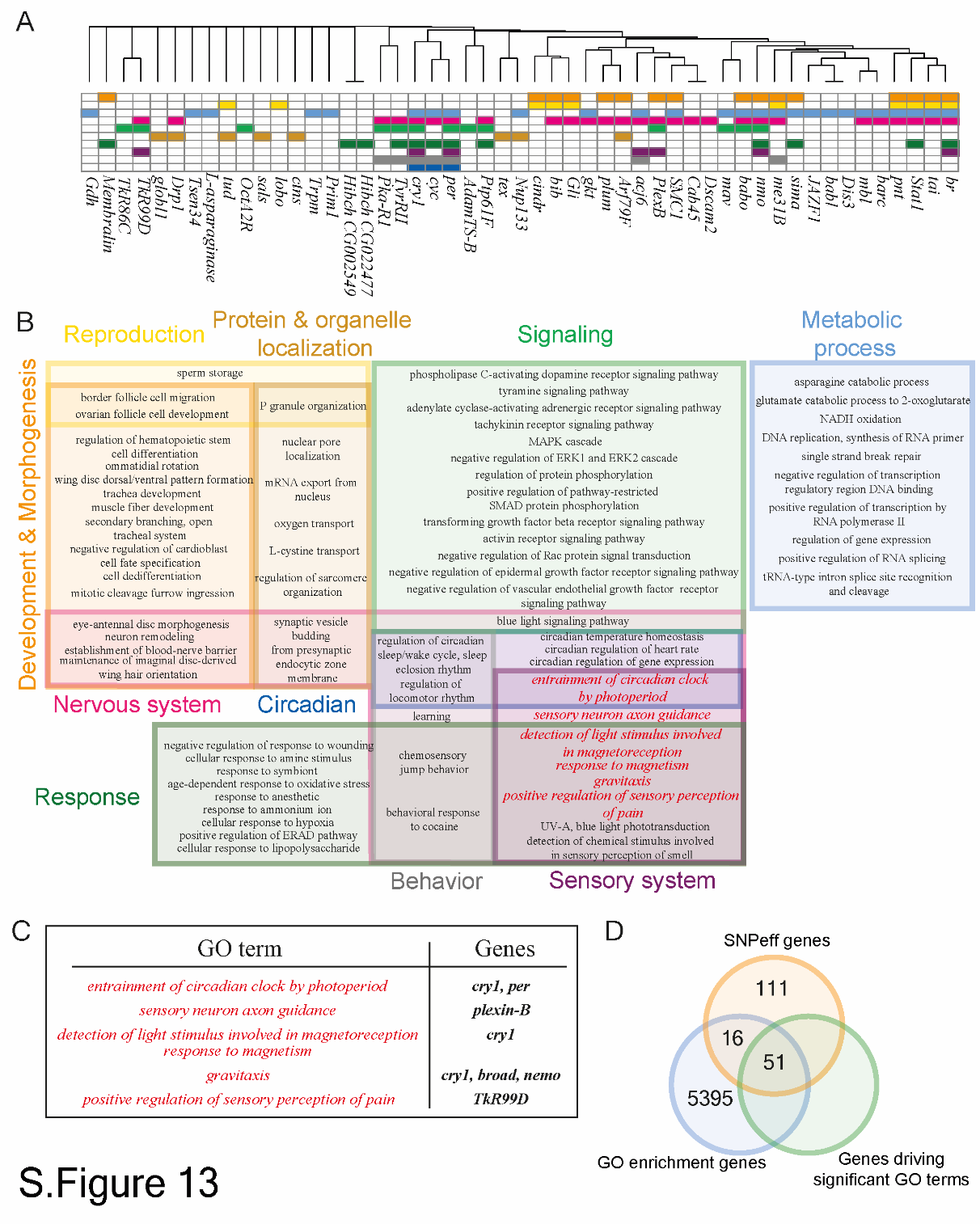

### **Supplemental Figure 13. GO term enrichment of genes associated with the loss of sensitivity to tidal turbulence in Ros-2FM**

BayPass and SNPeff were used to identify 178 genes associated with the loss of sensitivity to tidal turbulence in Ros-2FM. (A) 51 genes that are driving 78 significant GO terms are depicted. Hierarchical clustering of genes and GO terms reveals major clusters of GO terms (color-coded) and listed in panel (B). (C) Several GO terms related to sensory nervous system and potentially involved in mechanosensory entrainment are given in the table together with the corresponding genes. (D) Venn diagram is showing the number of genes that went into the GO term enrichment analysis.

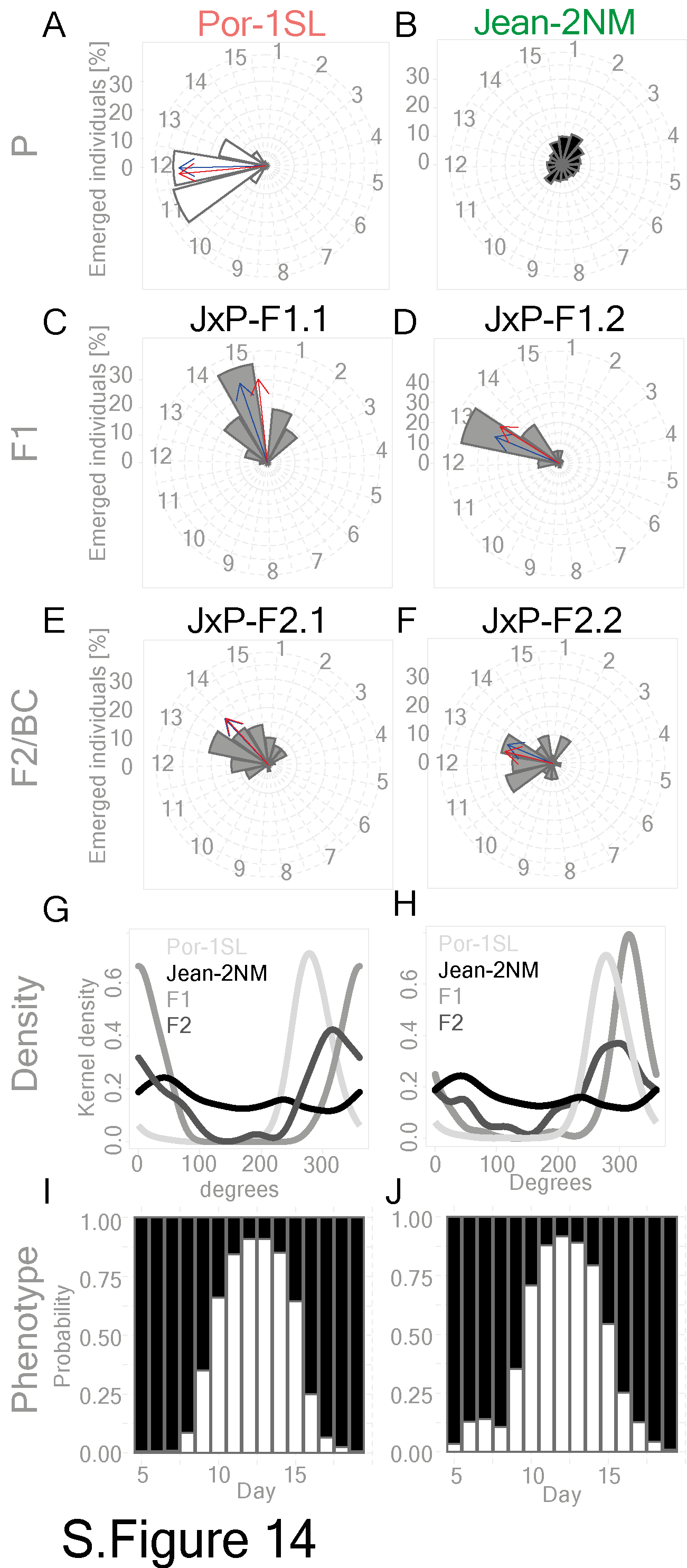

### **Supplemental Figure 14. Calculating probability phenotypes for JxP-F2.1 and JxP-F2.2**

(A-F) The fraction of emerged adults and the mean (red) and median (blue) vectors are plotted. (A) Por-1SL strain. (B) Jean-2NM strain. (C-D) F1 progenies of the two mapping families (E-F) F2 progenies of the two mapping families. (G-H) Kernel density estimates for parental, F1, and F2 generations for each of the two mapping families. Both F2 progenies show a phase shift as compared to the parental Por-1SL strain. (I-J) Bar graphs depict probabilities of finding sensitive - Por-1SL-like (white) or insensitive - Jean-2NM-like (black) individuals on each day in the two crossing families JxP-F2.1 (I) and JxP-F2.2 (J).

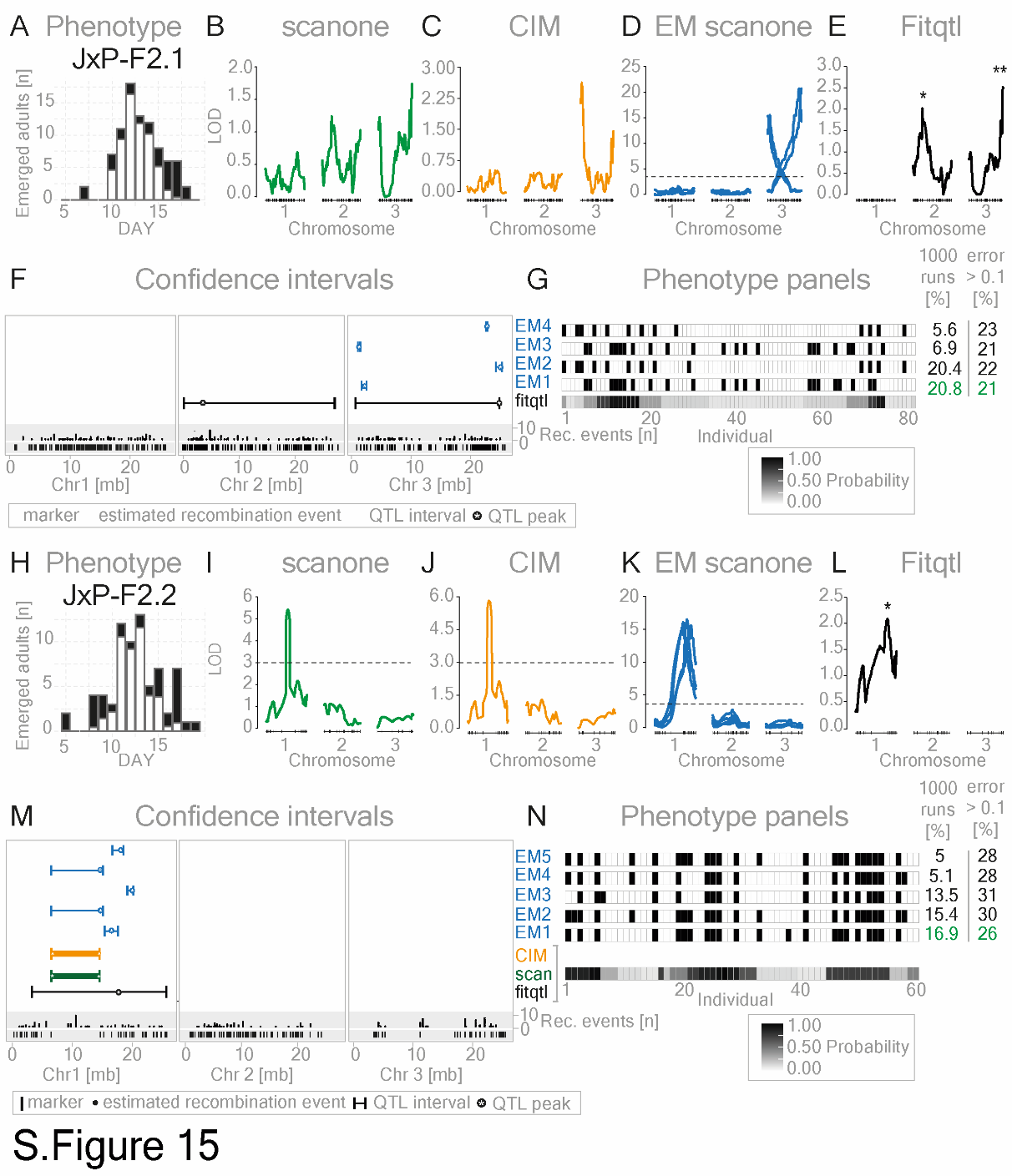

**Supplemental Figure 15. Jean-2NMxPor-1SL QTL mapping full analysis**

Complete QTL mapping results for the two crossing families: JxP-F2.1 and JxP-F2.2.

(A, H) Bar graphs show the number of emerged individuals per day. The predicted ratio of insensitive (black) and sensitive (white) individuals is plotted. (B, I) LOD scores of interval mapping analysis (*scanone*) are designed to detect additive QTLs. The significance threshold (dashed line) was estimated in 1000 permutations with a 5% cutoff. (C, J) CIM analysis with backward regression method, 5 control markers, and a window size of 10 cM. Threshold values are given in Supplemental Table 8. (D, K): LOD scores of *scanone* on EM-optimized binary phenotypes. Results are shown for panels obtained in at least 5% of the cases in 1000 runs (Supplemental Table 8). The significance threshold (dashed line) was estimated in 1000 permutations with a 5% cutoff. (E, L) LOD scores of significant QTLs in multiple QTL mapping pipeline (*fitqtl*). Black lines: additive QTLs, gray lines: QTLs in epistasis. p-value of F statistic is marked: * p-value < 0.05; ** p-value of <0.01. *Fitqlt* statistics are given in Supplemental Table 8. (F, M) Confidence intervals and for the full QTL analysis. QTL confidence intervals: composite interval mapping – orange, *scanone* – green, *fitqtl*: additive – black, *fitqtl*: epistatic – gray, EM-algorithm – blue (exact coordinates of the markers in Supplemental Table 8. (G, N): Phenotype panels for the corresponding QTL analysis. The probability of being sensitive (white) or insensitive (black) is shown for each individual. Numbers on the right indicate for each EM panel how many out of 1000 runs that panel was found, and the fraction of individuals in each panel which had an error > 0.10 from the original data (see methods QTL mapping/EM-pipeline, Supplemental Table 8). The green marks the panel with the highest convergence and lowest error.

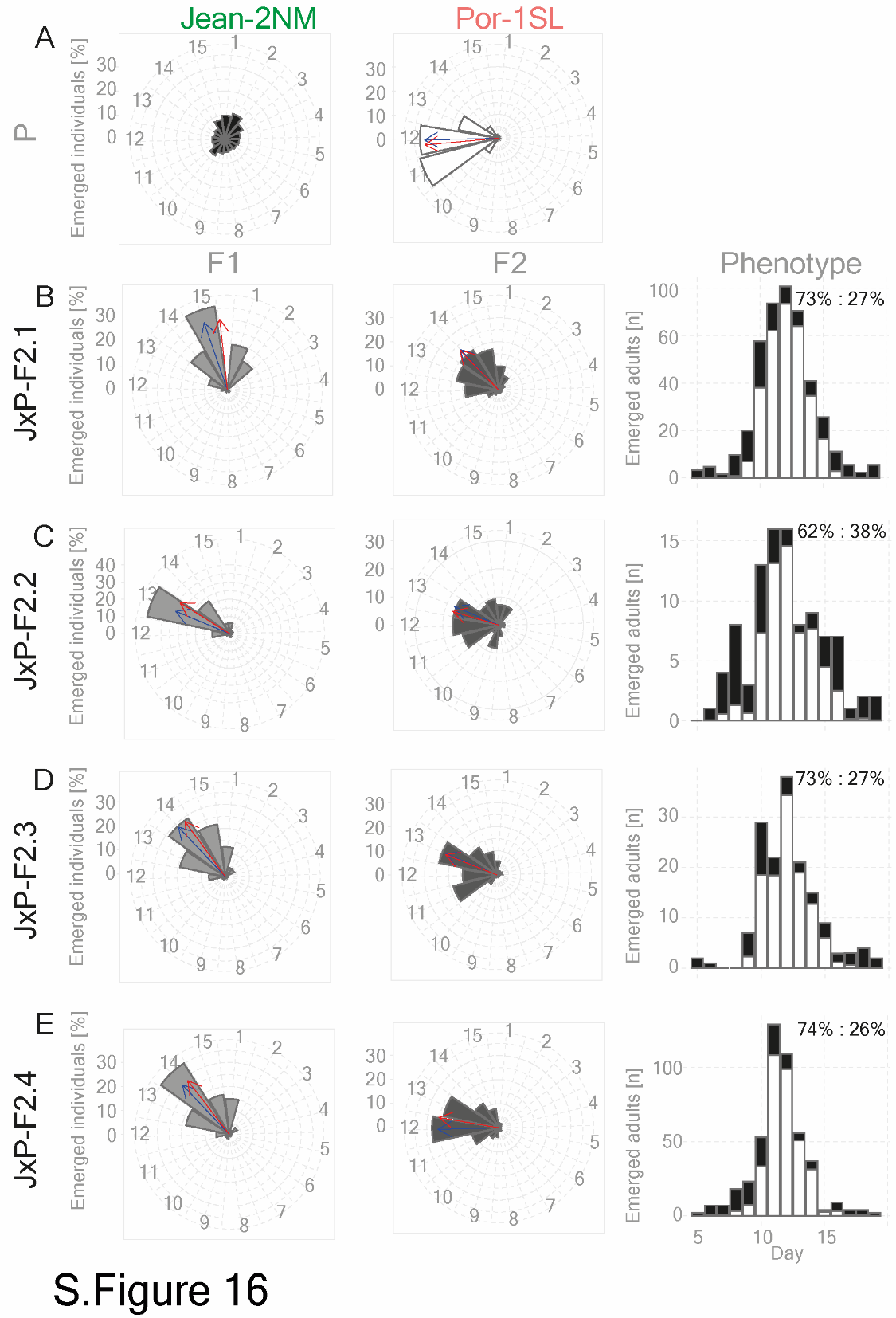

### **Supplemental Figure 16. Probability of finding sensitive and insensitive individuals in several Jean-2NMxPor-1SL intercross families**

(A-E left and middle) The fraction of emerged adults per generation is shown on a circular plot together with the mean and median vector. (A). Jean-2NM parental strain (B) Por-1SL parental strain. (B-E left) Independent F1 progenies of the four intercross families JxP-F2.1-4. (B-E middle) Combined emergence of several F2 families of four intercross families JxP-F2.1-4. Number of individuals is given in Supplemental Table 3. (B-E right) Bar graphs show probabilities of finding sensitive Por-1SL-like (white) or insensitive Jean-2NM-like (black) individuals on each day in the four intercross families.

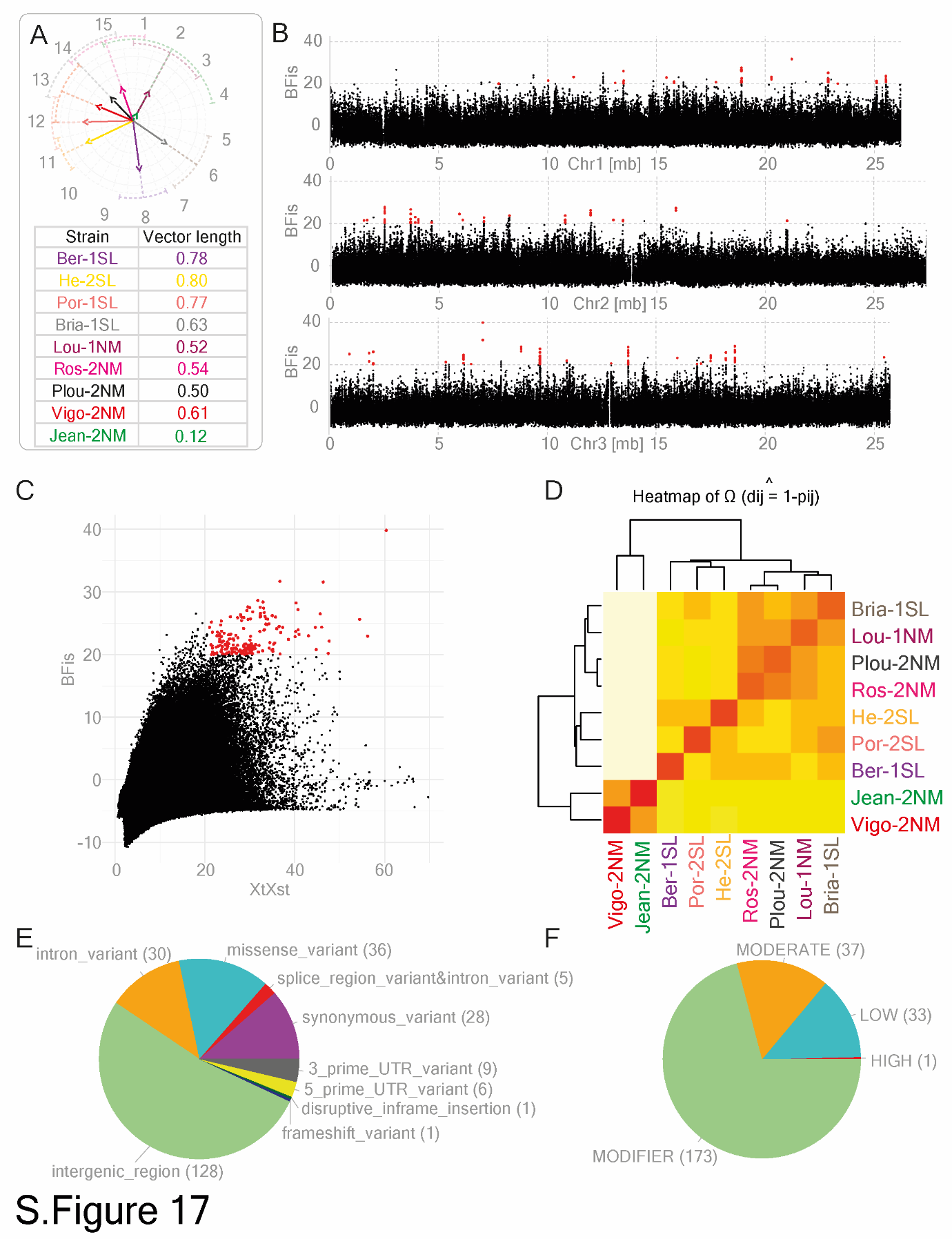

### **Supplemental Figure 17. BayPass analysis for Jean-2NM dataset**

Association analysis was performed to find mutations associated with the loss of sensitivity to tidal turbulence in the Jean-2NM population. (A) Median vector length was used as a proxy for sensitivity to this cue (Supplemental Table 3), (solid lines with arrows; values outside the circle). (B) Association analysis for median vector length with 769.379 SNPs and small indels. Bayesian factor (BFis) is plotted for each variant along the three chromosomes. We found 173 significantly associated SNPs and indels (BFis > 20, eBPis > 2, XtXst > 20.02; see Methods section for details) marked in red. A list of effects and genes affected by mutations is given in Supplemental Table 9.
