## Supplemental Methods for "Genetic analysis of a phenotypic loss in the mechanosensory entrainment of a circalunar clock"

### **Crosses**

Due to the genetically encoded differences in the phase of emergence (Kaiser and Heckel 2012), different strains emerge at different times of the day and month when entrained by the same light-dark (LD) regime. Thus, the three strains had to be kept under different LD and artificial moonlight regimes to synchronize the emergence of adults. Ros-2FM and Por-1SL cultures were kept under the same moonlight entrainment and different LD conditions, while Jean-2NM cultures were kept under alternative moonlight and LD regime. Virgin males and females were caught and placed together in a 10x10x5cm box with seawater and kept overnight to allow for mating and oviposition of eggs. On the next day, adults were collected in 100% ethanol and egg clutches in seawater. Egg clutches with over 30 fertilized eggs were raised individually in 10x10x5cm boxes under tidal turbulence entrainment. Emergence patterns in the F1 generation were recorded each day and adults were stored in 100% ethanol. Adults emerging on the same day freely mated and the resulting F2 offspring were raised in individual boxes. Backcrosses were performed by mating F1 adults with Ros-2FM (or Jean-2NM) adults raised under a moonlight regime. To maximize the chance of getting enough synchronized adults for backcrosses, some F1 cultures were also raised under the moonlight regime (Supplemental Table 3).

### **QTL mapping**: Phenotyping

Emergence data was collected for parental, F1 and F2, and BC generations, and lunar emergence days under turbulence entrainment were assigned as described above (Supplemental Table 3, Supplemental Figures 1 and 2). As expected from previous crossing experiments (Kaiser and Heckel 2012; Kaiser et al. 2016), a considerable phase-shift of the emergence peaks was sometimes found between F1, F2/BC families, and the rhythmic Por-1SL parental strain (Supplemental Table 2, Supplemental Figures 2, 4, 13 and 14). Since the phase is genetically determined (Kaiser et al. 2016; Kaiser and Heckel 2012), we corrected for the phase shift to not mistakenly identify QTLs encoding phase. We first calculated the phase of the peaks in the parental populations as well as F1, F2, and BC families, and if we found a phase difference, we then reassigned the “days under tidal turbulence entrainment” of F2/BC so that the peak days match the Por-1SL peak days (Supplemental Table 2, Supplemental Figures 4 and 14). More specifically, we used the R package “circular” (Pewsey Arthur et al. 2013) and calculated the vector mean direction *mean.circular,* vector median direction *median.circular*, direction length *rho.circular,* sample circular standard deviation *sd.circular,* and angular variance *angular.variance* (Script 1, Supplemental Table 3). Due to the big differences in the total numbers of individuals collected in parental strains (Supplemental Table 1) and F1/F2/BC families (Supplemental Table 3) we did not perform statistical tests designed to assess if the distributions were significant. According to the mean and median directions, peak days in the RxP-BC.1 family remained unchanged while RxP-F2.1, JxP-F2.1, and JxP-F2.2 families were shifted by 2 days (Supplemental Table 2, Supplemental Figures 4, and 14).

In order to phenotype sensitivity to tidal turbulence, we must distinguish between “sensitive” individuals that emerged within the Por-1SL-like peak and “insensitive” that can emerge on any lunar day. A phenotyping problem arises because the emergence peak does not only contain sensitive (rhythmic) individuals, but also some of the insensitive (arrhythmic) individuals. Hence, for individuals found in the emergence peak, we do not know a priori which phenotype they were, while individuals outside the emergence peak are certainly insensitive. To resolve this problem of overlapping phenotypes, we designed a pipeline to calculate the probability of finding “sensitive” and “insensitive” individuals on each day. To get an emergence estimate from raw emergence distributions we calculated kernels density estimates using a *density.circular(bw = 10)* function from R package “circular”. We then calculated the average kernel’s density estimate for each day for parental, F1, F2, and BC distributions. The probabilities of finding sensitive and insensitive individuals on a given day were calculated the following way.

Intercross:

Probability of being insensitive = insensitive parent density / (sensitive parent density + F1 density + insensitive parent density)

Probability of being sensitive = (sensitive parent density + F1 density) / (sensitive parent density + F1 density + insensitive parent density)

Backcross:

Probability of being insensitive = insensitive parent density / (F1 density + insensitive parent density)

Probability of being sensitive = F1 density / (F1 density + insensitive parent density)

We found that sensitivity to turbulence is inherited dominantly. As a result, in the second generation of an intercross family, sensitive individuals have either Por-1SL or F1 genotypes, while insensitive individuals have Ros-2FM or Jean-2NM genotypes. Furthermore, in a back-cross family, the sensitive individuals have F1 genotypes and insensitive individuals have Ros-2FM or Jean-2NM genotypes. The probability of being insensitive was used as a phenotypic score for QTL mapping. In addition, we noticed that on certain days the probability of finding sensitive and insensitive individuals is almost equal. To test how much those individuals influence mapping results, we generated reduced datasets by excluding individuals with probability phenotypes between 0.3 and 0.7 and assigning binary phenotypes to the remaining ones: probability > 0.7 gained phenotype 1, probability < 0.3 gained phenotype 0 (Script 1, Supplemental Figure 3).

### **QTL mapping**: Genotyping

DNA was extracted from adults collected in crossing experiments with the salting-out method (Reineke et al. 1998). Genomic DNA was amplified with standard RepliG protocol (REPLI-g Mini Kit QIAgen 150025). Single-digest RAD sequencing was used to sequence the RxP-BC.1 family (Baird et al. 2008). In brief, 100ng of DNA per sample was digested with BamHI-HF (NEB) at 37°C for 2 h, P1 adapters with BamHI sticky ends (Supplemental Table 10) were ligated with T4 Ligase (NEB) at room temperature for 1h, and heat-inactivated at 65°C for 20 minutes. Samples labelled with unique P1 adapters were pooled and sheared by sonication (Covaris S220 focused-ultrasonicator: Duty cycle = 10; Intensity = 7; Cycles/burst = 300). Following sonication, DNA was precipitated, loaded on the 1% TBE gel, and fragments of the size between 300 and 700bp were extracted from the gel (Zymoclean™ Gel DNA Recovery Kit). Quick Blunting Kit (NEB) was then applied to polish DNA ends and Klenow exo (NEB) to add A-overhangs. Finally, P2 adapters (Supplemental Table 10) were ligated with T4 ligase (NEB), and DNA was amplified with Phusion Master Mix (HF) using P1/P2 amplification primers (Supplemental Table 10) in 12 PCR cycles. Sequencing was performed on Illumina HiSeq3000 with single-end 150 bp reads. To genotype RxP-F2.1, JxP-F2.1, and JxP-F2.2 families, the double-digest RAD sequencing protocol was optimized (Etter et al. 2011; Etter and Johnson 2012). Briefly, 100 ng of DNA per individual was digested with BglII (#FD0084 Thermo Fisher) and MspI (#FD0544 Thermo Fisher) for 2h 37°C. P1 and P2 adapters (Supplemental Table 10) were ligated simultaneously with T4 ligase (#EK0032 Thermo Fisher) for 2 hours at 22°C and heat-inactivated at 70°C for 10min. Samples were pooled, precipitated, and loaded on the 1% TEB gel, and DNA fragments sizes between 200 and 1000bp were extracted (Zymoclean™ Gel DNA Recovery Kit) and amplified with Phusion Master Mix (HF) using P1/P2 amplification primers (Supplemental Table 10) in 12 PCR cycles. Sequencing was performed on Illumina HiSeq3000 with single-end reads for RxP-BC.1 family and paired-end for RxP-F2.1, JxP-F2.1, and JxP-F2.2 families.

### **QTL mapping:** Informative variants and genotype matrix in Jean-2NM x Por-1SL

Samples from parents’ and F1s of the two Jean-2NMxPor-1SL families, unfortunately, had very few good genotypes. Thus, we designed an alternative approach for reconstructing the recombination matrix. RAD sequencing VCF files were filtered for minGQ 20, max-alleles 2, and max-missing 0.60. The few genotypes for which we had parents’ and F1’s genotypes were used as fixed guides (the same coding strategy as in Supplemental Table 11). The remaining markers were kept if they had genotypes in the Jean-2NM and Por-1SL pool-sequencing data generated by sequencing 300 individuals of each strain (Kaiser et al. 2016). We kept markers that in F2 offspring had all 3 genotypes (0/0, 0/1, 1/1) and follow the Hardy–Weinberg principle: 0/0 > 10% or 1/1 > 10% or 0/1 > 40% (Hardy 1908). We then identified recombination events based on genotype-switching along the chromosomes from heterozygous to either of the two homozygous genotypes because we knew that 0/1 is certainly AB but 0/0 and 1/1 could have been either AA or BB depending on the genotypes of the parents (see Supplemental Table 11). We resolved the homozygous genotypes by using parents’ genotypes we had and the consistency genotype assignment in Script 2. Individuals with no genotypes were excluded. Since the two parent strains are fairly divergent, we used the markers’ allele frequency in the natural populations to compare the resolved parent genotypes. As a final quality control, we ran a QTL pipeline using sex as a phenotype since its confidence interval is known (Kaiser et al. 2016). The results were as expected (data not shown), so we are confident that the recombination matrix was correctly reconstructed. The final number of markers was 560 in JxP-F2.1 and 178 in the JxP-F2.2 family. The final genotype matrix is given in Supplemental Table 9.

### **QTL mapping:** Expectation-maximization (EM) algorithm

To explore the effect of uncertainty in phenotyping on the QTL mapping results, we devised an EM algorithm to assign binary phenotypes to the entire dataset (Supplemental Figure 5). We started from calculated insensitivity probabilities, ran the *scanone* function of the R/qtl package (Karl W. Broman and Saunak Sen 2009), and recorded the maximal log odds ratio (LOD) score. We then assigned binary phenotypes to all individuals considering their given insensitivity probabilities. This implies that individuals with probabilities of 0 or 1 could not change. Individuals with an insensitivity probability of say 0.3 would have a 30% chance of being recorded as insensitive (0) and a 70% chance of being recorded as sensitive (1). On the resulting binary phenotype set, we ran *scanone* again and recorded the maximal LOD score. This was repeated 10.000 times and the iteration that gave the highest LOD score overall was kept. In a second step, starting from the best binary phenotype panel we systematically flipped phenotypes of one individual at the time and kept only the single flipped score which increased the maximal LOD score the most. This was iterated until there was no further improvement (maximally 200 times), driving the binary phenotype panel into a local optimum for the LOD score. We ran the entire algorithm with both steps 1000 times from the start and each final optimized phenotype panel was recorded. We then calculated in how many of the 1000 iterations the same optimal panel was found, and how much the optimized binary phenotypes differed from the starting probabilities:

Error per individual in EM panel = ABS (probability of being insensitive – binary phenotype) x ABS (0.5 - the probability of being insensitive)

To assess how much QTL resultsbased on starting probabilities match the resultsfrom EM binary phenotypes we looked at 1) the percentage of convergence in 1000 EM loops and 2) the percentage of individuals in a binary panel with error > 0.10.

### **Selective sweep analysis**

We used selscan 2.0 to calculate XP-nSL (DeGiorgio and Szpiech 2021; Szpiech et al. 2021). Selscan 2.0 assumes that the data is polarized (ancestral and derived state of the alleles is known) and we do not have an outgroup or an ancestral DNA. Thus, we used the major allele in the entire dataset that consists of 15 populations and 349 individuals (20-24 per population) as the ancestral allele. The vcf file containing GATK-called SNPs and indels from 349 individuals was filtered for biallelic sites (leaving 7.134.648 variants) with vcftools version 0.1.14 (Danecek et al. 2011) and then polarized the dataset by setting 0 = major (ancestral) allele and 1 = minor (derived) allele the following way. Vcf files were converted to plink input files with vcftools, allele frequency was calculated for A1 (minor) and A2 (major) alleles in plink, with *--freq* parameter, and a text file was outputted that contains 2 columns: the identity of the major allele and the variant position, which is then used in plink to set the major allele as 0 with *--a1-allele*.

Finally, to search for loci that sweep through the turbulence-insensitive populations due to local adaptation, we contrasted Ros-2FM and Jean-2NM with the closest turbulence-sensitive populations: Ros-2NM and Vigo-2NM respectively. A vcf file consisting of 48 Ros-2FM and Ros-2NM individuals were filtered for minor allele frequency = 0.05, no missing data, and minimal quality (minQ) of 20 leaving 574.132 variants. Similarly, the vcf file containing 47 Jean-2NM and Vigo-2NM males was filtered the same way leaving 697.594 variants.

We calculated XP-nSL per chromosome with *selscan2.0* *--unphased* keeping the default settings: *--cutoff* (the EHH decay cut off) of 0.05; *--gap-scale* (if a gap is encountered between two SNPs the genetic distance is scalled by GAP_SCALE/GAP) was 20.000; *--max-extend* (the maximum distance the EHH decay curve is allowed to extend from the core) was 1.000.000; *--max-gap* (maximum allowed gap in bp between two SNPs) was 200.000. We then ran the *norm* to calculate normalized XP-nSL for all chromosomes together and to identify top candidate regions that contain a significantly high number of core alleles in a given window. We kept default parameters --*crit-val* (iHS based on (Voight et al. 2006)) of 2, *--min-SNPs* (only consider windows with at least this many SNPs) of 10, *--qbins* (the number of quantile bins to use when identifying significant windows binned by a number of sites within each) of 10. We changed the window size *--winsize* from the default value of 100.000 to 10.000 bp because the *Clunio* genome is much smaller than the *Human* genome on which the tool was optimized. In addition, the clusters of SNPs with high association values we identified thus far do not exceed 10kb (See Supplemental Tables 6 and 9).
