## Supplemental Script 1 for "Genetic analysis of a phenotypic loss in the mechanosensory entrainment of a circalunar clock": Script_1_PHENOTYPING_BC_1_calculate_phase_shit_generate_corrected_input_file.R

```
### Generate input file labeled “Input_BC.csv” WITH COLUMNS:
###COL1:TURB_DAY : turbulence day 1-15 	
###COL2:TURB_DAY_ANGLE: turbulence day * 24
###COL3:P1: rhythmic parent 1 emergence data (as they were collected) - not % but number of individuals per day
###COL4:P2: arrhythmic parent 1 emergence data (as they were collected) - not % but number of individuals per day
###COL5:F1: F1 emergence data (as they were collected) - not % but number of individuals per day
###COL6:F2: BC emergence data (as they were collected) - not % but number of individuals per day

### SET PATH TO YOUR INPUT FILE ###


setwd("C:/MY_FILES/Mapping_paper/Figures_data/PHENOTYPING_PIPELINE/BC/")
library(readr)
Input_BC <- read_csv("Input_BC.csv")


################################# 1. Calculate circular summary statistics – check if there is a phase shift ###################################
library(tibble)
library (circular)
library(stringr)
library(ggplot2)
labels <- as.character(1:15)
angles <- c(2*pi/15, 4*pi/15, 6*pi/15, 8*pi/15, 10*pi/15, 12*pi/15, 14*pi/15, 16*pi/15, 18*pi/15, 20*pi/15, 22*pi/15, 24*pi/15, 26*pi/15, 28*pi/15, 30*pi/15)
CIRCULAR_STAT <- c("dirrection_lenght", "circular_sd", "median_dirrection" ,"mean_dirrection", "angular_variance", "angular_deviation", "median_dirrection_positive", "mean_dirrection_positive", "median_dirrection_DAY", "mean_dirrection_DAY")
GROUPS_BC <- c("P1", "P2", "F1", "BC")
CIRCULAR_RESULTS <- matrix(nrow = length(CIRCULAR_STAT), ncol = length(GROUPS_BC))
rownames(CIRCULAR_RESULTS) <- CIRCULAR_STAT 
colnames(CIRCULAR_RESULTS) <- GROUPS_BC

### Calculate circular stats for Parent 1 ###
P1 <- cbind(Input_BC[,2], Input_BC[,3])
DATA <- as.data.frame(P1)
colnames(DATA) <- c("DAY", "NUMBER")
DATA2 <- add_column(DATA, V3 = 0)
DATA2$V3 <- 0
for(i in 1:length(DATA2$DAY)){DATA2[i,3] <- paste(replicate(DATA2[i,2], DATA2[i,1]), collapse = ",")}
DATA3 <- DATA2[DATA2[,2]!=0,3]
DATAforcircular <- paste (DATA3, collapse = ",")
DATAx<- str_split_fixed(string = DATAforcircular, pattern = ",", n=sum(DATA$NUMBER))
DATAx <- as.numeric(DATAx)
P1_C <- circular(DATAx, type="angles", units="degrees")


#direction_length
CIRCULAR_RESULTS[1,1] <- rho.circular(P1_C)
#sample circular standard dev
CIRCULAR_RESULTS[2,1] <- sd.circular(P1_C)
#median dirrection
CIRCULAR_RESULTS[3,1] <- median.circular(P1_C, deviation=TRUE) 
#mean dirrection
CIRCULAR_RESULTS[4,1] <- mean(P1_C)
#angular.variance
CIRCULAR_RESULTS[5,1] <- angular.variance(P1_C)
#angular.deviation
CIRCULAR_RESULTS[6,1] <- angular.deviation(P1_C)
### Calculate circular stats for Parent 2 ###
P2 <- cbind(Input_BC[,2], Input_BC[,4])
DATA <- as.data.frame(P2)
colnames(DATA) <- c("DAY", "NUMBER")
DATA2 <- add_column(DATA, V3 = 0)
DATA2$V3 <- 0
for(i in 1:length(DATA2$DAY)){DATA2[i,3] <- paste(replicate(DATA2[i,2], DATA2[i,1]), collapse = ",")}
DATA3 <- DATA2[DATA2[,2]!=0,3]
DATAforcircular <- paste (DATA3, collapse = ",")
DATAx<- str_split_fixed(string = DATAforcircular, pattern = ",", n=sum(DATA$NUMBER))
DATAx <- as.numeric(DATAx)
P2_C <- circular(DATAx, type="angles", units="degrees")


#direction_length
CIRCULAR_RESULTS[1,2] <- rho.circular(P2_C)
#sample circular standard dev
CIRCULAR_RESULTS[2,2] <- sd.circular(P2_C)
#median dirrection
CIRCULAR_RESULTS[3,2] <- median.circular(P2_C, deviation=TRUE) 
#mean dirrection
CIRCULAR_RESULTS[4,2] <- mean(P2_C)
#angular.variance
CIRCULAR_RESULTS[5,2] <- angular.variance(P2_C)
#angular.deviation
CIRCULAR_RESULTS[6,2] <- angular.deviation(P2_C)
### Calculate circular stats for F1 ###
F1 <- cbind(Input_BC[,2], Input_BC[,5])
DATA <- as.data.frame(F1)
colnames(DATA) <- c("DAY", "NUMBER")
DATA2 <- add_column(DATA, V3 = 0)
DATA2$V3 <- 0
for(i in 1:length(DATA2$DAY)){DATA2[i,3] <- paste(replicate(DATA2[i,2], DATA2[i,1]), collapse = ",")}
DATA3 <- DATA2[DATA2[,2]!=0,3]
DATAforcircular <- paste (DATA3, collapse = ",")
DATAx<- str_split_fixed(string = DATAforcircular, pattern = ",", n=sum(DATA$NUMBER))
DATAx <- as.numeric(DATAx)
F1_C <- circular(DATAx, type="angles", units="degrees")


#direction_length
CIRCULAR_RESULTS[1,3] <- rho.circular(F1_C)
#sample circular standard dev
CIRCULAR_RESULTS[2,3] <- sd.circular(F1_C)
#median dirrection
CIRCULAR_RESULTS[3,3] <- median.circular(F1_C, deviation=TRUE) 
#mean dirrection
CIRCULAR_RESULTS[4,3] <- mean(F1_C)
#angular.variance
CIRCULAR_RESULTS[5,3] <- angular.variance(F1_C)
#angular.deviation
CIRCULAR_RESULTS[6,3] <- angular.deviation(F1_C)
### Calculate circular stats for BC ###
BC <- cbind(Input_BC[,2], Input_BC[,6])
DATA <- as.data.frame(BC)
colnames(DATA) <- c("DAY", "NUMBER")
DATA2 <- add_column(DATA, V3 = 0)
DATA2$V3 <- 0
for(i in 1:length(DATA2$DAY)){DATA2[i,3] <- paste(replicate(DATA2[i,2], DATA2[i,1]), collapse = ",")}
DATA3 <- DATA2[DATA2[,2]!=0,3]
DATAforcircular <- paste (DATA3, collapse = ",")
DATAx<- str_split_fixed(string = DATAforcircular, pattern = ",", n=sum(DATA$NUMBER))
DATAx <- as.numeric(DATAx)
BC_C <- circular(DATAx, type="angles", units="degrees")


#direction_length
CIRCULAR_RESULTS[1,4] <- rho.circular(BC_C)
#sample circular standard dev
CIRCULAR_RESULTS[2,4] <- sd.circular(BC_C)
#median dirrection
CIRCULAR_RESULTS[3,4] <- median.circular(BC_C, deviation=TRUE) 
#mean dirrection
CIRCULAR_RESULTS[4,4] <- mean(BC_C)
#angular.variance
CIRCULAR_RESULTS[5,4] <- angular.variance(BC_C)
#angular.deviation
CIRCULAR_RESULTS[6,4] <- angular.deviation(BC_C)
### Calculate median and mean direction in turbulence days
if (CIRCULAR_RESULTS[3,1]<0){CIRCULAR_RESULTS[7,1] <- 360+CIRCULAR_RESULTS[3,1]} else{CIRCULAR_RESULTS[7,1] <- CIRCULAR_RESULTS[3,1]}
if (CIRCULAR_RESULTS[3,2]<0){CIRCULAR_RESULTS[7,2] <- 360+CIRCULAR_RESULTS[3,2]} else{CIRCULAR_RESULTS[7,2] <- CIRCULAR_RESULTS[3,2]}
if (CIRCULAR_RESULTS[3,3]<0){CIRCULAR_RESULTS[7,3] <- 360+CIRCULAR_RESULTS[3,3]} else{CIRCULAR_RESULTS[7,3] <- CIRCULAR_RESULTS[3,3]}
if (CIRCULAR_RESULTS[3,4]<0){CIRCULAR_RESULTS[7,4] <- 360+CIRCULAR_RESULTS[3,4]} else{CIRCULAR_RESULTS[7,4] <- CIRCULAR_RESULTS[3,4]}
if (CIRCULAR_RESULTS[4,1]<0){CIRCULAR_RESULTS[8,1] <- 360+CIRCULAR_RESULTS[4,1]} else{CIRCULAR_RESULTS[8,1] <- CIRCULAR_RESULTS[4,1]}
if (CIRCULAR_RESULTS[4,2]<0){CIRCULAR_RESULTS[8,2] <- 360+CIRCULAR_RESULTS[4,2]} else{CIRCULAR_RESULTS[8,2] <- CIRCULAR_RESULTS[4,2]}
if (CIRCULAR_RESULTS[4,3]<0){CIRCULAR_RESULTS[8,3] <- 360+CIRCULAR_RESULTS[4,3]} else{CIRCULAR_RESULTS[8,3] <- CIRCULAR_RESULTS[4,3]}
if (CIRCULAR_RESULTS[4,4]<0){CIRCULAR_RESULTS[8,4] <- 360+CIRCULAR_RESULTS[4,4]} else{CIRCULAR_RESULTS[8,4] <- CIRCULAR_RESULTS[4,4]}

CIRCULAR_RESULTS[9,1] <- CIRCULAR_RESULTS[7,1]/24
CIRCULAR_RESULTS[9,2] <- CIRCULAR_RESULTS[7,2]/24
CIRCULAR_RESULTS[9,3] <- CIRCULAR_RESULTS[7,3]/24
CIRCULAR_RESULTS[9,4] <- CIRCULAR_RESULTS[7,4]/24
CIRCULAR_RESULTS[10,1] <- CIRCULAR_RESULTS[8,1]/24
CIRCULAR_RESULTS[10,2] <- CIRCULAR_RESULTS[8,2]/24
CIRCULAR_RESULTS[10,3] <- CIRCULAR_RESULTS[8,3]/24
CIRCULAR_RESULTS[10,4] <- CIRCULAR_RESULTS[8,4]/24

write.csv(CIRCULAR_RESULTS, "Circular_statistics_results_BC.csv")


########################################## 2. If there is a phase-shift generate a corrected phenotype table ####################################
Median_phase_shift_P1_F1 <- CIRCULAR_RESULTS[9,1] - CIRCULAR_RESULTS[9,3]
Median_phase_shift_P1_BC <- CIRCULAR_RESULTS[9,1] - CIRCULAR_RESULTS[9,4]
Mean_phase_shift_P1_F1 <- CIRCULAR_RESULTS[10,1] - CIRCULAR_RESULTS[10,3]
Mean_phase_shift_P1_BC <- CIRCULAR_RESULTS[10,1] - CIRCULAR_RESULTS[10,4]
Input_BC_PHASE_CORRECTED <- Input_BC
Input_BC_PHASE_CORRECTED$F1_COR_DAY <- Input_BC_PHASE_CORRECTED$TURB_DAY - round(Median_phase_shift_P1_F1, digits = 0)
Input_BC_PHASE_CORRECTED$F1_COR_DAY[Input_BC_PHASE_CORRECTED$F1_COR_DAY %in% "0"] <- "15"
Input_BC_PHASE_CORRECTED$F1_COR_DAY[Input_BC_PHASE_CORRECTED$F1_COR_DAY %in% "-1"] <- "14"
Input_BC_PHASE_CORRECTED$F1_COR_DAY[Input_BC_PHASE_CORRECTED$F1_COR_DAY %in% "-2"] <- "13"
Input_BC_PHASE_CORRECTED$F1_COR_DAY[Input_BC_PHASE_CORRECTED$F1_COR_DAY %in% "-3"] <- "12"
Input_BC_PHASE_CORRECTED$F1_COR_DAY[Input_BC_PHASE_CORRECTED$F1_COR_DAY %in% "-4"] <- "11"
Input_BC_PHASE_CORRECTED$F1_COR_DAY[Input_BC_PHASE_CORRECTED$F1_COR_DAY %in% "-5"] <- "10"
Input_BC_PHASE_CORRECTED$F1_COR_DAY[Input_BC_PHASE_CORRECTED$F1_COR_DAY %in% "-6"] <- "9"
Input_BC_PHASE_CORRECTED$F1_COR_DAY[Input_BC_PHASE_CORRECTED$F1_COR_DAY %in% "-7"] <- "8"
Input_BC_PHASE_CORRECTED$F1_COR_DAY[Input_BC_PHASE_CORRECTED$F1_COR_DAY %in% "-8"] <- "7"
Input_BC_PHASE_CORRECTED$F1_COR_DAY[Input_BC_PHASE_CORRECTED$F1_COR_DAY %in% "-9"] <- "6"
Input_BC_PHASE_CORRECTED$F1_COR_DAY[Input_BC_PHASE_CORRECTED$F1_COR_DAY %in% "-10"] <- "5"
Input_BC_PHASE_CORRECTED$F1_COR_DAY[Input_BC_PHASE_CORRECTED$F1_COR_DAY %in% "-11"] <- "4"
Input_BC_PHASE_CORRECTED$F1_COR_DAY[Input_BC_PHASE_CORRECTED$F1_COR_DAY %in% "-12"] <- "3"
Input_BC_PHASE_CORRECTED$F1_COR_DAY[Input_BC_PHASE_CORRECTED$F1_COR_DAY %in% "-13"] <- "2"
Input_BC_PHASE_CORRECTED$F1_COR_DAY[Input_BC_PHASE_CORRECTED$F1_COR_DAY %in% "-14"] <- "1"


Input_BC_PHASE_CORRECTED$BC_COR_DAY <- Input_BC_PHASE_CORRECTED$TURB_DAY - round(Median_phase_shift_P1_BC, digits = 0)
Input_BC_PHASE_CORRECTED$BC_COR_DAY[Input_BC_PHASE_CORRECTED$BC_COR_DAY %in% "0"] <- "15"
Input_BC_PHASE_CORRECTED$BC_COR_DAY[Input_BC_PHASE_CORRECTED$BC_COR_DAY %in% "-1"] <- "14"
Input_BC_PHASE_CORRECTED$BC_COR_DAY[Input_BC_PHASE_CORRECTED$BC_COR_DAY %in% "-2"] <- "13"
Input_BC_PHASE_CORRECTED$BC_COR_DAY[Input_BC_PHASE_CORRECTED$BC_COR_DAY %in% "-3"] <- "12"
Input_BC_PHASE_CORRECTED$BC_COR_DAY[Input_BC_PHASE_CORRECTED$BC_COR_DAY %in% "-4"] <- "11"
Input_BC_PHASE_CORRECTED$BC_COR_DAY[Input_BC_PHASE_CORRECTED$BC_COR_DAY %in% "-5"] <- "10"
Input_BC_PHASE_CORRECTED$BC_COR_DAY[Input_BC_PHASE_CORRECTED$BC_COR_DAY %in% "-6"] <- "9"
Input_BC_PHASE_CORRECTED$BC_COR_DAY[Input_BC_PHASE_CORRECTED$BC_COR_DAY %in% "-7"] <- "8"
Input_BC_PHASE_CORRECTED$BC_COR_DAY[Input_BC_PHASE_CORRECTED$BC_COR_DAY %in% "-8"] <- "7"
Input_BC_PHASE_CORRECTED$BC_COR_DAY[Input_BC_PHASE_CORRECTED$BC_COR_DAY %in% "-9"] <- "6"
Input_BC_PHASE_CORRECTED$BC_COR_DAY[Input_BC_PHASE_CORRECTED$BC_COR_DAY %in% "-10"] <- "5"
Input_BC_PHASE_CORRECTED$BC_COR_DAY[Input_BC_PHASE_CORRECTED$BC_COR_DAY %in% "-11"] <- "4"
Input_BC_PHASE_CORRECTED$BC_COR_DAY[Input_BC_PHASE_CORRECTED$BC_COR_DAY %in% "-12"] <- "3"
Input_BC_PHASE_CORRECTED$BC_COR_DAY[Input_BC_PHASE_CORRECTED$BC_COR_DAY %in% "-13"] <- "2"
Input_BC_PHASE_CORRECTED$BC_COR_DAY[Input_BC_PHASE_CORRECTED$BC_COR_DAY %in% "-14"] <- "1"

Input_BC_PHASE_CORRECTED$F1_COR_EMERGENCE <- 0
Input_BC_PHASE_CORRECTED$BC_COR_EMERGENCE <- 0
Input_BC_PHASE_CORRECTED <- as.matrix(Input_BC_PHASE_CORRECTED)

Input_BC_PHASE_CORRECTED[1,9] <- Input_BC_PHASE_CORRECTED[as.numeric(Input_BC_PHASE_CORRECTED[1,7]),5]
Input_BC_PHASE_CORRECTED[2,9] <- Input_BC_PHASE_CORRECTED[as.numeric(Input_BC_PHASE_CORRECTED[2,7]),5]
Input_BC_PHASE_CORRECTED[3,9] <- Input_BC_PHASE_CORRECTED[as.numeric(Input_BC_PHASE_CORRECTED[3,7]),5]
Input_BC_PHASE_CORRECTED[4,9] <- Input_BC_PHASE_CORRECTED[as.numeric(Input_BC_PHASE_CORRECTED[4,7]),5]
Input_BC_PHASE_CORRECTED[5,9] <- Input_BC_PHASE_CORRECTED[as.numeric(Input_BC_PHASE_CORRECTED[5,7]),5]
Input_BC_PHASE_CORRECTED[6,9] <- Input_BC_PHASE_CORRECTED[as.numeric(Input_BC_PHASE_CORRECTED[6,7]),5]
Input_BC_PHASE_CORRECTED[7,9] <- Input_BC_PHASE_CORRECTED[as.numeric(Input_BC_PHASE_CORRECTED[7,7]),5]
Input_BC_PHASE_CORRECTED[8,9] <- Input_BC_PHASE_CORRECTED[as.numeric(Input_BC_PHASE_CORRECTED[8,7]),5]
Input_BC_PHASE_CORRECTED[9,9] <- Input_BC_PHASE_CORRECTED[as.numeric(Input_BC_PHASE_CORRECTED[9,7]),5]
Input_BC_PHASE_CORRECTED[10,9] <- Input_BC_PHASE_CORRECTED[as.numeric(Input_BC_PHASE_CORRECTED[10,7]),5]
Input_BC_PHASE_CORRECTED[11,9] <- Input_BC_PHASE_CORRECTED[as.numeric(Input_BC_PHASE_CORRECTED[11,7]),5]
Input_BC_PHASE_CORRECTED[12,9] <- Input_BC_PHASE_CORRECTED[as.numeric(Input_BC_PHASE_CORRECTED[12,7]),5]
Input_BC_PHASE_CORRECTED[13,9] <- Input_BC_PHASE_CORRECTED[as.numeric(Input_BC_PHASE_CORRECTED[13,7]),5]
Input_BC_PHASE_CORRECTED[14,9] <- Input_BC_PHASE_CORRECTED[as.numeric(Input_BC_PHASE_CORRECTED[14,7]),5]
Input_BC_PHASE_CORRECTED[15,9] <- Input_BC_PHASE_CORRECTED[as.numeric(Input_BC_PHASE_CORRECTED[15,7]),5]
Input_BC_PHASE_CORRECTED[1,10] <- Input_BC_PHASE_CORRECTED[as.numeric(Input_BC_PHASE_CORRECTED[1,8]),6]
Input_BC_PHASE_CORRECTED[2,10] <- Input_BC_PHASE_CORRECTED[as.numeric(Input_BC_PHASE_CORRECTED[2,8]),6]
Input_BC_PHASE_CORRECTED[3,10] <- Input_BC_PHASE_CORRECTED[as.numeric(Input_BC_PHASE_CORRECTED[3,8]),6]
Input_BC_PHASE_CORRECTED[4,10] <- Input_BC_PHASE_CORRECTED[as.numeric(Input_BC_PHASE_CORRECTED[4,8]),6]
Input_BC_PHASE_CORRECTED[5,10] <- Input_BC_PHASE_CORRECTED[as.numeric(Input_BC_PHASE_CORRECTED[5,8]),6]
Input_BC_PHASE_CORRECTED[6,10] <- Input_BC_PHASE_CORRECTED[as.numeric(Input_BC_PHASE_CORRECTED[6,8]),6]
Input_BC_PHASE_CORRECTED[7,10] <- Input_BC_PHASE_CORRECTED[as.numeric(Input_BC_PHASE_CORRECTED[7,8]),6]
Input_BC_PHASE_CORRECTED[8,10] <- Input_BC_PHASE_CORRECTED[as.numeric(Input_BC_PHASE_CORRECTED[8,8]),6]
Input_BC_PHASE_CORRECTED[9,10] <- Input_BC_PHASE_CORRECTED[as.numeric(Input_BC_PHASE_CORRECTED[9,8]),6]
Input_BC_PHASE_CORRECTED[10,10] <- Input_BC_PHASE_CORRECTED[as.numeric(Input_BC_PHASE_CORRECTED[10,8]),6]
Input_BC_PHASE_CORRECTED[11,10] <- Input_BC_PHASE_CORRECTED[as.numeric(Input_BC_PHASE_CORRECTED[11,8]),6]
Input_BC_PHASE_CORRECTED[12,10] <- Input_BC_PHASE_CORRECTED[as.numeric(Input_BC_PHASE_CORRECTED[12,8]),6]
Input_BC_PHASE_CORRECTED[13,10] <- Input_BC_PHASE_CORRECTED[as.numeric(Input_BC_PHASE_CORRECTED[13,8]),6]
Input_BC_PHASE_CORRECTED[14,10] <- Input_BC_PHASE_CORRECTED[as.numeric(Input_BC_PHASE_CORRECTED[14,8]),6]
Input_BC_PHASE_CORRECTED[15,10] <- Input_BC_PHASE_CORRECTED[as.numeric(Input_BC_PHASE_CORRECTED[15,8]),6]

write.csv(Input_BC_PHASE_CORRECTED, "Input_BC_PHASE_CORRECTED_FULL.csv", row.names = F) 
Input_BC_matrix <- as.matrix(Input_BC)
Input_BC_PHASE_SHIFT_COR <- cbind(Input_BC_matrix[,1:4], Input_BC_PHASE_CORRECTED[,9:10])
write.csv(Input_BC_PHASE_SHIFT_COR, "Input_BC_PHASE_SHIFT_COR.csv", row.names = F)
```
