## Supplemental Script 1 for "Genetic analysis of a phenotypic loss in the mechanosensory entrainment of a circalunar clock": Script_1_PHENOTYPING_BC_2_calculate_density_estimates_generate_prob.R

```
################################## 3. Generate kernels density estimates ###########################################
setwd("C:/MY_FILES/Mapping_paper/Figures_data/PHENOTYPING_PIPELINE/BC/")
library(readr)
Input_BC <- read_csv("Input_BC_PHASE_SHIFT_COR.csv")
############################# Generate circular files for P1, P2, F1, F2#######################################
library(tibble)
library (circular)
library(stringr)
library(ggplot2)
labels <- as.character(1:15)
angles <- c(2*pi/15, 4*pi/15, 6*pi/15, 8*pi/15, 10*pi/15, 12*pi/15, 14*pi/15, 16*pi/15, 18*pi/15, 20*pi/15, 22*pi/15, 24*pi/15, 26*pi/15, 28*pi/15, 30*pi/15)
CIRCULAR_STAT <- c("dirrection_lenght", "circular_sd", "median_dirrection" ,"mean_dirrection", "angular_variance", "angular_deviation", "median_dirrection_positive", "mean_dirrection_positive", "median_dirrection_DAY", "mean_dirrection_DAY")
GROUPS_BC <- c("P1", "P2", "F1", "BC")
CIRCULAR_RESULTS <- matrix(nrow = length(CIRCULAR_STAT), ncol = length(GROUPS_BC))
rownames(CIRCULAR_RESULTS) <- CIRCULAR_STAT 
colnames(CIRCULAR_RESULTS) <- GROUPS_BC


### Calculate circular stats for Parent 1 ###
P1 <- cbind(Input_BC[,2], Input_BC[,3])
DATA <- as.data.frame(P1)
colnames(DATA) <- c("DAY", "NUMBER")
DATA2 <- add_column(DATA, V3 = 0)
DATA2$V3 <- 0
for(i in 1:length(DATA2$DAY)){DATA2[i,3] <- paste(replicate(DATA2[i,2], DATA2[i,1]), collapse = ",")}
DATA3 <- DATA2[DATA2[,2]!=0,3]
DATAforcircular <- paste (DATA3, collapse = ",")
DATAx<- str_split_fixed(string = DATAforcircular, pattern = ",", n=sum(DATA$NUMBER))
DATAx <- as.numeric(DATAx)
P1_C <- circular(DATAx, type="angles", units="degrees")


#direction_length
CIRCULAR_RESULTS[1,1] <- rho.circular(P1_C)
#sample circular standard dev
CIRCULAR_RESULTS[2,1] <- sd.circular(P1_C)
#median dirrection
CIRCULAR_RESULTS[3,1] <- median.circular(P1_C, deviation=TRUE) 
#mean dirrection
CIRCULAR_RESULTS[4,1] <- mean(P1_C)
#angular.variance
CIRCULAR_RESULTS[5,1] <- angular.variance(P1_C)
#angular.deviation
CIRCULAR_RESULTS[6,1] <- angular.deviation(P1_C)
### Calculate circular stats for Parent 2 ###
P2 <- cbind(Input_BC[,2], Input_BC[,4])
DATA <- as.data.frame(P2)
colnames(DATA) <- c("DAY", "NUMBER")
DATA2 <- add_column(DATA, V3 = 0)
DATA2$V3 <- 0
for(i in 1:length(DATA2$DAY)){DATA2[i,3] <- paste(replicate(DATA2[i,2], DATA2[i,1]), collapse = ",")}
DATA3 <- DATA2[DATA2[,2]!=0,3]
DATAforcircular <- paste (DATA3, collapse = ",")
DATAx<- str_split_fixed(string = DATAforcircular, pattern = ",", n=sum(DATA$NUMBER))
DATAx <- as.numeric(DATAx)
P2_C <- circular(DATAx, type="angles", units="degrees")


#direction_length
CIRCULAR_RESULTS[1,2] <- rho.circular(P2_C)
#sample circular standard dev
CIRCULAR_RESULTS[2,2] <- sd.circular(P2_C)
#median dirrection
CIRCULAR_RESULTS[3,2] <- median.circular(P2_C, deviation=TRUE) 
#mean dirrection
CIRCULAR_RESULTS[4,2] <- mean(P2_C)
#angular.variance
CIRCULAR_RESULTS[5,2] <- angular.variance(P2_C)
#angular.deviation
CIRCULAR_RESULTS[6,2] <- angular.deviation(P2_C)
### Calculate circular stats for F1 ###
F1 <- cbind(Input_BC[,2], Input_BC[,5])
DATA <- as.data.frame(F1)
colnames(DATA) <- c("DAY", "NUMBER")
DATA2 <- add_column(DATA, V3 = 0)
DATA2$V3 <- 0
for(i in 1:length(DATA2$DAY)){DATA2[i,3] <- paste(replicate(DATA2[i,2], DATA2[i,1]), collapse = ",")}
DATA3 <- DATA2[DATA2[,2]!=0,3]
DATAforcircular <- paste (DATA3, collapse = ",")
DATAx<- str_split_fixed(string = DATAforcircular, pattern = ",", n=sum(DATA$NUMBER))
DATAx <- as.numeric(DATAx)
F1_C <- circular(DATAx, type="angles", units="degrees")


#direction_length
CIRCULAR_RESULTS[1,3] <- rho.circular(F1_C)
#sample circular standard dev
CIRCULAR_RESULTS[2,3] <- sd.circular(F1_C)
#median dirrection
CIRCULAR_RESULTS[3,3] <- median.circular(F1_C, deviation=TRUE) 
#mean dirrection
CIRCULAR_RESULTS[4,3] <- mean(F1_C)
#angular.variance
CIRCULAR_RESULTS[5,3] <- angular.variance(F1_C)
#angular.deviation
CIRCULAR_RESULTS[6,3] <- angular.deviation(F1_C)
### Calculate circular stats for BC ###
BC <- cbind(Input_BC[,2], Input_BC[,6])
DATA <- as.data.frame(BC)
colnames(DATA) <- c("DAY", "NUMBER")
DATA2 <- add_column(DATA, V3 = 0)
DATA2$V3 <- 0
for(i in 1:length(DATA2$DAY)){DATA2[i,3] <- paste(replicate(DATA2[i,2], DATA2[i,1]), collapse = ",")}
DATA3 <- DATA2[DATA2[,2]!=0,3]
DATAforcircular <- paste (DATA3, collapse = ",")
DATAx<- str_split_fixed(string = DATAforcircular, pattern = ",", n=sum(DATA$NUMBER))
DATAx <- as.numeric(DATAx)
BC_C <- circular(DATAx, type="angles", units="degrees")


#direction_length
CIRCULAR_RESULTS[1,4] <- rho.circular(BC_C)
#sample circular standard dev
CIRCULAR_RESULTS[2,4] <- sd.circular(BC_C)
#median dirrection
CIRCULAR_RESULTS[3,4] <- median.circular(BC_C, deviation=TRUE) 
#mean dirrection
CIRCULAR_RESULTS[4,4] <- mean(BC_C)
#angular.variance
CIRCULAR_RESULTS[5,4] <- angular.variance(BC_C)
#angular.deviation
CIRCULAR_RESULTS[6,4] <- angular.deviation(BC_C)

### Calculate median and mean direction in turbulence days
if (CIRCULAR_RESULTS[3,1]<0){CIRCULAR_RESULTS[7,1] <- 360+CIRCULAR_RESULTS[3,1]} else{CIRCULAR_RESULTS[7,1] <- CIRCULAR_RESULTS[3,1]}
if (CIRCULAR_RESULTS[3,2]<0){CIRCULAR_RESULTS[7,2] <- 360+CIRCULAR_RESULTS[3,2]} else{CIRCULAR_RESULTS[7,2] <- CIRCULAR_RESULTS[3,2]}
if (CIRCULAR_RESULTS[3,3]<0){CIRCULAR_RESULTS[7,3] <- 360+CIRCULAR_RESULTS[3,3]} else{CIRCULAR_RESULTS[7,3] <- CIRCULAR_RESULTS[3,3]}
if (CIRCULAR_RESULTS[3,4]<0){CIRCULAR_RESULTS[7,4] <- 360+CIRCULAR_RESULTS[3,4]} else{CIRCULAR_RESULTS[7,4] <- CIRCULAR_RESULTS[3,4]}
if (CIRCULAR_RESULTS[4,1]<0){CIRCULAR_RESULTS[8,1] <- 360+CIRCULAR_RESULTS[4,1]} else{CIRCULAR_RESULTS[8,1] <- CIRCULAR_RESULTS[4,1]}
if (CIRCULAR_RESULTS[4,2]<0){CIRCULAR_RESULTS[8,2] <- 360+CIRCULAR_RESULTS[4,2]} else{CIRCULAR_RESULTS[8,2] <- CIRCULAR_RESULTS[4,2]}
if (CIRCULAR_RESULTS[4,3]<0){CIRCULAR_RESULTS[8,3] <- 360+CIRCULAR_RESULTS[4,3]} else{CIRCULAR_RESULTS[8,3] <- CIRCULAR_RESULTS[4,3]}
if (CIRCULAR_RESULTS[4,4]<0){CIRCULAR_RESULTS[8,4] <- 360+CIRCULAR_RESULTS[4,4]} else{CIRCULAR_RESULTS[8,4] <- CIRCULAR_RESULTS[4,4]}

CIRCULAR_RESULTS[9,1] <- CIRCULAR_RESULTS[7,1]/24
CIRCULAR_RESULTS[9,2] <- CIRCULAR_RESULTS[7,2]/24
CIRCULAR_RESULTS[9,3] <- CIRCULAR_RESULTS[7,3]/24
CIRCULAR_RESULTS[9,4] <- CIRCULAR_RESULTS[7,4]/24
CIRCULAR_RESULTS[10,1] <- CIRCULAR_RESULTS[8,1]/24
CIRCULAR_RESULTS[10,2] <- CIRCULAR_RESULTS[8,2]/24
CIRCULAR_RESULTS[10,3] <- CIRCULAR_RESULTS[8,3]/24
CIRCULAR_RESULTS[10,4] <- CIRCULAR_RESULTS[8,4]/24

write.csv(CIRCULAR_RESULTS, "Circular_statistics_after_phase_correction_results_BC.csv")
density_P1 <- density.circular(P1_C, bw = 10)
density_P2 <- density.circular(P2_C, bw = 10)
density_F1 <- density.circular(F1_C, bw = 10)
density_BC <- density.circular(BC_C, bw = 10)
density <- matrix(nrow = length(density_P1$x), ncol = 5)
colnames(density) <- c("angle", "P1", "P2", "F1", "BC")
 density[,1] <- density_P1$x
density[,2] <- density_P1$y
density[,3] <- density_P2$y
density[,4] <- density_F1$y
density[,5] <- density_BC$y


average_density_per_day <- matrix(nrow = 15, ncol = 5)
colnames(average_density_per_day) <- c("DAY", "P1", "P2", "F1", "BC")
average_density_per_day[,1] <- c(1:15)
average_density_per_day[1,2] <- mean(density[1:35,2])
average_density_per_day[2,2] <- mean(density[36:69,2])
average_density_per_day[3,2] <- mean(density[70:103,2])
average_density_per_day[4,2] <- mean(density[104:137,2])
average_density_per_day[5,2] <- mean(density[138:171,2])
average_density_per_day[6,2] <- mean(density[172:211,2])
average_density_per_day[7,2] <- mean(density[212:245,2])
average_density_per_day[8,2] <- mean(density[246:279,2])
average_density_per_day[9,2] <- mean(density[280:313,2])
average_density_per_day[10,2] <- mean(density[314:347,2])
average_density_per_day[11,2] <- mean(density[348:381,2])
average_density_per_day[12,2] <- mean(density[382:415,2])
average_density_per_day[13,2] <- mean(density[416:449,2])
average_density_per_day[14,2] <- mean(density[450:483,2])
average_density_per_day[15,2] <- mean(density[484:512,2])

average_density_per_day[1,3] <- mean(density[1:35,3])
average_density_per_day[2,3] <- mean(density[36:69,3])
average_density_per_day[3,3] <- mean(density[70:103,3])
average_density_per_day[4,3] <- mean(density[104:137,3])
average_density_per_day[5,3] <- mean(density[138:171,3])
average_density_per_day[6,3] <- mean(density[172:211,3])
average_density_per_day[7,3] <- mean(density[212:245,3])
average_density_per_day[8,3] <- mean(density[246:279,3])
average_density_per_day[9,3] <- mean(density[280:313,3])
average_density_per_day[10,3] <- mean(density[314:347,3])
average_density_per_day[11,3] <- mean(density[348:381,3])
average_density_per_day[12,3] <- mean(density[382:415,3])
average_density_per_day[13,3] <- mean(density[416:449,3])
average_density_per_day[14,3] <- mean(density[450:483,3])
average_density_per_day[15,3] <- mean(density[484:512,3])

average_density_per_day[1,4] <- mean(density[1:35,4])
average_density_per_day[2,4] <- mean(density[36:69,4])
average_density_per_day[3,4] <- mean(density[70:103,4])
average_density_per_day[4,4] <- mean(density[104:137,4])
average_density_per_day[5,4] <- mean(density[138:171,4])
average_density_per_day[6,4] <- mean(density[172:211,4])
average_density_per_day[7,4] <- mean(density[212:245,4])
average_density_per_day[8,4] <- mean(density[246:279,4])
average_density_per_day[9,4] <- mean(density[280:313,4])
average_density_per_day[10,4] <- mean(density[314:347,4])
average_density_per_day[11,4] <- mean(density[348:381,4])
average_density_per_day[12,4] <- mean(density[382:415,4])
average_density_per_day[13,4] <- mean(density[416:449,4])
average_density_per_day[14,4] <- mean(density[450:483,4])
average_density_per_day[15,4] <- mean(density[484:512,4])
average_density_per_day[1,5] <- mean(density[1:35,5])
average_density_per_day[2,5] <- mean(density[36:69,5])
average_density_per_day[3,5] <- mean(density[70:103,5])
average_density_per_day[4,5] <- mean(density[104:137,5])
average_density_per_day[5,5] <- mean(density[138:171,5])
average_density_per_day[6,5] <- mean(density[172:211,5])
average_density_per_day[7,5] <- mean(density[212:245,5])
average_density_per_day[8,5] <- mean(density[246:279,5])
average_density_per_day[9,5] <- mean(density[280:313,5])
average_density_per_day[10,5] <- mean(density[314:347,5])
average_density_per_day[11,5] <- mean(density[348:381,5])
average_density_per_day[12,5] <- mean(density[382:415,5])
average_density_per_day[13,5] <- mean(density[416:449,5])
average_density_per_day[14,5] <- mean(density[450:483,5])
average_density_per_day[15,5] <- mean(density[484:512,5])

write.csv(average_density_per_day, "average_density_per_day_BC_after_phase_correction.csv", row.names = F)
write.csv(density, "density_BC_after_phase_correction.csv", row.names = F)

######################################################## Calculate probability phenotypes############################################
probability_phenotypes_BC <- matrix(nrow = 15, ncol = 3)
colnames(probability_phenotypes_BC) <- c("DAY", "prob_sensitive", "prob_insensitive")
probability_phenotypes_BC[,1] <- c(1:15)
probability_phenotypes_BC[,2] <- (average_density_per_day[,4])/( average_density_per_day[,3] + average_density_per_day[,4])
probability_phenotypes_BC[,3] <- (average_density_per_day[,3])/( average_density_per_day[,3] + average_density_per_day[,4])
probability_phenotypes_BC_reduced <- cbind(probability_phenotypes_BC, 0)
colnames(probability_phenotypes_BC_reduced) <- c("DAY", "prob_sensitive", "prob_insensitive", "prob_insensitive_bin_03_07")
probability_phenotypes_BC_reduced[,4] <- probability_phenotypes_BC_reduced[,3]
probability_phenotypes_BC_reduced[,4][probability_phenotypes_BC_reduced[,4] <= 0.3] <- 0
probability_phenotypes_BC_reduced[,4][probability_phenotypes_BC_reduced[,4] >= 0.7] <- 1
probability_phenotypes_BC_reduced[,4][probability_phenotypes_BC_reduced[,4] > 0.3 & probability_phenotypes_BC_reduced[,4] < 0.7] <- "NA"

write.csv(probability_phenotypes_BC_reduced, "probability_phenotypes_BC_reduced_after_phase_correction.csv", row.names = F)
write.csv(probability_phenotypes_BC, "probability_phenotypes_BC_after_phase_correction.csv", row.names = F)
```
