## Supplemental Script 1 for "Genetic analysis of a phenotypic loss in the mechanosensory entrainment of a circalunar clock": Script_1_PHENOTYPING_README.txt

REQUIRED PACKAGES:

library(readr)
library(tibble)
library (circular)
library(stringr)


SCRIPT1: 

The pipeline first calculates summary circular statistics for P1,P2,F1 and F2 : "Circular_statistics_results_F2.csv"

Based on the difference in median dirrections between P1 and F1/F2, turbulence days of F1 and F2 are corrected for the phase shift: "Input_F2_PHASE_CORRECTED_FULL.csv"

Reduced matrix "Input_F2_PHASE_SHIFT_COR.csv" has the same columns as Input_F2.csv file so that both can be used for SCRIPT2. 

SCRIPT2:

We first recalculate circular statistics after phase correction to make sure that emergence peaks are matching : "Circular_statistics_after_phase_correction_results_F2.csv"

We then calculate kernels density estimates for P1,P2,F1 and F2: "density_F2_after_phase_correction.csv"

And average density estimates per emergence day: "average_density_per_day_F2_after_phase_correction.csv"

Finally, probability to find sensitive (rhythmic) or insensitive (arrhythmic) individuals on a corrected turbulence day were calculated: 

"probability_phenotypes_F2_after_phase_correction.csv"

Pipeline also generates binary phenotype based on exclusion of individuals with probabilities between 0.7 and 0.3. 

"probability_phenotypes_F2_reduced_after_phase_correction.csv"
