## Supplemental Script 2 for "Genetic analysis of a phenotypic loss in the mechanosensory entrainment of a circalunar clock": genotype_visualisation_and_classify_recombination_BACKcross_QTLpipeline.py

```
import argparse, sys
from argparse import RawTextHelpFormatter
import os
import subprocess
import math
import numpy as np
import pandas as pd
import matplotlib
matplotlib.use('agg')
import matplotlib.pyplot as plt
import seaborn as sns; sns.set()
from datetime import datetime

def parse_arguments():
	parser=argparse.ArgumentParser(description='Converts input VCF file from a BACKcross to a matrix and produces a heatmap as well as detects sites of recombination. \n\nNote that this script condenses marker patterns to a consensus pattern when markers are within the user-specified distance of each other and removes positions where \n\ti. the distribution of genotypes of individuals do not meet a user-specified-minimum fraction for mendalian inheritance requirement\n\tii. more than a user-specified fraction of the sites have either a genotype that differs from the parents or are missing \n\tii. more than a user-specified expected number of individuals show a switch in genotype\n\n',formatter_class=RawTextHelpFormatter)
	parser.add_argument('--vcffile', '-f', help='Input VCF file, an output from informative_variants.py script', type=str)
	parser.add_argument('--depthfilter', '-d', nargs='?', help='Minimum sample depth filter [0].', type=int, default=0)
	parser.add_argument('--fractionindividuals', '-p', nargs='?', help='Minimum fraction of individuals for minimum depth requirement [0.9].', type=float, default=0.9)
	parser.add_argument('--mendelianfraction', '-m', nargs='?', help='Minimum fraction of individuals for mendalian inheritance requirement [0.25].', type=float, default=0.25)
	parser.add_argument('--maxdiffmissing', '-i', nargs='?', help='Maximum fraction of the sites that can have either a genotype that differs from the parents or missing information  [0.25].', type=float, default=0.25)
	parser.add_argument('--mergedistance', '-v', nargs='?', help='Distance for merging neighbouring variant positions that are from the same RAD marker [150].', type=int, default=150)
	parser.add_argument('--maxrecombinationindividuals', '-r', nargs='?', help='Maximum number of individuals one would expect to have recombination at the same position [5].', type=int, default=5)
	parser.add_argument('--minmarkersforrecombinationregion', '-l', nargs='?', help='Minimum number of consecutive markers of the same genotype (between recombinations or between a recombination and the chr end) to accept as true recombination [4].', type=int, default=4)
	parser.add_argument('--excludesamples', '-e', help='Sample names of individuals to be excluded separated by ","', type=str)
	args=parser.parse_args()
	return(args)

def mode(vector):
    mostfreq=max(sorted(set(vector)),key=vector.count)
    return(mostfreq)

def flipcheck(hetparentgt,homparentgt,grandp1gt,grandp2gt):
	if grandp1gt!='0/1':
		if grandp1gt==homparentgt:
			switch="leave"
		else:
			switch="flip"
	elif grandp2gt!='0/1':
		if grandp2gt==homparentgt:
			switch="flip"
		else:
			switch="leave"
	else:
		switch="unsure"
	return(switch)

def window_consensus(windowmatrix):
    consensusvector=[]
    for i in range(len(windowmatrix[0])):
        windowsamplevector=[row[i] for row in windowmatrix]
        mostfreq=mode(windowsamplevector)
        #sorted is added to ensure that a consistent value is returned when there is more than one most frequent value
        consensusvector.append(mostfreq)
    return(consensusvector)

def row_check(prevrow,row):
	checksum=sum(abs(float(row[i])+float(prevrow[i])) for i in range(len(row)))
	switchrow=[float(x)*-1 for x in row]
	checksumrev=sum(abs(float(switchrow[i])+float(prevrow[i])) for i in range(len(switchrow)))
	switchrow=[str(x) for x in switchrow]
	return(checksum,checksumrev,switchrow)

def update_row(fullrow,switchrow):
	newrow=fullrow[0:3]
	newrow.extend(switchrow)
	new_genotype=int(fullrow[-1])*-1
	newrow.append(str(new_genotype))
	return(newrow)

def heatmap_check(outlistlist):
	#Run through heatmap once more to check if rows should be switched
	#start from the middle of the heatmap upwards:
	#now only switch rows when both grandparents are heterozygous, i.e. they have the same genotype
	rowsinheatmap=len(outlistlist)
	middle=int(round(rowsinheatmap/2))
	prevrow=outlistlist[middle][3:-1]
	for j in reversed(range(middle)):
		row=outlistlist[j][3:-1]
		#check if grandparents have the same genotype:
		if outlistlist[j][3]==outlistlist[j][4]:
			checksum,checksumrev,switchrow=row_check(prevrow,row)
			if checksum<checksumrev:
				newrow=update_row(outlistlist[j],switchrow)
				outlistlist[j]=newrow
				prevrow=switchrow
			else:
				prevrow=row
		else:
			prevrow=row
	#start from middle to the end of the heatmap
	prevrow=outlistlist[middle][3:-1]
	for j in range(middle+1,rowsinheatmap):
		row=outlistlist[j][3:-1]
		#check if grandparents have the same genotype:
		if outlistlist[j][3]==outlistlist[j][4]:
			checksum,checksumrev,switchrow=row_check(prevrow,row)
			if checksum<checksumrev:
				newrow=update_row(outlistlist[j],switchrow)
				outlistlist[j]=newrow
				prevrow=switchrow
			else:
				prevrow=row
		else:
			prevrow=row
	return(outlistlist)

def number_of_matches(vector1,vector2):
	#requires that both vectors are made up of values of only either 0.5 or -0.5
	nummatch=sum(vector1[i]==vector2[i] for i in range(len(vector1)))
	return(int(nummatch))

def too_many_miscalls_check(inputmatrix,maxrecombinationindividuals):
	#Run through heatmap once more to check if rows should be removed
	if len(inputmatrix)==1:
		reducedmatrix=inputmatrix
	else:
		reducedmatrix=[]
		if len(inputmatrix)>20:
			first10=[]
			for i in range(10):
				first10.append(inputmatrix[i][3:-1])
			previouspositionpatternvector=window_consensus(first10)
		else:
			firstfew=[]
			few=int(round(0.3*len(inputmatrix)))
			for i in range(few):
				firstfew.append(inputmatrix[i][3:-1])
			previouspositionpatternvector=window_consensus(firstfew)
			#previouspositionpatternvector=list(map(float,previouspositionpatternvector))
		for positioninmatrix in range(len(inputmatrix)):
			currentpositionpatternvector=inputmatrix[positioninmatrix][3:-1]
			match=number_of_matches(currentpositionpatternvector,previouspositionpatternvector)
			if ((len(currentpositionpatternvector)-match)<=maxrecombinationindividuals):
				reducedmatrix.append(inputmatrix[positioninmatrix])
				previouspositionpatternvector=currentpositionpatternvector
	return(reducedmatrix)

def collapse_markers(inputlist,distanceformerging):
	#runs through matrix to check if markers are within a specified distance and produces an reduced matrix with the consensus of neighbouring markers
	#check if marker position is within user specified distance of each other
	#if yes, concatenate the vector to the current consensus vector
	#if not, produce a consensus vector and append it to the output list
	#also produce the consensus genotype call
	collapsedoutlist=[]
	firstline=inputlist[0]
	colname=firstline[0]
	prevposition=int(colname.split(":")[1])
	currentconsensuslist=[firstline[1:]]
	for line in inputlist[1:]:
		position=int(line[0].split(":")[1])
		if position-prevposition<=distanceformerging:
			colname+="-"+str(position)
			currentconsensuslist.append(line[1:])
			prevposition=position
		else:
			outlist=[colname]
			radconsensus=window_consensus(currentconsensuslist)
			outlist.extend(radconsensus)
			collapsedoutlist.append(outlist)
			colname=line[0]
			prevposition=position
			currentconsensuslist=[line[1:]]
	outlist=[colname]
	radconsensus=window_consensus(currentconsensuslist)
	outlist.extend(radconsensus)
	collapsedoutlist.append(outlist)
	return(collapsedoutlist)

def determine_recombination(inputmatrix,header,minmarkersforrecombinationregion):
	#Determines recombination positions per individual. If an individual has more than 2 recombination positions, it is suggested to be excluded from the analysis
	#Grandparents are excluded
	exclude_samples=[]
	samplesrecombination=[]
	noindividuals=len(inputmatrix[0])-6
	chrposition=[]
	QTLtable={}
	for row in range(len(inputmatrix)):
		chrposition.append(inputmatrix[row][0])
	for ind in range(noindividuals):
		individual=ind+5
		individualvector=[]
		subsetindividualvector=[]
		originalindex=[]
		missingindex=[]
		unexpectedindex=[]
		for row in range(len(inputmatrix)):
			individualvector.append(inputmatrix[row][individual])
			if abs(float(inputmatrix[row][individual]))==0.5:
				subsetindividualvector.append(inputmatrix[row][individual])
				originalindex.append(row)
			elif float(inputmatrix[row][individual])==float(0):
				#missing
				missingindex.append(row)
			elif abs(float(inputmatrix[row][individual]))==0.25:
				#unexpected
				unexpectedindex.append(row)
		recombinationindex=[originalindex[i+1] for i in range(len(subsetindividualvector)-1) if float(subsetindividualvector[i])+float(subsetindividualvector[i+1])==0]
		#print(recombinationindex)
		#remove recombinations that within min markers for recombination distance to the ends:
		while len(recombinationindex)>0 and recombinationindex[0]<minmarkersforrecombinationregion:
			recombinationindex=recombinationindex[1:]
		while len(recombinationindex)>0 and len(inputmatrix)-recombinationindex[-1]<minmarkersforrecombinationregion:
			recombinationindex=recombinationindex[:-1]
		#minimum number of markers
		if len(recombinationindex)>1:
			recombinationdistances=[]
			minimumdistance=0
			while(minimumdistance<minmarkersforrecombinationregion and len(recombinationindex)>1):
				for pos in range(1,len(recombinationindex)):
					if pos>=len(recombinationindex):
						break
					recomdistance=recombinationindex[pos]-recombinationindex[pos-1]
					recombinationdistances.append(recomdistance)
				minimumdistance=min(recombinationdistances)
				if minimumdistance<minmarkersforrecombinationregion:
					minindex=recombinationdistances.index(minimumdistance)
					recombinationindex=recombinationindex[:minindex]+recombinationindex[minindex+2:]
					recombinationdistances=[]
				if len(recombinationindex)<=1:
						break
		if len(recombinationindex)>2:
			exclude_samples.append(header[individual-1])
		recombinationpositions=[chrposition[index] for index in recombinationindex]
		recom=header[individual-1]+"\t"+",".join(recombinationpositions)
		samplesrecombination.append(recom)
		#QTLtable info
		start=0
		value=[]
		recombinationindex.append(len(inputmatrix))
		for j in range(len(recombinationindex)):
			if mode(individualvector[start:(recombinationindex[j]+1)])=='-0.5':
				value.extend(['AA']*(recombinationindex[j]-start))
			elif mode(individualvector[start:(recombinationindex[j]+1)])=='0.5':
				value.extend(['AB']*(recombinationindex[j]-start))
			else:
				for individualpos in range(start,(recombinationindex[j])):
					if individualvector[individualpos]=='-0.5':
						value.append('AA')
					if individualvector[individualpos]=='0.5':
						value.append('AB')
					if individualvector[individualpos]=='0' or individualvector[individualpos]=='-0.0':
						value.append('NA')
					if individualvector[individualpos]=='0.25' or individualvector[individualpos]=='-0.25':
						value.append('-')
			start=recombinationindex[j]
		for j in missingindex:
			value[j]='NA'
		for j in unexpectedindex:
			value[j]='-'
		QTLtable[header[individual-1]]=value
	QTLtable['ID']=chrposition
	return(samplesrecombination,exclude_samples,QTLtable)

def int_or_str_sort(vector):
	try:
		list(map(str,sorted(map(int,vector))))
		sortedvector=list(map(str,sorted(map(int,vector))))
	except ValueError:
		sortedvector=list(sorted(vector))
	return(sortedvector)

if __name__ == '__main__':
	##input arguments and output files
	args=parse_arguments()
	print("Maximum number of individuals one would expect to have recombination at the same position is %s"%(args.maxrecombinationindividuals))
	if not args.vcffile is None:
		splitinfile=args.vcffile.split('.')
	else:
		print("Input VCF file is required")
		sys.exit()
	if not args.excludesamples is None:
		exclude=args.excludesamples.split(",")
	else:
		exclude=[]
	outputfilebase=".".join(splitinfile[:-1])+"_minDP%s_%sfraction_mend%s_diffmissing%s_distmerge%s_maxrecomb%s_minmarkers%s_exclude%sindividuals%s"%(args.depthfilter,args.fractionindividuals,args.mendelianfraction,args.maxdiffmissing,args.mergedistance,args.maxrecombinationindividuals,args.minmarkersforrecombinationregion,len(set(exclude)),"-".join(sorted(set(exclude))))
	inf=open(args.vcffile,"r+")
	##general columns are from 0 to 8
	##0 #CHROM, 1 POS, 2 ID, 3 REF, 4 ALT, 5 QUAL, 6 FILTER, 7 INFO, 8 FORMAT
	currentchr=""
	chrlist=[]
	outlistlist=[]
	finaloutlistlist={}
	allQTLtables={}
	excludeindices=[]
	firstline=True
	firstline_of_chr=True
	nextexclude=[]
	recombinationlist={}
	for line in inf:
		if line.startswith("#CHROM"):
			header=line.rstrip('\n').split('\t')
			outline=[]
			for i,name in enumerate(header[9:]):
				if len(exclude)>0:
					if name in exclude:
						excludeindices.append("%s"%(i))
					else:
						outline.append(name)
				else:
					outline.append(name)
			if len(excludeindices)>0:
				print("Excluding samples %s"%(set(exclude)))
				#print("Excluding indices %s"%(','.join(excludeindices)))
			headerlist=outline
			outheader="\t"+"\t".join(outline)+"\tGenotype_1\n"
		elif not line.startswith("##"):
			linesplit=line.rstrip('\n').split('\t')
			if firstline:
				colformat=linesplit[8].split(":")
				GT_index=int([index for index,value in enumerate(colformat) if value=='GT'][0])
				DP_index=int([index for index,value in enumerate(colformat) if value=='DP'][0])
				AD_index=int([index for index,value in enumerate(colformat) if value=='AD'][0])
				firstline=False
			#check if there max 2 alleles for the marker based on the ALT column (col 5) not containing ','
			if ',' not in linesplit[4]:
				##remove samples to be excluded
				for j in range(len(excludeindices)):
					del linesplit[int(excludeindices[j])-j+9]
				##check depth threshold
				linemin=[]
				for i in range(9,len(linesplit)):
					if len(linesplit[i])>1 and './.' not in linesplit[i]:
						depth=linesplit[i].split(":")[DP_index]
						if depth.isdigit():
							linemin.append(int(depth))
						else:
							depths=[int(alleledepth) for alleledepth in linesplit[i].split(":")[AD_index].split(",") if alleledepth.isdigit()]
							if len(depths)>0:
								linemin.append(sum(depths))
							else:
								linemin.append(0)
					else:
						linemin.append(0)
				fractionsamplesdepthfilter=sum(min>args.depthfilter for min in linemin)/float(len(linemin))
				print("Samples pasing: %s"%sum(min>args.depthfilter for min in linemin))
				print("Total samples: %s"%len(linemin))
				print("Fraction of samples passing the depth filter %s"%fractionsamplesdepthfilter)
				if fractionsamplesdepthfilter>args.fractionindividuals:
					colname="%s:%s"%(linesplit[0],linesplit[1])
					outlist=[colname]
					if linesplit[0] != currentchr:
						if len(currentchr)>0:
							if len(outlistlist)>0:
								#Run through heatmap once more to check if rows should be switched
								outlistlist=heatmap_check(outlistlist)
								if len(outlistlist)>0:
									#Run through heatmap iteratively to check if rows should be removed
									outlistlist=too_many_miscalls_check(outlistlist,args.maxrecombinationindividuals)
									if len(outlistlist)>0:
										#Run through heatmap and collapse neighbouring markers:
										outlistlist=collapse_markers(outlistlist,args.mergedistance)
										if len(outlistlist)>0:
											chrlist.append(currentchr)
											finaloutlistlist[currentchr]=outlistlist
											#Check recombination positions:
											samplesrecombination,exclude_samples,QTLtable=determine_recombination(outlistlist,headerlist,args.minmarkersforrecombinationregion)
											nextexclude.extend(exclude_samples)
											recombinationlist[currentchr]=samplesrecombination
											allQTLtables[currentchr]=QTLtable
										else:
											print("No variants retained for %s"%currentchr)
						currentchr=linesplit[0]
						print("Current Chrom is %s"%currentchr)
						outlistlist=[]
						firstline_of_chr=True
					##check parent genotypes and specify in matrix
					parent1genotype=linesplit[9].split(":")[GT_index].split("/")
					parent2genotype=linesplit[10].split(":")[GT_index].split("/")
					if len(set(parent1genotype))>len(set(parent2genotype)):
						heteroindex=9
						homoindex=10
						outlist.append('1')
						outlist.append('-1')
					elif len(set(parent2genotype))>len(set(parent1genotype)):
						heteroindex=10
						homoindex=9
						outlist.append('-1')
						outlist.append('1')
					grandparent1genotype=linesplit[11].split(":")[GT_index].split("/")
					grandparent2genotype=linesplit[12].split(":")[GT_index].split("/")
					##count genotype occurrences
					het_count=0
					hom_count=0
					unexpected_count=0
					missing_count=0
					offspringvector=[]
					skip=False
					pos_genotype=""
					for i in range(11,len(linesplit)):
						if len(linesplit[i])==0 or linesplit[i].split(":")[GT_index]=="./.":
							offspringvector.append(0)
							missing_count+=1
						elif linesplit[i].split(":")[GT_index]==linesplit[heteroindex].split(":")[GT_index]:
							het_count+=1
							offspringvector.append(0.5)
						elif linesplit[i].split(":")[GT_index]==linesplit[homoindex].split(":")[GT_index]:
							hom_count+=1
							offspringvector.append(-0.5)
						else:
							##unexpected genotype
							unexpected_count+=1
							#skip=True
							offspringvector.append(-0.25)
					if (float(unexpected_count+missing_count)/(het_count+hom_count+unexpected_count+missing_count))>args.maxdiffmissing:
						#more than 25% of the sites have either a genotype that differ from the parents or missing information
						print("Too many genotypes differ from parents or missing information. Unexpected: %s, Missing:%s"%(unexpected_count,missing_count))
						skip=True
					if (het_count+hom_count)>0:
						if float(min(het_count,hom_count))/(het_count+hom_count) < args.mendelianfraction:
							##does not meet mendalian fraction expectation
							print("Does not meet mendalian fraction expectation. Heterozygous: %s, Homozygous:%s"%(het_count,hom_count))
							skip=True
					if not skip:
						if firstline_of_chr:
							if het_count<hom_count:
								#to ensure that Genotype_1 is mostly homozygous (and so interpreting colours is more intuitive)
								prev_row=[[-0.5]*(len(outline)-2) for times in range(50)]
							else:
								prev_row=[[0.5]*(len(outline)-2) for times in range(50)]
							firstline_of_chr=False
						#flipping genotypes
						hetparentgt=linesplit[heteroindex].split(":")[GT_index]
						homparentgt=linesplit[homoindex].split(":")[GT_index]
						grandp1gt=linesplit[11].split(":")[GT_index]
						grandp2gt=linesplit[12].split(":")[GT_index]
						switch=flipcheck(hetparentgt,homparentgt,grandp1gt,grandp2gt)
						if switch=="leave":
							pos_genotype='-1'
						elif switch=="flip":
							offspringvector=[x*-1 for x in offspringvector]
							pos_genotype='1'
						elif switch=="unsure":
							check1list=[]
							check2list=[]
							##check for reversal of colours -> marker in opposition
							switchoffspringvector=[x*-1 for x in offspringvector]
							for row in prev_row:
								check1list.append(sum(abs(row[i]+offspringvector[i]) for i in range(len(offspringvector))))
								check2list.append(sum(abs(row[i]+switchoffspringvector[i]) for i in range(len(switchoffspringvector))))
							check1=sum(check1list)/float(len(check1list))
							check2=sum(check2list)/float(len(check2list))
							if check1<check2:
								##reversing the colours would show more consistency with the previous rows
								offspringvector=[x*-1 for x in offspringvector]
								pos_genotype='1'
							else:
								##not reversing the colours would show more consistency with the previous rows
								pos_genotype='-1'
						prev_row=prev_row[1:]+[offspringvector]
						for offspring in offspringvector:
							outlist.append(str(offspring))
						outlist.append(pos_genotype)
						outlistlist.append(outlist)
				else:
					print("Depth filter is not fulfilled")
					print("%s is not > %s"%(fractionsamplesdepthfilter,args.fractionindividuals))
			else:
				print("More than 2 alleles")
	print("Current number of variants: %s"%len(outlistlist))
	if len(outlistlist)>0:
		#Run through heatmap once more to check if rows should be switched
		outlistlist=heatmap_check(outlistlist)
		if len(outlistlist)>0:
			#Run through heatmap iteratively to check if rows should be removed
			outlistlist=too_many_miscalls_check(outlistlist,args.maxrecombinationindividuals)
			if len(outlistlist)>0:
				#Run through heatmap and collapse neighbouring markers:
				outlistlist=collapse_markers(outlistlist,args.mergedistance)
				if len(outlistlist)>0:
					chrlist.append(currentchr)
					finaloutlistlist[currentchr]=outlistlist
					#Check recombination positions:
					samplesrecombination,exclude_samples,QTLtable=determine_recombination(outlistlist,headerlist,args.minmarkersforrecombinationregion)
					nextexclude.extend(exclude_samples)
					recombinationlist[currentchr]=samplesrecombination
					allQTLtables[currentchr]=QTLtable
				else:
					print("No variants retained for %s"%currentchr)
	if len(finaloutlistlist)>0:
		separator=','
		pdfcmd=["pdfunite"]
		QTLout={}
		QTLout['ID']="Pheno%sSex%sID"%(separator,separator)
		QTLout['chr']="%s%s"%(separator,separator)
		sampleids=list(QTLtable.keys())
		sampleids.remove('ID')
		sampleids=int_or_str_sort(sampleids)
		for sample in sampleids:
			QTLout[sample]="%s%s%s"%(separator,separator,sample)
		for chr in chrlist:
			#extract chr number for chr string
			chrno=''.join([str(s) for s in list(chr) if s.isdigit()])
			outlistlist=finaloutlistlist[chr]
			tablefile=outputfilebase+"_"+chr+"_table.txt"
			if len(outlistlist)>0:
				out=open(tablefile,"w+")
				out.write(outheader)
				for outelement in outlistlist:
					outstr="\t".join(outelement)
					out.write(outstr+"\n")
				out.close()
			recombinationfile=outputfilebase+"_"+chr+"_recombination_detected_samplelist.txt"
			outlist=open(recombinationfile,"w+")
			outlist.write("Sample\tRecombination_Chr_Position(s)\n")
			for line in recombinationlist[chr]:
				outlist.write(line+"\n")
			outlist.close()
			pdffile=outputfilebase+"_"+chr+"_table.pdf"
			pdfcmd.append(pdffile)

			#create heatmap
			df=pd.read_csv(tablefile,sep="\t",index_col=0)
			plt.subplots(figsize=(len(df.columns)*0.25+5,len(df.index)*0.25+2))
			ax = sns.heatmap(df,cmap="RdYlBu",xticklabels=1,yticklabels=1,vmin=-1,vmax=1,linewidths=0.1,linecolor="black",square=True,cbar_kws={"shrink": 0.25})
			cbar=ax.collections[0].colorbar
			cbar.set_ticks([-1,-0.5,0,0.5,1])
			cbar.set_ticklabels(['Homozygous','Genotype_1','No Call','Genotype_2','Heterozygous'])
			plt.savefig(pdffile,bbox_inches='tight')

			QTLtable=allQTLtables[chr]
			QTLout['ID']+=separator+separator.join(QTLtable['ID'])
			QTLout['chr']+=separator+separator.join([str(chrno)]*len(QTLtable['ID']))
			for sample in sampleids:
				QTLout[sample]+=separator+separator.join(QTLtable[sample])
		outQTLfile=outputfilebase+"_QTLinput.csv"
		outQTL=open(outQTLfile,"w+")
		outQTL.write(QTLout['ID']+"\n")
		outQTL.write(QTLout['chr']+"\n")
		for sample in sampleids:
			outQTL.write(QTLout[sample]+"\n")
		outpdf=outputfilebase+".pdf"
		pdfcmd.append(outpdf)
		suffix=datetime.now().strftime('%Y-%m-%d_%H:%M:%S')
		pdffout="/".join(args.vcffile.split('/')[:-2])+"./pdfunite_command"+suffix+".txt"
		f2=open(pdffout,"w+")
		f2.write(" ".join(pdfcmd)+"\n")
		f2.close()
	else:
		print("No variants retained for all chromosomes")
	if len(set(nextexclude))>0:
		print("Multiple recombination events found in samples %s. It is recommended to exclude them from the analysis"%set(nextexclude))
```
