## Supplemental Script 2 for "Genetic analysis of a phenotypic loss in the mechanosensory entrainment of a circalunar clock": genotype_visualisation_and_classify_recombination_F1cross_QTLpipeline.py

```
import argparse, sys
from argparse import RawTextHelpFormatter
import os
import subprocess
import math
import numpy as np
import pandas as pd
import matplotlib
matplotlib.use('agg')
import matplotlib.pyplot as plt
import seaborn as sns; sns.set()
from datetime import datetime

def parse_arguments():
	parser=argparse.ArgumentParser(description='Converts input VCF file from an F1 cross to a matrix and produces a heatmap as well as detects sites of recombination. \n\nNote that this script condenses marker patterns to a consensus pattern when markers are within the user-specified distance of each other and removes positions where \n\ti. the distribution of genotypes of individuals do not meet a user-specified-minimum fraction for mendalian inheritance requirement\n\tii. more than a user-specified fraction of the sites have either a genotype that differs from the parents or are missing \n\tii. more than a user-specified expected number of individuals show a switch in genotype\n\n',formatter_class=RawTextHelpFormatter)
	parser.add_argument('--vcffile', '-f', help='Input VCF file, an output from informative_variants.py script', type=str)
	parser.add_argument('--depthfilter', '-d', nargs='?', help='Minimum sample depth filter [0].', type=int, default=0)
	parser.add_argument('--fractionindividuals', '-p', nargs='?', help='Minimum fraction of individuals for minimum depth requirement [0.9].', type=float, default=0.9)
	parser.add_argument('--mendelianfraction', '-m', nargs='?', help='Minimum fraction of individuals for mendalian inheritance requirement [0.25].', type=float, default=0.25)
	parser.add_argument('--maxdiffmissing', '-i', nargs='?', help='Maximum fraction of the sites that can have either a genotype that differs from the parents or missing information  [0.25].', type=float, default=0.25)
	parser.add_argument('--mergedistance', '-v', nargs='?', help='Distance for merging neighbouring variant positions that are from the same RAD marker [150].', type=int, default=150)
	parser.add_argument('--maxrecombinationindividuals', '-r', nargs='?', help='Maximum number of individuals one would expect to have recombination at the same position [5].', type=int, default=5)
	parser.add_argument('--minmarkersforrecombinationregion', '-l', nargs='?', help='Minimum number of consecutive markers of the same genotype (between recombinations or between a recombination and the chr end) to accept as true recombination [4].', type=int, default=4)
	parser.add_argument('--excludesamples', '-e', help='Sample names of individuals to be excluded separated by ","', type=str)
	parser.add_argument('--numberofparents', '-n', help='Number of samples that are parents', type=int)
	args=parser.parse_args()
	return(args)

def mode(vector):
    mostfreq=max(sorted(set(vector)),key=vector.count)
    return(mostfreq)

def window_consensus(windowmatrix):
    consensusvector=[]
    for i in range(len(windowmatrix[0])):
        windowsamplevector=[row[i] for row in windowmatrix]
        mostfreq=mode(windowsamplevector)
        #sorted is added to ensure that a consistent value is returned when there is more than one most frequent value
        consensusvector.append(mostfreq)
    return(consensusvector)


def number_of_matches(vector1,vector2):
	#requires that both vectors are made up of values of only either 0.5 or -0.5
	nummatch=sum(vector1[i]==vector2[i] for i in range(len(vector1)))
	return(int(nummatch))

def too_many_miscalls_check(inputmatrix,maxrecombinationindividuals):
	#Run through heatmap once more to check if rows should be removed
	##note for the F1 cross heatmap, the last position of the matrix is not the position genotype
	if len(inputmatrix)==1:
		reducedmatrix=inputmatrix
	else:
		reducedmatrix=[]
		if len(inputmatrix)>20:
			first10=[]
			for i in range(10):
				first10.append(inputmatrix[i][3:]) #not -1 for F1 cross
			previouspositionpatternvector=window_consensus(first10)
		else:
			firstfew=[]
			few=int(round(0.3*len(inputmatrix)))
			for i in range(few):
				firstfew.append(inputmatrix[i][3:]) #not -1 for F1 cross
			previouspositionpatternvector=window_consensus(firstfew)
		for positioninmatrix in range(len(inputmatrix)):
			currentpositionpatternvector=inputmatrix[positioninmatrix][3:] #not -1 for F1 cross
			match=number_of_matches(currentpositionpatternvector,previouspositionpatternvector)
			if ((len(currentpositionpatternvector)-match)<=maxrecombinationindividuals):
				reducedmatrix.append(inputmatrix[positioninmatrix])
				previouspositionpatternvector=currentpositionpatternvector
	return(reducedmatrix)

def collapse_markers(inputlist,distanceformerging):
	#runs through matrix to check if markers are within a specified distance and produces an reduced matrix with the consensus of neighbouring markers
	#check if marker position is within user specified distance of each other
	#if yes, concatenate the vector to the current consensus vector
	#if not, produce a consensus vector and append it to the output list
	collapsedoutlist=[]
	firstline=inputlist[0]
	colname=firstline[0]
	prevposition=int(colname.split(":")[1])
	currentconsensuslist=[firstline[1:]]
	for line in inputlist[1:]:
		position=int(line[0].split(":")[1])
		if position-prevposition<=distanceformerging:
			colname+="-"+str(position)
			currentconsensuslist.append(line[1:])
			prevposition=position
		else:
			outlist=[colname]
			radconsensus=window_consensus(currentconsensuslist)
			outlist.extend(radconsensus)
			collapsedoutlist.append(outlist)
			colname=line[0]
			prevposition=position
			currentconsensuslist=[line[1:]]
	outlist=[colname]
	radconsensus=window_consensus(currentconsensuslist)
	outlist.extend(radconsensus)
	collapsedoutlist.append(outlist)
	return(collapsedoutlist)

def determine_recombination(inputmatrix,header,minmarkersforrecombinationregion,numofparents):
	#Determines recombination positions per individual. If an individual has more than 2 recombination positions, it is suggested to be excluded from the analysis
	#Grandparents are excluded
	#F1 requires input number of parents and should be excluded from the recombination determination
	exclude_samples=[]
	samplesrecombination=[]
	noindividuals=len(inputmatrix[0])-(3+numofparents) #1. chr_position, 2. GP1, 3. GP2, (+numberofparents) & no position genotype
	chrposition=[]
	QTLtable={}
	for row in range(len(inputmatrix)):
		chrposition.append(inputmatrix[row][0])
	for ind in range(noindividuals):
		individual=ind+(3+numofparents)
		individualvector=[]
		subsetindividualvector=[]
		originalindex=[]
		missingindex=[]
		unexpectedindex=[]
		for row in range(len(inputmatrix)):
			individualvector.append(inputmatrix[row][individual])
			if abs(float(inputmatrix[row][individual]))==float(1):
				subsetindividualvector.append(inputmatrix[row][individual])
				originalindex.append(row)
			elif float(inputmatrix[row][individual])==float(0):
				#heterozygous
				subsetindividualvector.append(inputmatrix[row][individual])
				originalindex.append(row)
			elif float(inputmatrix[row][individual])==0.5:
				#missing
				missingindex.append(row)
			elif float(inputmatrix[row][individual])==-0.5:
				#unexpected
				unexpectedindex.append(row)
		recombinationindex=[originalindex[i+1] for i in range(len(subsetindividualvector)-1) if float(subsetindividualvector[i])-float(subsetindividualvector[i+1])!=0]
		#remove recombinations that within min markers for recombination distance to the ends:
		while len(recombinationindex)>0 and recombinationindex[0]<minmarkersforrecombinationregion:
			recombinationindex=recombinationindex[1:]
		while len(recombinationindex)>0 and len(inputmatrix)-recombinationindex[-1]<minmarkersforrecombinationregion:
			recombinationindex=recombinationindex[:-1]
		#minimum number of markers
		if len(recombinationindex)>1:
			recombinationdistances=[]
			minimumdistance=0
			while(minimumdistance<minmarkersforrecombinationregion and len(recombinationindex)>1):
				for pos in range(1,len(recombinationindex)):
					if pos>=len(recombinationindex):
						break
					recomdistance=recombinationindex[pos]-recombinationindex[pos-1]
					recombinationdistances.append(recomdistance)
				minimumdistance=min(recombinationdistances)
				if minimumdistance<minmarkersforrecombinationregion:
					minindex=recombinationdistances.index(minimumdistance)
					recombinationindex=recombinationindex[:minindex]+recombinationindex[minindex+2:]
					recombinationdistances=[]
				if len(recombinationindex)<=1:
						break
		if len(recombinationindex)>2:
			exclude_samples.append(header[individual-1])
		recombinationpositions=[chrposition[index] for index in recombinationindex]
		recom=header[individual-1]+"\t"+",".join(recombinationpositions)
		samplesrecombination.append(recom)
		#QTLtable info
		start=0
		value=[]
		recombinationindex.append(len(inputmatrix))
		for j in range(len(recombinationindex)):
			if mode(individualvector[start:(recombinationindex[j]+1)])=='1':
				value.extend(['AA']*(recombinationindex[j]-start))
			elif mode(individualvector[start:(recombinationindex[j]+1)])=='0':
				value.extend(['AB']*(recombinationindex[j]-start))
			elif mode(individualvector[start:(recombinationindex[j]+1)])=='-1':
				value.extend(['BB']*(recombinationindex[j]-start))
			else:
				for individualpos in range(start,(recombinationindex[j])):
					if individualvector[individualpos]=='1':
						value.append('AA')
					if individualvector[individualpos]=='0':
						value.append('AB')
					if individualvector[individualpos]=='-1':
						value.append('BB')
					if individualvector[individualpos]=='0.5':
						value.append('NA')
					if individualvector[individualpos]=='-0.5':
						value.append('-')
			start=recombinationindex[j]
		for j in missingindex:
			value[j]='NA'
		for j in unexpectedindex:
			value[j]='-'
		QTLtable[header[individual-1]]=value
	QTLtable['ID']=chrposition
	return(samplesrecombination,exclude_samples,QTLtable)

def int_or_str_sort(vector):
	try:
		list(map(str,sorted(map(int,vector))))
		sortedvector=list(map(str,sorted(map(int,vector))))
	except ValueError:
		sortedvector=list(sorted(vector))
	return(sortedvector)

if __name__ == '__main__':
	##input arguments and output files
	args=parse_arguments()
	print("Maximum number of individuals one would expect to have recombination at the same position is %s"%(args.maxrecombinationindividuals))
	if not args.vcffile is None:
		splitinfile=args.vcffile.split('.')
	else:
		print("Input VCF file is required")
		sys.exit()
	if not args.excludesamples is None:
		exclude=args.excludesamples.split(",")
	else:
		exclude=[]
	outputfilebase=".".join(splitinfile[:-1])+"_minDP%s_%sfraction_mend%s_diffmissing%s_distmerge%s_maxrecomb%s_minmarkers%s_exclude%sindividuals%s"%(args.depthfilter,args.fractionindividuals,args.mendelianfraction,args.maxdiffmissing,args.mergedistance,args.maxrecombinationindividuals,args.minmarkersforrecombinationregion,len(set(exclude)),"-".join(sorted(set(exclude))))
	inf=open(args.vcffile,"r+")
	##general columns are from 0 to 8
	##0 #CHROM, 1 POS, 2 ID, 3 REF, 4 ALT, 5 QUAL, 6 FILTER, 7 INFO, 8 FORMAT
	currentchr=""
	chrlist=[]
	outlistlist=[]
	finaloutlistlist={}
	allQTLtables={}
	excludeindices=[]
	firstline=True
	nextexclude=[]
	recombinationlist={}
	for line in inf:
		if line.startswith("#CHROM"):
			header=line.rstrip('\n').split('\t')
			outline=[]
			for i,name in enumerate(header[9:]):
				if len(exclude)>0:
					if name in exclude:
						excludeindices.append("%s"%(i))
					else:
						outline.append(name)
				else:
					outline.append(name)
			if len(excludeindices)>0:
				print("Excluding samples %s"%(set(exclude)))
			headerlist=outline
			outheader="\t"+"\t".join(outline)+"\n"
		elif not line.startswith("##"):
			linesplit=line.rstrip('\n').split('\t')
			if firstline:
				colformat=linesplit[8].split(":")
				GT_index=int([index for index,value in enumerate(colformat) if value=='GT'][0])
				DP_index=int([index for index,value in enumerate(colformat) if value=='DP'][0])
				AD_index=int([index for index,value in enumerate(colformat) if value=='AD'][0])
				firstline=False
			#check if there are only 2 alleles for the marker based on the ALT column (col 5) not containing ','
			if ',' not in linesplit[4]:
				##remove samples to be excluded
				for j in range(len(excludeindices)):
					del linesplit[int(excludeindices[j])-j+9]
				##check depth threshold
				linemin=[]
				for i in range(9,len(linesplit)):
					if len(linesplit[i])>1 and './.' not in linesplit[i]:
						depth=linesplit[i].split(":")[DP_index]
						if depth.isdigit():
							linemin.append(int(depth))
						else:
							depths=[int(alleledepth) for alleledepth in linesplit[i].split(":")[AD_index].split(",") if alleledepth.isdigit()]
							if len(depths)>0:
								linemin.append(sum(depths))
							else:
								linemin.append(0)
					else:
						linemin.append(0)
				fractionsamplesdepthfilter=sum(min>args.depthfilter for min in linemin)/float(len(linemin))
				print("Samples pasing: %s"%sum(min>args.depthfilter for min in linemin))
				print("Total samples: %s"%len(linemin))
				print("Fraction of samples passing the depth filter %s"%fractionsamplesdepthfilter)
				if fractionsamplesdepthfilter>args.fractionindividuals:
					colname="%s:%s"%(linesplit[0],linesplit[1])
					outlist=[colname]
					if linesplit[0] != currentchr:
						if len(currentchr)>0:
							if len(outlistlist)>0:
								#Run through heatmap iteratively to check if rows should be removed
								outlistlist=too_many_miscalls_check(outlistlist,args.maxrecombinationindividuals)
								if len(outlistlist)>0:
									#Run through heatmap and collapse neighbouring markers:
									outlistlist=collapse_markers(outlistlist,args.mergedistance)
									if len(outlistlist)>0:
										chrlist.append(currentchr)
										finaloutlistlist[currentchr]=outlistlist
										#Check recombination positions:
										samplesrecombination,exclude_samples,QTLtable=determine_recombination(outlistlist,headerlist,args.minmarkersforrecombinationregion,args.numberofparents)
										nextexclude.extend(exclude_samples)
										recombinationlist[currentchr]=samplesrecombination
										allQTLtables[currentchr]=QTLtable
									else:
										print("No variants retained for %s"%currentchr)
						currentchr=linesplit[0]
						print("Current Chrom is %s"%currentchr)
						outlistlist=[]
					##check grandparent genotypes
					grandparent1genotype=linesplit[9].split(":")[GT_index].split("/")
					grandparent2genotype=linesplit[10].split(":")[GT_index].split("/")
					grandparent1allele=grandparent1genotype[0]
					grandparent2allele=grandparent2genotype[1]
					#specify grandparent1's genotype as 1 and grandparent2's genotype as -1
					outlist.append('1')
					outlist.append('-1')
					##check parent genotypes and specify in matrix
					hetero_genotype=str(min(int(grandparent1allele),int(grandparent2allele)))+"/"+str(max(int(grandparent1allele),int(grandparent2allele)))
					for parentindex in range(args.numberofparents):
						parentgenotype=linesplit[11+parentindex].split(":")[GT_index].split("/")
						if grandparent1allele in parentgenotype and grandparent2allele in parentgenotype:
							outlist.append('0')
						elif set(parentgenotype)==set(grandparent1genotype):
							outlist.append('1')
						elif set(parentgenotype)==set(grandparent2genotype):
							outlist.append('-1')
						else:
							outlist.append('-0.5')
					##count genotype occurrences
					het_count=0
					hom1_count=0
					hom2_count=0
					unexpected_count=0
					missing_count=0
					offspringvector=[]
					skip=False
					for i in range(11+args.numberofparents,len(linesplit)):
						if len(linesplit[i])==0 or linesplit[i].split(":")[GT_index]=="./.":
							offspringvector.append(0.5)
							missing_count+=1
						elif linesplit[i].split(":")[GT_index]==hetero_genotype:
							het_count+=1
							offspringvector.append(0)
						elif linesplit[i].split(":")[GT_index]==linesplit[9].split(":")[GT_index]:
							#grandparent1 genotype
							hom1_count+=1
							offspringvector.append(1)
						elif linesplit[i].split(":")[GT_index]==linesplit[10].split(":")[GT_index]:
							#grandparent2 genotype
							hom2_count+=1
							offspringvector.append(-1)
						else:
							##unexpected genotype
							unexpected_count+=1
							#skip=True
							offspringvector.append(-0.5)
					if (float(unexpected_count+missing_count)/(het_count+hom1_count+hom2_count+unexpected_count+missing_count))>args.maxdiffmissing:
						#more than 25% of the sites have either a genotype that differ from the parents or missing information
						print("Too many genotypes differ from grandparents, parents or missing information. Unexpected: %s, Missing:%s"%(unexpected_count,missing_count))
						skip=True
					if (het_count+hom1_count+hom2_count)>0:
						if float(min(het_count,hom1_count,hom2_count))/(het_count+hom1_count+hom2_count) < args.mendelianfraction:
							##does not meet mendalian fraction expectation
							print("Does not meet mendalian fraction expectation. Heterozygous: %s, Homozygous A:%s, Homozygous B:%s"%(het_count,hom1_count,hom2_count))
							skip=True
					if not skip:
						for offspring in offspringvector:
							outlist.append(str(offspring))
						outlistlist.append(outlist)
				else:
					print("Depth filter is not fulfilled")
					print("%s is not > %s"%(fractionsamplesdepthfilter,args.fractionindividuals))
			else:
				print("More than 2 alleles")
	print("Current number of variants: %s"%len(outlistlist))
	if len(outlistlist)>0:
		#Run through heatmap iteratively to check if rows should be removed
		outlistlist=too_many_miscalls_check(outlistlist,args.maxrecombinationindividuals)
		if len(outlistlist)>0:
			#Run through heatmap and collapse neighbouring markers:
			outlistlist=collapse_markers(outlistlist,args.mergedistance)
			if len(outlistlist)>0:
				chrlist.append(currentchr)
				finaloutlistlist[currentchr]=outlistlist
				#Check recombination positions:
				samplesrecombination,exclude_samples,QTLtable=determine_recombination(outlistlist,headerlist,args.minmarkersforrecombinationregion,args.numberofparents)
				nextexclude.extend(exclude_samples)
				recombinationlist[currentchr]=samplesrecombination
				allQTLtables[currentchr]=QTLtable
			else:
				print("No variants retained for %s"%currentchr)
	if len(finaloutlistlist)>0:
		separator=','
		pdfcmd=["pdfunite"]
		QTLout={}
		QTLout['ID']="Pheno%sSex%sID"%(separator,separator)
		QTLout['chr']="%s%s"%(separator,separator)
		sampleids=list(QTLtable.keys())
		sampleids.remove('ID')
		sampleids=int_or_str_sort(sampleids)
		for sample in sampleids:
			QTLout[sample]="%s%s%s"%(separator,separator,sample)
		for chr in chrlist:
			#extract chr number for chr string
			chrno=''.join([str(s) for s in list(chr) if s.isdigit()])
			outlistlist=finaloutlistlist[chr]
			tablefile=outputfilebase+"_"+chr+"_table.txt"
			if len(outlistlist)>0:
				out=open(tablefile,"w+")
				out.write(outheader)
				for outelement in outlistlist:
					outstr="\t".join(outelement)
					out.write(outstr+"\n")
				out.close()
			recombinationfile=outputfilebase+"_"+chr+"_recombination_detected_samplelist.txt"
			outlist=open(recombinationfile,"w+")
			outlist.write("Sample\tRecombination_Chr_Position(s)\n")
			for line in recombinationlist[chr]:
				outlist.write(line+"\n")
			outlist.close()
			pdffile=outputfilebase+"_"+chr+"_table.pdf"
			pdfcmd.append(pdffile)

			#create heatmap
			df=pd.read_csv(tablefile,sep="\t",index_col=0)
			plt.subplots(figsize=(len(df.columns)*0.25+5,len(df.index)*0.25+2))
			ax = sns.heatmap(df,cmap="RdYlBu",xticklabels=1,yticklabels=1,vmin=-1,vmax=1,linewidths=0.1,linecolor="black",square=True,cbar_kws={"shrink": 0.25})
			cbar=ax.collections[0].colorbar
			cbar.set_ticks([-1,-0.5,0,0.5,1])
			cbar.set_ticklabels(['Homozygous B','Unexpected','Heterozygous','No Call','Homozygous A'])
			plt.savefig(pdffile,bbox_inches='tight')

			QTLtable=allQTLtables[chr]
			QTLout['ID']+=separator+separator.join(QTLtable['ID'])
			QTLout['chr']+=separator+separator.join([str(chrno)]*len(QTLtable['ID']))
			for sample in sampleids:
				QTLout[sample]+=separator+separator.join(QTLtable[sample])
		outQTLfile=outputfilebase+"_QTLinput.csv"
		outQTL=open(outQTLfile,"w+")
		outQTL.write(QTLout['ID']+"\n")
		outQTL.write(QTLout['chr']+"\n")
		for sample in sampleids:
			outQTL.write(QTLout[sample]+"\n")
		outpdf=outputfilebase+".pdf"
		pdfcmd.append(outpdf)
		suffix=datetime.now().strftime('%Y-%m-%d_%H:%M:%S')
		pdffout="/".join(args.vcffile.split('/')[:-2])+"./pdfunite_command"+suffix+".txt"
		f2=open(pdffout,"w+")
		f2.write(" ".join(pdfcmd)+"\n")
		f2.close()
	else:
		print("No variants retained for all chromosomes")
	if len(set(nextexclude))>0:
		print("Multiple recombination events found in samples %s. It is recommended to exclude them from the analysis"%set(nextexclude))
```
