## Supplemental Script 2 for "Genetic analysis of a phenotypic loss in the mechanosensory entrainment of a circalunar clock": informative_variants_QTLpipeline.py

```
import argparse, sys
import re

def parse_arguments():
	parser=argparse.ArgumentParser(description='Parse VCF File for Informative Variants. Variants that are heterozygous in one parent and homozygous in another parent are kept')
	parser.add_argument('--vcffile', '-f', help='Input VCF file',type=str)
	parser.add_argument('--outputdirectory', '-d', help='Directory for the output VCF file',type=str)
	parser.add_argument('--crosstype', '-c', help='Cross type, either "Backcross" or "F1cross"',type=str)
	parser.add_argument('--heterozygousparent', '-t', help='If "Backcross", sample name of the heterozygous parent',type=str)
	parser.add_argument('--homozygousparent', '-m', help='If "Backcross", sample name of the homozygous parent',type=str)
	parser.add_argument('--parents', '-p', nargs='?', help='If "F1cross", sample names of both F1 parents, separated by ","',type=str,default="")
	parser.add_argument('--grandparents', '-g', help='Sample names of both grandparents,separated by ","',type=str)
	args=parser.parse_args()
	return(args)

if __name__ == '__main__':
	##input arguments and output files
	args=parse_arguments()
	infile=args.vcffile.split('/')[-1]
	splitinfile=infile.split('.')
	outputfilebase=".".join(splitinfile[:-1])
	inf=open(args.vcffile,"r+")
	grandparents=args.grandparents.split(",")
	if args.crosstype=="F1cross" and args.parents is not None and args.parents is not "":
		parents=args.parents.split(",")
	if args.crosstype=="Backcross":
		grandparentsindices=['','']
	else:
		grandparentsindices=[]
	parentindices=[]
	firstline=True
	hetindex=[]
	variantlinecounter=0
	vcfheader=[]
	informative_counter=0
	for line in inf:
		if line.startswith("##"):
			vcfheader.append(line)
		elif line.startswith("#CHROM"):
			header=line.rstrip('\n').split('\t')
			for i,name in enumerate(header):
				if args.crosstype=="Backcross":
					##grandparent indices
					if name==grandparents[0]:
						grandparentsindices[0]=i
					if name==grandparents[1]:
						grandparentsindices[1]=i
					##parent indices
					if name==args.heterozygousparent:
						hetparentindex=i
						parentindices.append(i)
					if name==args.homozygousparent:
						homparentindex=i
						parentindices.append(i)
				elif args.crosstype=="F1cross":
					##grandparent indices
					if name in grandparents:
						grandparentsindices.append(i)
					##parent indices
					if args.parents is not None and args.parents is not "":
						if name in parents:
							parentindices.append(i)
				else:
					print("Unrecognised cross type: %s"%args.crosstype)
					sys.exit()
			print("Grandparent indices are %s"%(','.join(map(str,grandparentsindices))))
			if args.crosstype=="Backcross":
				print("Heterozygous parent index is %d"%(hetparentindex))
				print("Homozygous parent index is %d"%(homparentindex))
			else:
				if len(parentindices)>0:
					print("Parent indices are %s"%(','.join(map(str,parentindices))))
		    ##general columns are from 0 to 8
			##0 #CHROM, 1 POS, 2 ID, 3 REF, 4 ALT, 5 QUAL, 6 FILTER, 7 INFO, 8 FORMAT
			##creating an index that rearranges the columns for Backcross: parents to be the first two samples and grandparents to be in cols 3 and 4 or F1cross: the grandparents are in the first two columns and the parents are in col3 onwwards
			order=list(range(9))
			sampleorder=list(range(9,len(header)))
			if args.crosstype=="Backcross":
				order.extend([hetparentindex])
				order.extend([homparentindex])
				sampleorder.remove(hetparentindex)
				sampleorder.remove(homparentindex)
				order.extend(grandparentsindices)
				sampleorder.remove(grandparentsindices[0])
				sampleorder.remove(grandparentsindices[1])
			else:
				order.extend(grandparentsindices)
				sampleorder.remove(grandparentsindices[0])
				sampleorder.remove(grandparentsindices[1])
				order.extend(parentindices)
				if args.parents is not None and args.parents is not "":
					for parentindex in parentindices:
						sampleorder.remove(parentindex)
			order.extend(sampleorder)
			##re-ordered header as list
			outline=[header[i] for i in order]
			if args.crosstype=="Backcross":
				outputfile=args.outputdirectory+"/"+outputfilebase+"_%s_%s_het_%s_hom.vcf"%(args.crosstype,header[hetparentindex],header[homparentindex])
			else:
				outputfile=args.outputdirectory+"/"+outputfilebase+"_%s_grandparents_%s.vcf"%(args.crosstype,"_".join(grandparents))
			out=open(outputfile,"w+")
			##write headers out to file
			for vcfh in vcfheader:
				out.write(vcfh)
			realheader='\t'.join(outline)+"\n"
			out.write(realheader)
		else:
			linesplit=line.rstrip('\n').split('\t')
			if firstline:
				colformat=linesplit[8].split(":")
				##Format GT:DP:AD:RO:QR:AO:QA:GL (not necessarily)
				GT_index=int([index for index,value in enumerate(colformat) if value=='GT'][0])
				firstline=False
			else:
				##grandparent genotype filter
				##check if both have a genotype call
				print("Grandparents: %s"%[linesplit[x] for x in grandparentsindices])
				if not any(len(s)==1 for s in [linesplit[x] for x in grandparentsindices]):
					grandparent1genotype=linesplit[grandparentsindices[0]].split(":")[GT_index].split("/")
					grandparent2genotype=linesplit[grandparentsindices[1]].split(":")[GT_index].split("/")
					##CHECK: if both homozygous the alleles are different between the grandparents, else at least one grandparent is heterozygous
					if (len(set(grandparent1genotype))==1 and len(set(grandparent2genotype))==1 and set(grandparent1genotype)!=set(grandparent2genotype)) or (max(len(set(grandparent1genotype)),len(set(grandparent2genotype)))==2):
						#CHECK: no genotypes are ./. i.e. missing
						if '.' not in grandparent1genotype and '.' not in grandparent2genotype:
							if args.crosstype=="Backcross":
								print("grandparents fulfill criteria: %s %s"%(grandparent1genotype,grandparent2genotype))
								##parent genotype filter
								##check if both have a genotype call
								if not any(len(s)==1 for s in [linesplit[x] for x in parentindices]):
									hetparentgenotype=linesplit[hetparentindex].split(":")[GT_index].split("/")
									homparentgenotype=linesplit[homparentindex].split(":")[GT_index].split("/")
									##CHECK: Hom Parent has 1 allele twice and Het Parent has 2 different alleles
									if (len(set(homparentgenotype))==1 and len(set(hetparentgenotype))==2):
										##CHECK: the homozygous parent allele is present in the heterozygous parent
										if set(homparentgenotype).issubset(set(hetparentgenotype)):
											#check if heterozygous parent has genotype 0/1
											if linesplit[hetparentindex].split(":")[GT_index]=='0/1':
												outline=[linesplit[i] for i in order]
												outstr='\t'.join(outline)+"\n"
												out.write(outstr)
												informative_counter+=1
							elif args.crosstype=="F1cross":
								##CHECK: Both grandparents are homozygous
								if (len(set(grandparent1genotype))==1 and len(set(grandparent2genotype))==1):
									print("grandparents fulfill criteria: %s %s"%(grandparent1genotype,grandparent2genotype))
									if args.parents is not None and args.parents is not "":
										##CHECK: confirm that all of the parents have the same genotype
										parent_genotypes={}
										for parentindex in parentindices:
											if len(linesplit[parentindex])>1:
												parentgenotype=linesplit[parentindex].split(":")[GT_index]
												print(parentgenotype)
												if parentgenotype in list(parent_genotypes.keys()):
													parent_genotypes[parentgenotype]+=1
												else:
													parent_genotypes[parentgenotype]=1
										print("most freq: %d"%(max(parent_genotypes.values())))
										print("frac: %d"%(float(max(parent_genotypes.values()))/sum(parent_genotypes.values())))
										if float(max(parent_genotypes.values()))/sum(parent_genotypes.values())==1:
											most_freq_genotype=max(parent_genotypes,key=parent_genotypes.get)
											most_freq_genotype_split=most_freq_genotype.split(":")[GT_index].split("/")
											##CHECK: confirm that the majority genotype is heterozygous
											if len(set(most_freq_genotype_split))==2:
												##CHECK: confirm that the grandparent's alleles are in the genotype
												print("genotype: %s"%most_freq_genotype)
												if set(grandparent1genotype).issubset(set(most_freq_genotype_split)) and set(grandparent2genotype).issubset(set(most_freq_genotype_split)):
													outline=[linesplit[i] for i in order]
													outstr='\t'.join(outline)+"\n"
													out.write(outstr)
													informative_counter+=1
									else:
										outline=[linesplit[i] for i in order]
										outstr='\t'.join(outline)+"\n"
										out.write(outstr)
										informative_counter+=1
	if args.crosstype=="Backcross":
		print("%s heterozygous sites in %s and homozygous sites in %s"%(informative_counter,header[hetparentindex],header[homparentindex]))
	else:
		print("%s homozygous sites with different alleles in %s and in %s"%(informative_counter,header[int(grandparentsindices[0])],header[int(grandparentsindices[1])]))
```
