## Supplemental Script 2 for "Genetic analysis of a phenotypic loss in the mechanosensory entrainment of a circalunar clock": QTL_input_pipeline.py

```
from __future__ import print_function
import os
import os.path
import argparse, sys
from argparse import RawTextHelpFormatter
import subprocess

#input arguments:
# - Input table of sample id, fastq file, (fastq file 2)
# - Output directory
# - Reference genome file
# - sample id for grandmother
# - sample id for grandfather
# - sample id for mother
# - sample id for father

#1. Read input table:
#input table should have sample id, fastq file, (optional fastq file 2)
#For each sample
    #1. Trim adapters
    #2. Merge overlapping reads with PEAR
    #3. Map reads
    #4. Sort Reads
    #5. Index Reads
    #6. Variant Calling with GATK
#2. Merge all samples
#3. Informative variants
#3. Genotype visualistation
#4. Identification of sites of recombination

def parse_arguments():
    parser=argparse.ArgumentParser(description='Pipeline to prepare RAD-seq data for QTL analysis:\n\n\t1. trimming of adapters with trimmomatic\n\t2. merging overlapping reads with PEAR \n\t3. mapping with NextGenMap\n\t4. variant calling with GATK\n\t5. obtaining variants that are informative\n\t6. converting the resulting VCF file to a matrix and producing a heatmap\n\t7. detecting sites of recombination\n\t8. producing an input file for QTL analysis.\n\n\n    *Please run "module load python/2.7.13" and "module load java/x64/8u121" before using this script*\n\n\n', formatter_class=RawTextHelpFormatter)
    parser.add_argument('--inputtable', '-i', help='Input table without a header. Columns of table should be: 1. sample id  2. input fastq file (with path), [if paired-end: 3. input fastq file for read 2]',type=str)
    parser.add_argument('--outputdirectory', '-o', help='Output directory where output files should be written',type=str)
    parser.add_argument('--adapterfile', '-g', nargs='?', help='Adapter sequences for Trimmomatic',type=str, default="")
    parser.add_argument('--mintrimmedlength', '-t', nargs='?', help='Trimmomatic minimum trimmed read length [50]',type=int, default=50)
    parser.add_argument('--reference', '-r', help='Reference genome file (with path)',type=str)
    parser.add_argument('--minreadidentity', '-j', nargs='?', help='NextGenMap minimum read mapping identity [0.95]',type=float, default=0.95)
    parser.add_argument('--minreadfraction', '-s', nargs='?', help='NextGenMap minimum read mapping fraction [1]',type=float, default=1)
    parser.add_argument('--mergedvcffile', '-v', help='Filename for merged VCF file',type=str)
    parser.add_argument('--maxprocesses', '-m', nargs='?', help='Maximum number of processes to run in parallel [1]',type=int,default=1)
    parser.add_argument('--crosstype', '-c', nargs='?', help='Cross type, either "Backcross" or "F1cross" [Backcross]',type=str,default="Backcross")
    parser.add_argument('--heterozygousparent', '-a', help='If "Backcross", sample name of the heterozygous parent',type=str)
    parser.add_argument('--homozygousparent', '-b', help='If "Backcross", sample name of the homozygous parent',type=str)
    parser.add_argument('--grandparentstrain1', '-d', help='If "Backcross", sample name of grandparent that is of the same strain as the homozygous parent',type=str)
    parser.add_argument('--grandparentstrain2', '-e', help='If "Backcross", sample name of grandparent that is not of the same strain as the homozygous parent',type=str)
    parser.add_argument('--parents', '-z', nargs='?', help='If "F1cross", sample names of both F1 parents, separated by ","',type=str,default="")
    parser.add_argument('--grandparents', '-p', help='Sample names of both grandparents, separated by ","',type=str)
    args=parser.parse_args()
    return(args)

def run_cmd_check(cmd,outfile):
    if not os.path.isfile(outfile):
        print(cmd)
        print("")
        runcmd=True
    else:
        print("%s exists"%(outfile))
        print("")
        runcmd=False
    return(runcmd)

def input_absent(inputfiles):
    absent=[]
    for file in inputfiles:
        if not os.path.isfile(file):
            absent.append(file)
    return(absent)

def common_prefix(a,b):
    return(a[:max(j for j in range(len(b)) if a.startswith(b[:j]))])

def trim_adapters(fastqfile,adapterfile,mintrimmedreadlength,outputdirectory):
    base=fastqfile.split("/")[-1].split(".")[0]
    trimmedfile="%s/%s_trimmed.fastq"%(outputdirectory,base)
    cmd="java -jar /data/biosoftware/Trimmomatic/Trimmomatic-0.38/trimmomatic-0.38.jar SE -threads 1 -summary %s/%s.statsSummaryFile %s %s ILLUMINACLIP:%s:2:30:10 LEADING:20 TRAILING:20 MINLEN:%s"%(outputdirectory,base,fastqfile,trimmedfile,adapterfile,mintrimmedreadlength)
    return(cmd,trimmedfile)

def trim_adapters_paired(fastqfile1,fastqfile2,adapterfile,mintrimmedreadlength,outputdirectory):
    base1=fastqfile1.split("/")[-1].split(".")[0]
    pairedtrimmedfile1="%s/%s_paired_trimmed.fastq"%(outputdirectory,base1)
    unpairedtrimmedfile1="%s/%s_unpaired_trimmed.fastq"%(outputdirectory,base1)
    base2=fastqfile2.split("/")[-1].split(".")[0]
    pairedtrimmedfile2="%s/%s_paired_trimmed.fastq"%(outputdirectory,base2)
    unpairedtrimmedfile2="%s/%s_unpaired_trimmed.fastq"%(outputdirectory,base2)
    base=common_prefix(base1,base2)
    cmd="java -jar /data/biosoftware/Trimmomatic/Trimmomatic-0.38/trimmomatic-0.38.jar PE -threads 1 -summary %s/%s.statsSummaryFile %s %s %s %s %s %s ILLUMINACLIP:%s:2:30:10:2:true LEADING:20 TRAILING:20 MINLEN:%s"%(outputdirectory,base,fastqfile1,fastqfile2,pairedtrimmedfile1,unpairedtrimmedfile1,pairedtrimmedfile2,unpairedtrimmedfile2,adapterfile,mintrimmedreadlength)
    return(cmd,pairedtrimmedfile1,unpairedtrimmedfile1,pairedtrimmedfile2,unpairedtrimmedfile2)

def assemble_overlap_pairs(fastqfile1,fastqfile2,outputdirectory):
    commonbase=common_prefix(fastqfile1,fastqfile2)
    base=commonbase.split("/")[-1]+"_PEAR"
    assembledfile="%s/%s.assembled.fastq"%(outputdirectory,base)
    unassembledforwardfile="%s/%s.unassembled.forward.fastq"%(outputdirectory,base)
    unassembledreversefile="%s/%s.unassembled.reverse.fastq"%(outputdirectory,base)
    discardfile="%s/%s.discarded.fastq"%(outputdirectory,base)
    cmd="/data/biosoftware/pear/pear-0.9.10-bin-64/pear-0.9.10-bin-64 -f %s -r %s -o %s/%s -n 50 -c 40 -k"%(fastqfile1,fastqfile2,outputdirectory,base)
    return(cmd,assembledfile,unassembledforwardfile,unassembledreversefile,discardfile)

def ngm_interleave(fastqfile1,fastqfile2):
    commonbase=common_prefix(fastqfile1,fastqfile2)
    interleavedfile=commonbase+"_interleaved.fastq"
    cmd="ngm-utils interleave -1 %s -2 %s -o %s"%(fastqfile1,fastqfile2,interleavedfile)
    return(cmd,interleavedfile)

def map_reads(trimmedfile,reference,sampleid,outputdirectory,minidentity,minreadlengthfraction,paired):
    base=".".join(trimmedfile.split("/")[-1].split(".")[0:-1])
    mappedfile="%s/%s_ngm_i%s_R%s.bam"%(outputdirectory,base,minidentity,minreadlengthfraction)
    if paired:
        cmd="ngm -q %s -r %s -b -o %s --rg-id %s --rg-sm %s --rg-lb lb --rg-pl illumina --rg-pu pu -i %s -R %s -p"%(trimmedfile,reference,mappedfile,sampleid,sampleid,minidentity,minreadlengthfraction)
    else:
        cmd="ngm -q %s -r %s -b -o %s --rg-id %s --rg-sm %s --rg-lb lb --rg-pl illumina --rg-pu pu -i %s -R %s"%(trimmedfile,reference,mappedfile,sampleid,sampleid,minidentity,minreadlengthfraction)
    return(cmd,mappedfile)

def sort_bam(mappedfile):
    base=".".join(mappedfile.split(".")[0:-1])
    sortedfile="%s_sorted.bam"%(base)
    cmd="samtools sort -o %s %s"%(sortedfile,mappedfile)
    return(cmd,sortedfile)

def index_bam(sortedfile):
    cmd="samtools index %s"%(sortedfile)
    return(cmd)

def haplotype_caller(reference,sortedfile,outputdirectory):
    base=".".join(sortedfile.split("/")[-1].split(".")[0:-1])
    vcffile="%s/%s_q20.g.vcf"%(outputdirectory,base)
    cmd="java -jar /data/biosoftware/GATK/GATK/GenomeAnalysisTK.jar -T HaplotypeCaller --emitRefConfidence GVCF -R %s -I %s -stand_call_conf 30 -o %s"%(reference,sortedfile,vcffile)
    return(cmd,vcffile)

def select_variants(reference,vcffile):
    base=".".join(vcffile.split(".")[0:-2])
    filteredvcf="%s_filtered.g.vcf"%(base)
    cmd="java -jar /data/biosoftware/GATK/GATK/GenomeAnalysisTK.jar -T SelectVariants -R %s --variant %s -o %s -select"%(reference,vcffile,filteredvcf)
    return(cmd,filteredvcf)

def base_recalibrator(reference,filteredvcf,sortedfile,outputdirectory):
    base=".".join(sortedfile.split("/")[-1].split(".")[0:-1])
    reportfile="%s/%s_recalibration_report.grp"%(outputdirectory,base)
    cmd="java -jar /data/biosoftware/GATK/GATK/GenomeAnalysisTK.jar -T BaseRecalibrator -R %s -knownSites %s -I %s -o %s"%(reference,filteredvcf,sortedfile,reportfile)
    return(cmd,reportfile)

def print_reads(reference,sortedfile,reportfile,outputdirectory):
    base=".".join(sortedfile.split("/")[-1].split(".")[0:-1])
    recallbam="%s/%s.recall.bam"%(outputdirectory,base)
    cmd="java -jar /data/biosoftware/GATK/GATK/GenomeAnalysisTK.jar -T PrintReads -R %s -I %s -BQSR %s -o %s"%(reference,sortedfile,reportfile,recallbam)
    return(cmd,recallbam)

def delete_files(listoffiles):
    flat_list = [item for sublist in listoffiles for item in sublist]
    for file in flat_list:
        if os.path.isfile(file):
            os.system("rm %s"%file)

if __name__ == '__main__':
    ##input arguments and output files
    args=parse_arguments()

    print("")
    print("")
    print("---------------------------------------------------------------------------------------------------------------------------------")
    print("                                                 ##QTL Input Pipeline##")
    print("---------------------------------------------------------------------------------------------------------------------------------")
    print("")
    print("")
    #check if output directory exists, if not make it.
    if not os.path.isdir(args.outputdirectory):
        cmd="mkdir %s"%(args.outputdirectory)
        print("Making directory %s"%(args.outputdirectory))
        print(cmd)
        print("")
        os.system(cmd)

    #trimming reads
    print("##Trimming Reads With Trimmomatic##")
    print("---------------------------------------------------------------------------------------------------------------------------------")
    print("")
    print("")

    #check if output directory for trimmed reads exists, if not make it.
    trimmeddir="%s/trimmed_reads"%(args.outputdirectory)
    if not os.path.isdir(trimmeddir):
    	cmd="mkdir %s"%(trimmeddir)
    	print("Making directory %s"%(trimmeddir))
    	print(cmd)
    	print("")
    	os.system(cmd)

    processes=set()
    trimmedfiles={}
    sampleids=[]
    error_counter=0
    #read input table
    table=open(args.inputtable,"r+")
    for line in table:
        line=line.rstrip("\n").split()
        #Columns of table should be: sample id, input fastq file (with path), input fastq file 2 (with path)
        if len(line)>1:
            sampleid=line[0]
            sampleids.append(sampleid)
            fastqfile=line[1]
            inputfiles=[fastqfile]
            if len(line)>2:
                fastqfile2=line[2]
                inputfiles.append(fastqfile2)
                cmd,pairedtrimmedfile1,unpairedtrimmedfile1,pairedtrimmedfile2,unpairedtrimmedfile2=trim_adapters_paired(fastqfile,fastqfile2,args.adapterfile,args.mintrimmedlength,outputdirectory=trimmeddir)
                trimmedfiles[sampleid]=[pairedtrimmedfile1,unpairedtrimmedfile1,pairedtrimmedfile2,unpairedtrimmedfile2]
                paired=True
                runcmd=run_cmd_check(cmd,pairedtrimmedfile1)
            else:
                cmd,trimmedfile=trim_adapters(fastqfile,args.adapterfile,args.mintrimmedlength,outputdirectory=trimmeddir)
                trimmedfiles[sampleid]=trimmedfile
                paired=False
                runcmd=run_cmd_check(cmd,trimmedfile)
            if len(input_absent(inputfiles))>0:
               print("missing: "+",".join(input_absent(inputfiles)))
               sys.exit(1)
            if runcmd:
               processes.add(subprocess.Popen(cmd.split()))
               if len(processes)>args.maxprocesses:
                   os.wait()
                   processes.difference_update([p for p in processes if p.poll() is not None])
    #check if all the processes were closed
    for p in processes:
    	if p.poll() is None:
    		p.wait()
    for p in processes:
       stdout,stderr=p.communicate()
       if stderr is not None and stderr !=0:
           error_counter+=1
    if error_counter>0:
       delete_files(list(trimmedfiles.values()))
       sys.exit(1)

    #assemble overlapping read pairs
    if paired:
        print("##Assembling Overlapping Read-Pairs With PEAR##")
        print("---------------------------------------------------------------------------------------------------------------------------------")
        print("")
        print("")

        assembleddir="%s/assembled_reads"%(args.outputdirectory)
        if not os.path.isdir(assembleddir):
        	cmd="mkdir %s"%(assembleddir)
        	print("Making directory %s"%(assembleddir))
        	print(cmd)
        	print("")
        	os.system(cmd)

        pear_out_files={}
        error_counter=0

        for sampleid in sampleids:
            pairedtrimmedfile1=trimmedfiles[sampleid][0]
            pairedtrimmedfile2=trimmedfiles[sampleid][2]
            cmd,assembledfile,unassembledforwardfile,unassembledreversefile,discardfile=assemble_overlap_pairs(pairedtrimmedfile1,pairedtrimmedfile2,outputdirectory=assembleddir)
            pear_out_files[sampleid]=[assembledfile,unassembledforwardfile,unassembledreversefile,discardfile]
            runcmd=run_cmd_check(cmd,assembledfile)
            inputfiles=[pairedtrimmedfile1,pairedtrimmedfile2]
            if len(input_absent(inputfiles))>0:
               print("missing: "+",".join(input_absent(inputfiles)))
               sys.exit(1)
            if runcmd:
               processes.add(subprocess.Popen(cmd.split()))
               if len(processes)>args.maxprocesses:
                   os.wait()
                   processes.difference_update([p for p in processes if p.poll() is not None])

        #check if all the processes were closed
        for p in processes:
        	if p.poll() is None:
        		p.wait()
        for p in processes:
           if p.poll() is not None and p.poll() !=0:
               error_counter+=1
        if error_counter>0:
           delete_files(list(pear_out_files.values()))
           sys.exit(1)

    #interleaving unassembled paired end reads before mapping
    if paired:
        print("##Interleaving Reads With NextGenMap##")
        print("---------------------------------------------------------------------------------------------------------------------------------")
        print("")
        print("")

        interleavedfiles={}
        error_counter=0
        for sampleid in sampleids:
            unassembledtrimmedfile1=pear_out_files[sampleid][1]
            unassembledtrimmedfile2=pear_out_files[sampleid][2]
            cmd,interleavedfile=ngm_interleave(unassembledtrimmedfile1,unassembledtrimmedfile2)
            interleavedfiles[sampleid]=interleavedfile
            runcmd=run_cmd_check(cmd,interleavedfile)
            inputfiles=[unassembledtrimmedfile1,unassembledtrimmedfile2]
            if len(input_absent(inputfiles))>0:
               print("missing: "+",".join(input_absent(inputfiles)))
               sys.exit(1)
            if runcmd:
               processes.add(subprocess.Popen(cmd.split()))
               if len(processes)>args.maxprocesses:
                   os.wait()
                   processes.difference_update([p for p in processes if p.poll() is not None])
        #check if all the processes were closed
        for p in processes:
        	if p.poll() is None:
        		p.wait()
        for p in processes:
           if p.poll() is not None and p.poll() !=0:
               error_counter+=1
        if error_counter>0:
           delete_files(list(interleavedfiles.values()))
           sys.exit(1)


    #concatenating single read sets
    #single reads from the paired-end library -> assembled pairs, trimmed-unpaired forward and trimmed-unpaired reverse
    if paired:
        print("##Concatenating single read sets (PEAR assembled and Trimmomatic trimmed but unpaired)##")
        print("---------------------------------------------------------------------------------------------------------------------------------")
        print("")
        print("")

        #concatenate read sets
        singledir="%s/single_reads"%(args.outputdirectory)
        if not os.path.isdir(singledir):
        	cmd="mkdir %s"%(singledir)
        	print("Making directory %s"%(singledir))
        	print(cmd)
        	print("")
        	os.system(cmd)

        singlereadsfiles={}
        error_counter=0
        for sampleid in sampleids:
            assembledfile=pear_out_files[sampleid][0]
            unpairedtrimmedfile1=trimmedfiles[sampleid][1]
            unpairedtrimmedfile2=trimmedfiles[sampleid][3]
            singlereadsfile="%s/%s_single_reads.fastq"%(singledir,sampleid)
            cmd="cat %s %s %s > %s"%(assembledfile,unpairedtrimmedfile1,unpairedtrimmedfile2,singlereadsfile)
            singlereadsfiles[sampleid]=singlereadsfile
            runcmd=run_cmd_check(cmd,singlereadsfile)
            if runcmd:
                os.system(cmd)


	#mapping reads
    print("##Mapping Reads With NextGenMap##")
    print("---------------------------------------------------------------------------------------------------------------------------------")
    print("")
    print("")

    mappeddir="%s/mapped_reads"%(args.outputdirectory)
    if not os.path.isdir(mappeddir):
    	cmd="mkdir %s"%(mappeddir)
    	print("Making directory %s"%(mappeddir))
    	print(cmd)
    	print("")
    	os.system(cmd)

    mappedfiles={}
    error_counter=0
    for sampleid in sampleids:
        if paired:
            trimmedfile=interleavedfiles[sampleid]
        else:
            trimmedfile=trimmedfiles[sampleid]
        cmd,mappedfile=map_reads(trimmedfile,args.reference,sampleid,mappeddir,args.minreadidentity,args.minreadfraction,paired)
        mappedfiles[sampleid]=mappedfile
        runcmd=run_cmd_check(cmd,mappedfile)
        inputfiles=[trimmedfile]
        if len(input_absent(inputfiles))>0:
           print("missing: "+",".join(input_absent(inputfiles)))
           sys.exit(1)
        if runcmd:
            processes.add(subprocess.Popen(cmd.split()))
            if len(processes)>args.maxprocesses:
                os.wait()
                processes.difference_update([p for p in processes if p.poll() is not None])
    #check if all the processes were closed
    for p in processes:
    	if p.poll() is None:
    		p.wait()
    for p in processes:
        if p.poll() is not None and p.poll() !=0:
            error_counter+=1
    if error_counter>0:
        delete_files(list(mappedfiles.values()))
        sys.exit(1)

    if paired:
        #map single reads
        singlereadsmappedfiles={}
        error_counter=0
        for sampleid in sampleids:
            singlereadsfile=singlereadsfiles[sampleid]
            cmd,singlereadsmappedfile=map_reads(singlereadsfile,args.reference,sampleid,mappeddir,args.minreadidentity,args.minreadfraction,paired=False)
            singlereadsmappedfiles[sampleid]=singlereadsmappedfile
            runcmd=run_cmd_check(cmd,singlereadsmappedfile)
            inputfiles=[singlereadsfile]
            if len(input_absent(inputfiles))>0:
               print("missing: "+",".join(input_absent(inputfiles)))
               sys.exit(1)
            if runcmd:
                processes.add(subprocess.Popen(cmd.split()))
                if len(processes)>args.maxprocesses:
                    os.wait()
                    processes.difference_update([p for p in processes if p.poll() is not None])
        #check if all the processes were closed
        for p in processes:
        	if p.poll() is None:
        		p.wait()
        for p in processes:
            if p.poll() is not None and p.poll() !=0:
                error_counter+=1
        if error_counter>0:
            delete_files(list(singlereadsmappedfiles.values()))
            sys.exit(1)


    #sorting reads
    print("##Sorting Reads With Samtools##")
    print("----------------------------------------------------------------")
    print("")
    print("")

    sortedfiles={}
    error_counter=0
    for sampleid in sampleids:
        mappedfile=mappedfiles[sampleid]
        cmd,sortedfile=sort_bam(mappedfile)
        sortedfiles[sampleid]=sortedfile
        runcmd=run_cmd_check(cmd,sortedfile)
        inputfiles=[mappedfile]
        if len(input_absent(inputfiles))>0:
           print("missing: "+",".join(input_absent(inputfiles)))
           sys.exit(1)
        if runcmd:
           processes.add(subprocess.Popen(cmd.split()))
           if len(processes)>args.maxprocesses:
               os.wait()
               processes.difference_update([p for p in processes if p.poll() is not None])
    #check if all the processes were closed
    for p in processes:
    	if p.poll() is None:
    		p.wait()
    for p in processes:
       if p.poll() is not None and p.poll() !=0:
           error_counter+=1
    if error_counter>0:
       delete_files(list(sortedfiles.values()))
       sys.exit(1)

    if paired:
        sortedsinglefiles={}
        error_counter=0
        #sorting single mapped reads:
        for sampleid in sampleids:
            singlereadsmappedfile=singlereadsmappedfiles[sampleid]
            cmd,sortedfile=sort_bam(singlereadsmappedfile)
            sortedsinglefiles[sampleid]=sortedfile
            runcmd=run_cmd_check(cmd,sortedfile)
            inputfiles=[singlereadsmappedfile]
            if len(input_absent(inputfiles))>0:
               print("missing: "+",".join(input_absent(inputfiles)))
               sys.exit(1)
            if runcmd:
               processes.add(subprocess.Popen(cmd.split()))
               if len(processes)>args.maxprocesses:
                   os.wait()
                   processes.difference_update([p for p in processes if p.poll() is not None])
        #check if all the processes were closed
        for p in processes:
        	if p.poll() is None:
        		p.wait()
        for p in processes:
           if p.poll() is not None and p.poll() !=0:
               error_counter+=1
        if error_counter>0:
           delete_files(list(sortedsinglefiles.values()))
           sys.exit(1)

        #merge bam files for mapped single reads and mapped paired reads
        sortedpairedfiles=sortedfiles
        mergedfiles={}
        error_counter=0
        for sampleid in sampleids:
            pairedsortedfile=sortedpairedfiles[sampleid]
            singlesortedfile=sortedsinglefiles[sampleid]
            mergedmappedfile="%s/%s_paired_and_single_merged_ngm_i%s_R%s.bam"%(mappeddir,sampleid,args.minreadidentity,args.minreadfraction)
            cmd="samtools merge -u -c -p %s %s %s"%(mergedmappedfile,pairedsortedfile,singlesortedfile)
            mergedfiles[sampleid]=mergedmappedfile
            runcmd=run_cmd_check(cmd,mergedmappedfile)
            inputfiles=[pairedsortedfile,singlesortedfile]
            if len(input_absent(inputfiles))>0:
               print("missing: "+",".join(input_absent(inputfiles)))
               sys.exit(1)
            if runcmd:
               processes.add(subprocess.Popen(cmd.split()))
               if len(processes)>args.maxprocesses:
                   os.wait()
                   processes.difference_update([p for p in processes if p.poll() is not None])
        #check if all the processes were closed
        for p in processes:
        	if p.poll() is None:
        		p.wait()
        for p in processes:
           if p.poll() is not None and p.poll() !=0:
               error_counter+=1
        if error_counter>0:
           delete_files(list(mergedfiles.values()))
           sys.exit(1)

        #sort merged bam files
        sortedfiles=[]
        error_counter=0
        for sampleid in sampleids:
            mergedmappedfile=mergedfiles[sampleid]
            cmd,sortedfile=sort_bam(mergedmappedfile)
            sortedfiles.append(sortedfile)
            runcmd=run_cmd_check(cmd,sortedfile)
            inputfiles=[mergedmappedfile]
            if len(input_absent(inputfiles))>0:
               print("missing: "+",".join(input_absent(inputfiles)))
               sys.exit(1)
            if runcmd:
                processes.add(subprocess.Popen(cmd.split()))
                if len(processes)>args.maxprocesses:
                    os.wait()
                    processes.difference_update([p for p in processes if p.poll() is not None])
        #check if all the processes were closed
        for p in processes:
        	if p.poll() is None:
        		p.wait()
        for p in processes:
            if p.poll() is not None and p.poll() !=0:
                error_counter+=1
        if error_counter>0:
            delete_files(sortedfiles)
            sys.exit(1)
    else:
        sortedfiles_dict=sortedfiles
        sortedfiles=list(sortedfiles_dict.values())

    #indexing reads
    print("##Indexing Reads With Samtools##")
    print("----------------------------------------------------------------")
    print("")
    print("")

    error_counter=0
    for sortedfile in sortedfiles:
        cmd=index_bam(sortedfile)
        indexfile="%s.bai"%(sortedfile)
        runcmd=run_cmd_check(cmd,indexfile)
        inputfiles=[sortedfile]
        if len(input_absent(inputfiles))>0:
           print("missing: "+",".join(input_absent(inputfiles)))
           sys.exit(1)
        if runcmd:
            processes.add(subprocess.Popen(cmd.split()))
            if len(processes)>args.maxprocesses:
                os.wait()
                processes.difference_update([p for p in processes if p.poll() is not None])
    #check if all the processes were closed
    for p in processes:
    	if p.poll() is None:
    		p.wait()
    for p in processes:
        if p.poll() is not None and p.poll() !=0:
            error_counter+=1
    if error_counter>0:
        delete_files([sf+".bai" for sf in sortedfiles])
        sys.exit(1)

    print("##Variant Calling With GATK##")
    print("---------------------------------------------------------------------------------------------------------------------------------")
    print("")
    print("")

    #check if .dict file does exist, if not use picard to create it
    base=".".join(args.reference.split(".")[0:-1])
    dictfile="%s.dict"%(base)
    if not os.path.isfile(dictfile):
        cmd="java -jar /data/biosoftware/Picard/v2.9.0/picard.jar CreateSequenceDictionary R=%s O=%s"%(args.reference,dictfile)
        print("Creating dict file for reference")
        print(cmd)
        print("")
        os.system(cmd)

    #check if output directory for variant calling files exists, if not make it.
    gatkdir="%s/gatk_files"%(args.outputdirectory)
    if not os.path.isdir(gatkdir):
    	cmd="mkdir %s"%(gatkdir)
    	print("Making directory %s"%(gatkdir))
    	print(cmd)
    	print("")
    	os.system(cmd)

    print("##1. HaplotypeCaller##")
    print("----------------------------------------------------------------")
    print("")
    print("")

    vcffiles={}
    error_counter=0
    for sortedfile in sortedfiles:
        cmd,vcffile=haplotype_caller(args.reference,sortedfile,gatkdir)
        vcffiles[sortedfile]=vcffile
        runcmd=run_cmd_check(cmd,vcffile)
        inputfiles=[sortedfile]
        if len(input_absent(inputfiles))>0:
           print("missing: "+",".join(input_absent(inputfiles)))
           sys.exit(1)
        if runcmd:
            processes.add(subprocess.Popen(cmd.split()))
            if len(processes)>args.maxprocesses:
                os.wait()
                processes.difference_update([p for p in processes if p.poll() is not None])

    #check if all the processes were closed
    for p in processes:
    	if p.poll() is None:
    		p.wait()
    for p in processes:
        if p.poll() is not None and p.poll() !=0:
            error_counter+=1
    if error_counter>0:
        delete_files(list(vcffiles.values()))
        sys.exit(1)

    print("##2. SelectVariants##")
    print("----------------------------------------------------------------")
    print("")
    print("")

    filteredvcffiles={}
    error_counter=0
    for sortedfile in sortedfiles:
        vcffile=vcffiles[sortedfile]
        cmd,filteredvcf=select_variants(args.reference,vcffile)
        filteredvcffiles[sortedfile]=filteredvcf
        runcmd=run_cmd_check(cmd,filteredvcf)
        inputfiles=[vcffile]
        if len(input_absent(inputfiles))>0:
           print("missing: "+",".join(input_absent(inputfiles)))
           sys.exit(1)
        if runcmd:
            cmd=cmd.split()
            cmd.append('DP > 30.0')
            processes.add(subprocess.Popen(cmd))
            if len(processes)>args.maxprocesses:
                os.wait()
                processes.difference_update([p for p in processes if p.poll() is not None])

    #check if all the processes were closed
    for p in processes:
    	if p.poll() is None:
    		p.wait()
    for p in processes:
        if p.poll() is not None and p.poll() !=0:
            error_counter+=1
    if error_counter>0:
        delete_files(list(filteredvcffiles.values()))
        sys.exit(1)

    print("##3. BaseRecalibrator##")
    print("----------------------------------------------------------------")
    print("")
    print("")

    reportfiles={}
    error_counter=0
    for sortedfile in sortedfiles:
        filteredvcf=filteredvcffiles[sortedfile]
        cmd,reportfile=base_recalibrator(args.reference,filteredvcf,sortedfile,gatkdir)
        reportfiles[sortedfile]=reportfile
        runcmd=run_cmd_check(cmd,reportfile)
        inputfiles=[filteredvcf]
        if len(input_absent(inputfiles))>0:
           print("missing: "+",".join(input_absent(inputfiles)))
           sys.exit(1)
        if runcmd:
            processes.add(subprocess.Popen(cmd.split()))
            if len(processes)>args.maxprocesses:
                os.wait()
                processes.difference_update([p for p in processes if p.poll() is not None])

    #check if all the processes were closed
    for p in processes:
    	if p.poll() is None:
    		p.wait()
    for p in processes:
        if p.poll() is not None and p.poll() !=0:
            error_counter+=1
    if error_counter>0:
        delete_files(list(reportfiles.values()))
        sys.exit(1)

    print("##4. PrintReads##")
    print("----------------------------------------------------------------")
    print("")
    print("")

    recallbamfiles={}
    error_counter=0
    for sortedfile in sortedfiles:
        reportfile=reportfiles[sortedfile]
        cmd,recallbam=print_reads(args.reference,sortedfile,reportfile,gatkdir)
        recallbamfiles[sortedfile]=recallbam
        runcmd=run_cmd_check(cmd,recallbam)
        inputfiles=[reportfile]
        if len(input_absent(inputfiles))>0:
           print("missing: "+",".join(input_absent(inputfiles)))
           sys.exit(1)
        if runcmd:
            processes.add(subprocess.Popen(cmd.split()))
            if len(processes)>args.maxprocesses:
                os.wait()
                processes.difference_update([p for p in processes if p.poll() is not None])

    #check if all the processes were closed
    for p in processes:
    	if p.poll() is None:
    		p.wait()
    for p in processes:
        if p.poll() is not None and p.poll() !=0:
            error_counter+=1
    if error_counter>0:
        delete_files(list(recallbamfiles.values()))
        sys.exit(1)

    print("##5. HaplotypeCaller On Recalled Reads##")
    print("----------------------------------------------------------------")
    print("")
    print("")

    finalvcffiles=[]
    error_counter=0
    for sortedfile in sortedfiles:
        recallbam=recallbamfiles[sortedfile]
        cmd,finalvcf=haplotype_caller(args.reference,recallbam,gatkdir)
        finalvcffiles.append(finalvcf)
        runcmd=run_cmd_check(cmd,finalvcf)
        inputfiles=[recallbam]
        if len(input_absent(inputfiles))>0:
           print("missing: "+",".join(input_absent(inputfiles)))
           sys.exit(1)
        if runcmd:
            processes.add(subprocess.Popen(cmd.split()))
            if len(processes)>args.maxprocesses:
                os.wait()
                processes.difference_update([p for p in processes if p.poll() is not None])

    #check if all the processes were closed
    for p in processes:
    	if p.poll() is None:
    		p.wait()
    for p in processes:
        if p.poll() is not None and p.poll() !=0:
            error_counter+=1
    if error_counter>0:
        delete_files(finalvcffiles)
        sys.exit(1)

    print("")
    print("")

    print("##Merging VCF files##")
    print("---------------------------------------------------------------------------------------------------------------------------------")
    print("")
    print("")

    mergedvcffile="%s/%s"%(gatkdir,args.mergedvcffile)
    cmd="java -jar /data/biosoftware/GATK/GATK/GenomeAnalysisTK.jar -T GenotypeGVCFs -R %s --variant %s -o %s"%(args.reference," --variant ".join(finalvcffiles),mergedvcffile)
    if not os.path.isfile(mergedvcffile):
        print(cmd)
        print("")
        os.system(cmd)
    else:
        print("%s exists"%(mergedvcffile))
        print("")
        print("")


    print("##Obtaining Informative Variants##")
    print("---------------------------------------------------------------------------------------------------------------------------------")
    print("")
    print("")

    #check if output directory for informative variants analysis exists, if not make it.
    analysisdir="%s/informative_variants"%(args.outputdirectory)
    if not os.path.isdir(analysisdir):
    	cmd="mkdir %s"%(analysisdir)
    	print("Making directory %s"%(analysisdir))
    	print(cmd)
    	print("")
    	os.system(cmd)

    inputfiles=[mergedvcffile]
    if len(input_absent(inputfiles))>0:
        print("missing: "+",".join(input_absent(inputfiles)))
        sys.exit(1)
    splitvcffile=mergedvcffile.split('/')[-1].split('.')
    vcfbase=".".join(splitvcffile[:-1])

    if args.crosstype=="Backcross":
        informativevariantsfile="%s/%s_%s_%s_het_%s_hom.vcf"%(analysisdir,vcfbase,args.crosstype,args.heterozygousparent,args.homozygousparent)
        grandparents=args.grandparentstrain1+","+args.grandparentstrain2
        cmd="python informative_variants_QTLpipeline.py -f %s -d %s -c %s -t %s -m %s -g %s"%(mergedvcffile,analysisdir,args.crosstype,args.heterozygousparent,args.homozygousparent,grandparents)
    elif args.crosstype=="F1cross":
        grandparents=args.grandparents.split(",")
        informativevariantsfile="%s/%s_%s_grandparents_%s.vcf"%(analysisdir,vcfbase,args.crosstype,"_".join(grandparents))
        if args.parents is not None:
            cmd="python informative_variants_QTLpipeline.py -f %s -d %s -c %s -p %s -g %s"%(mergedvcffile,analysisdir,args.crosstype,args.parents,args.grandparents)
        else:
            cmd="python informative_variants_QTLpipeline.py -f %s -d %s -c %s -g %s"%(mergedvcffile,analysisdir,args.crosstype,args.grandparents)
    else:
        print("Cross type: %s is not recognised"%args.crosstype)
        sys.exit()

    if not os.path.isfile(informativevariantsfile):
        print(cmd)
        print("")
        os.system(cmd)
    else:
        print("%s exists"%(informativevariantsfile))
        print("")

    print("##Filtering Variants, Visualising Genotypes in a Heatmap, Determining Positions of Recombination and Creating a QTL Input Table##")
    print("---------------------------------------------------------------------------------------------------------------------------------")
    print("")
    print("")

    ##This pipeline runs genotype visualisation and recombination determnination with no filtering
    depthfilter=0
    fractionindividuals=0
    mendelianfraction=0
    maxdiffmissing=1
    mergedistance=0
    maxrecombinationindividuals=len(sampleids)
    minmarkersforrecombinationregion=0
    exclude=""

    #unfiltered
    inputfiles=[informativevariantsfile]
    if len(input_absent(inputfiles))>0:
        print("missing: "+",".join(input_absent(inputfiles)))
        sys.exit(1)
    informativevariantsbase=".".join(informativevariantsfile.split('.')[:-1])
    QTLinputfile=informativevariantsbase+"_minDP%s_%sfraction_mend%s_diffmissing%s_distmerge%s_maxrecomb%s_minmarkers%s_exclude%sindividuals%s_QTLinput.csv"%(depthfilter,fractionindividuals,mendelianfraction,maxdiffmissing,mergedistance,maxrecombinationindividuals,minmarkersforrecombinationregion,len(set(exclude)),"-".join(sorted(set(exclude))))

    if args.crosstype=="Backcross":
        cmd="python genotype_visualisation_and_classify_recombination_BACKcross_QTLpipeline.py --vcffile %s --depthfilter %s --fractionindividuals %s --mendelianfraction %s --maxdiffmissing %s --mergedistance %s --maxrecombinationindividuals %s --minmarkersforrecombinationregion %s"%(informativevariantsfile,depthfilter,fractionindividuals,mendelianfraction,maxdiffmissing,mergedistance,maxrecombinationindividuals,minmarkersforrecombinationregion)
    elif args.crosstype=="F1cross":
        if args.parents is None:
            numberofparents=0
        else:
            numberofparents=len(args.parents.split(","))
        cmd="python genotype_visualisation_and_classify_recombination_F1cross_QTLpipeline.py --vcffile %s --depthfilter %s --fractionindividuals %s --mendelianfraction %s --maxdiffmissing %s --mergedistance %s --maxrecombinationindividuals %s --minmarkersforrecombinationregion %s --numberofparents %s"%(informativevariantsfile,depthfilter,fractionindividuals,mendelianfraction,maxdiffmissing,mergedistance,maxrecombinationindividuals,minmarkersforrecombinationregion,numberofparents)

    if not os.path.isfile(QTLinputfile):
        print(cmd)
        print("")
        os.system(cmd)
    else:
        print("%s exists"%(QTLinputfile))
        print("")
    f1=open("%s/QTLinput_command.txt"%args.outputdirectory,"w+")
    print("module load python/2.7.13",file=f1)
    print(cmd,file=f1)
    f1.close()
```
