## Supplemental Script 2 for "Genetic analysis of a phenotypic loss in the mechanosensory entrainment of a circalunar clock": README.md

# QTL input pipeline
A Pipeline to prepare Restriction site-associated DNA sequencing (RAD-seq) data for quantitative trait locus (QTL) analysis.

Raw fastq files from a mapping family are processed into a genotype matrix that can be imported into R/qtl.

Please refer to the included [PDF](docs/QTL_Input_Pipeline_Documentation.pdf) for full documentation of function, description of parameters and the input and output files.

## Requirements
* python > 2.7
* pandas
* numpy
* matplotlib
* seaborn

## Usage
Example command:
`python QTL_input_pipeline.py -i INPUT_TABLE.tsv -g ADAPTERS.fa -r REFERENCE.fa -j 0.9 -s 0.9 -m THREADS -c CROSSTYPE -z PARENTS_LIST -p GRANDPARENTS_LIST -o OUTPUT_DIR -v OUTPUT.vcf`

Please refer to `python QTL_input_pipeline.py --help` for a complete list of options.
