## Supplemental Script 3 for "Genetic analysis of a phenotypic loss in the mechanosensory entrainment of a circalunar clock": Script_3_EM_scanone_README.txt

Take input files from combined geno+pheno and deposit them on the cluster.

Use <EM_SCRIPTS.txt> to generate 1000 R scripts, 1000 bash scripts to submit R scripts and a loop to submit the scripts. 

########################### RESULTS #################################################

1) load 1000 output panels in R on your PC: uniquepanels.systematic_1loops_10000iterations <1-1000> .csv
generate < all_pannels > matrix containing 1000 found phenotypes

2) Find frequency of all found panels: 

library(qtl)
library(mgcv)
library(plyr)

uniquepanels.systematic <- uniquecombs(all_pannels)
panellist <- attributes(uniquepanels.systematic)
freq.uniquepanels.systematic <- count(panellist\$index)
uniquepanels.systematic <- cbind(uniquepanels.systematic,freq.uniquepanels.systematic\$freq)

3) Calculate pentalties for unique panels <EM_penalties_1.xlsx.>

4) Use R/qtl to run scanone and record QTL intervals for all phenotype panels found in more than 5% out of 1000 runs.
