## Supplemental Script 4 for "Genetic analysis of a phenotypic loss in the mechanosensory entrainment of a circalunar clock": README.md

# vcf_update_cluma
Script _update_vcf.py_ converts a VCF file's genomic coordinates from reference A to manually edited reference B. 

As input, in addition to the VCF file, the script uses a GFF file containing all manual reference sequence edits. The GFF file is exported from WebApollo (version 2.5.0). Please see file _reference_manual_edits_29102020.gff3_ for all manual edits made to CLUMA2.0, to produce CLUMA2.0_M.

usage: 
`python update_vcf.py <vcf_file> <gff_file_with_reference_edits>`
