## Supplemental Script 4 for "Genetic analysis of a phenotypic loss in the mechanosensory entrainment of a circalunar clock": update_vcf.py

```
import sys
import numpy as np

if len(sys.argv)<3:
    print("usage: python update_vcf.py <vcf_file> <gff_file_with_reference_edits>")
    print("Converts a VCF file's genomic coordinates from reference A to manually edited reference B.")
    print("Differences (insertions and deletions) between ref A and ref B should be supplied in the GFF file.")
    exit()

annotations_gff=sys.argv[1]
withoutpath=annotations_gff.split("/")[-1]
outfile=".".join(withoutpath.split(".")[:-1])+"_positions_updated."+withoutpath.split(".")[-1]
reference_edit_gff_files=sys.argv[2]
new_positions_dict={}
withoutpath=reference_edit_gff_files.split("/")[-1]
editfile=".".join(withoutpath.split(".")[:-1])+"_list_of_edits.txt"
editout=open(editfile,"w+")
outstr="Chromosome\tPosition\tType\tLength\tString\n"
editout.write(outstr)
linecounter=0
for line in open(reference_edit_gff_files):
    if line.startswith("##sequence-region"):
        print(line)
        linesplit=line.split()
        chrom=linesplit[1]
        new_positions_dict[chrom]=np.arange(int(linesplit[2]),int(linesplit[3])+1).tolist()
        print("new positions list created")
    elif not line.startswith("#"):
        linecounter+=1
        if linecounter%10==0:
            print(linecounter,"edits")
        linesplit=line.strip().split("\t")
        chrom=linesplit[0]
        position=int(linesplit[3])
        type=linesplit[2]
        info=linesplit[8].split(";")[1]
        if info.startswith("residues"):
            editlength=len(info.split("=")[1])
        else:
            print("'residues' not found")
        editout.write("%s\t%d\t%s\t%d\t%s\n"%(chrom,position,type,editlength,info.split("=")[1]))
        position_index=position-1
        if type=="insertion_artifact":
            for i in range(position_index+1,len(new_positions_dict[chrom])):
                new_positions_dict[chrom][i]+=editlength
        elif type=="deletion_artifact":
            for i in range(1,editlength):
                new_positions_dict[chrom][position_index+i]-=i
            for i in range(position_index+editlength,len(new_positions_dict[chrom])):
                new_positions_dict[chrom][i]-=editlength
editout.close()
out=open(outfile,"w+")
linecounter=0
for line in open(annotations_gff):
    if line.startswith("#"):
        out.write(line)
    else:
        linecounter+=1
        if linecounter%100==0:
            print(linecounter,"updated")
        linesplit=line.split("\t")
        chrom=linesplit[0]
        start=int(linesplit[1])
        position_index=start-1
        newstart=new_positions_dict[chrom][position_index]
        linesplit[1]=str(newstart)
        lineout="\t".join(linesplit)
        out.write(lineout)
out.close()
```
